# Behavioral differences among domestic cats in the response to cat-attracting plants and their volatile compounds reveal a potential distinct mechanism of action for actinidine

**DOI:** 10.1101/2022.03.05.483118

**Authors:** Sebastiaan Bol, Adrian Scaffidi, Evelien M. Bunnik, Gavin R. Flematti

## Abstract

It has been known for centuries that cats respond euphorically to *Nepeta cataria* (catnip). Recently, we have shown that *Lonicera tatarica* (Tatarian honeysuckle), *Actinidia polygama* (silver vine) and *Valeriana officinalis* (valerian) can also elicit this “catnip response”. The aim of this study was to learn if the behavior seen in response to these plants is similar to the response to catnip. Furthermore, we studied if these responses are fixed or if there are differences between cats. While nepetalactone was identified decades ago as the molecule responsible for the “catnip response”, we know that this volatile is found almost exclusively in catnip. Therefore, we also aimed to identify other compounds in these alternative plants that can elicit the blissful behavior in cats. Bioassays with 6 cats were performed in a stress-free environment, where 6 plants and 13 single compounds were each tested for at least 100 and 17 hours, respectively. All responses were video recorded and BORIS software was used to analyze the cats’ behavior. Both response duration and behavior differed significantly between the cats. While individual cats had preferences for particular plants, the behavior of individual cats was consistent among all plants. About half a dozen lactones similar in structure to nepetalactone were able to elicit the “catnip response”, as were the structurally more distinct molecules actinidine and dihydroactinidiolide. Most cats did not respond to actinidine, whereas those who did, responded longer to this volatile than any of the other secondary plant metabolites, and different behavior was observed. Interestingly, dihydroactinidiolide was also found in excretions and secretions of the red fox, making this the first report of a compound produced by a mammal, that can elicit the “catnip response”. A range of different cat-attracting compounds was detected by chemical analysis of plant materials but differences in cat behavior could not be directly related to differences in chemical composition of the plants. Together with among other results of habituation / dishabituation experiments, this indicates that additional cat-attracting compounds may be present in the plant materials that remain to be discovered. Collectively, these findings suggest that both the personality of the cat and genetic variation in the genes encoding olfactory receptors may play a role in how cats respond to cat-attracting plants. Furthermore, the data suggest a potential distinct mechanism of action for actinidine.

## Introduction

Cats are lured by the volatiles of several plant species and unlike any other animal they demonstrate what appears to be blissful behavior in response to smelling them. Of these plants, the species *Nepeta cataria* (catnip) and *Actinidia polygama* (silver vine) are the best-known to elicit such a response. The former is commonly used by cat caregivers in Europe and North America, while the latter is more popular in Asia, where it is also known as matatabi. After sniffing these plants, head rubbing and rolling over are typically observed, and this behavior is generally referred to as the “catnip response” (Bol et al. 2017, Todd 1963). While the joyful effects of some plants from the genus *Nepeta* on cats has been known to humans for centuries (Ray 1660, Salmon 1710), it is still unclear if there is a biological reason for the response of cats to this select group of plants. Felines are believed not to be the intended recipients of the allomones produced by these plants. This unique response of cats appears to be fortuitous, since plants produce these secondary metabolites to protect themselves against phytophagous or parasitic insects. The cat-attracting compounds synthesized by a small number of species within the plant kingdom are identical or closely related to insect pheromones or allomones (Beran et al. 2019, Eisner 1964).

Insects release these chemicals when in danger (Ho and Chow 1993, Kanehisa, Tsumuki and Kawazu 1994) and for this reason it is assumed plants produce and release these chemicals to send a warning message to phytophagous insects (Ebrahim et al. 2015, Stökl et al. 2012, Welzel et al. 2018). Recently, Nadia Melo and her colleagues revealed the molecular mechanism by which the iridoid nepetalactone repels insects (Melo et al. 2021).

Nepetalactone, found in *Nepeta cataria*, was the first compound identified as being able to elicit the catnip response (McElvain, Bright and Johnson 1941). Several other compounds similar in structure, have been reported to have effects comparable to nepetalactone (Sakan et al. 1959, Sakan, Fujino and Murai 1960, Sakan et al. 1965, Johnson and Waller 1971, Scaffidi et al. 2016), but bioassays with cats were not performed. However, behavior analogous to the “catnip response” was observed when felines were exposed to *A. polygama*, *Lonicera tatarica* (Tatarian honeysuckle) and *Valeriana officinalis* (valerian) root, all containing little to no nepetalactone (Bol et al. 2017). Those results suggest other compounds are also able to elicit the “catnip response”. Unpublished work (doctoral dissertation) by Nelson and Wolinsky done more than 50 years ago provided some more insight into which compounds might be able to elicit the “catnip response” in domestic cats, which included several lactones (nepetalactone, epinepetalactone, iridomyrmecin, isoiridomyrmecin, dihydronepetalactone, isodihydronepetalactone, neonepetalactone) and matatabiether (Nelson 1968). Results from a recent study by Reiko Uenoyama *et al*., that were published while this manuscript was in preparation, indicated that domestic cats respond to a variety of lactones (nepetalactone, iridomyrmecin, isoiridomyrmecin, dihydronepetalactone, isodihydronepetalactone) as well as nepetalactol (Uenoyama et al. 2021). Most of what is known about the behavior of domestic cats seen in response to cat-attracting plants originates from a limited number of studies where only catnip was used (Todd 1962, Todd 1963, Hill et al. 1976, Palen and Goddard 1966). With this study we tried to answer several questions, including the following. (**i**) We wanted to know if the cats’ behavior to other known cat-attracting plants is the same as to catnip. To this end, we performed comprehensive behavioral analysis of 6 domestic cats in response to *Actinidia polygama* (silver vine), *Lonicera tatarica* (Tatarian honeysuckle), *Valeriana officinalis* (valerian) and the arcane *Acalypha indica* (Indian nettle), and compared these responses to the behavior seen in response to *Nepeta cataria* (catnip). (**ii**) In addition, we wanted to learn if the “catnip response” is a fixed, predictable, biological response to these cat-attracting plants, or if there is variation in the response between cats. Therefore, we also compared the observed behavior between the 6 cats. (**iii**) Furthermore, we wanted to know which single compounds the cats respond to and understand which features of these molecules are responsible for the response. For this reason, we studied the response of domestic cats to all lactones tested by Uenoyama *et al*., but also included indole, neonepetalactone, isoneonepetalactone, and the structurally more distinct actinidine (a pyridine) and dihydroactinidiolide (a furanone), both known to be present in *A. polygama* (Sakan et al. 1965, Sakan, Isoe and Hyeon 1967, Sakan et al. 1969, Bol et al. 2017). Not only did we test if cats responded to these compounds from different classes, but (**iv**) we were also interested to see if the cats’ behavior varies between the different compounds or between cats. After video recording the responses of 6 domestic cats to 5 different plants and 13 single compounds on 72 days between the summer of 2018 and the winter of 2020, we analyzed 470 responses to plants, totaling over 8 hours of response time, and 217 responses to single compounds, totaling over 2.5 hours of response time. Of these, the behavior of 179 responses (88 to plants and 91 to single compounds), totaling over 77 and 80 minutes, respectively, were analyzed in detail using behavioral analysis software. In addition to the behavioral studies, (**v**) we quantified the amount of the various single compounds in the plants that were used in this study in an attempt to correlate these with the duration and behavior seen in response to the plants.

## Materials and methods

### Ethics

No cats were hurt or distressed for this study, nor were they ever forced to act, respond, or behave in any way. All research with cats was non-invasive and did not involve pharmacological, medical, or surgical intervention. All participating cats were adopted and are permanent residents of Cowboy Cat Ranch, living together with authors SB and EMB. The study protocol was approved by the Cowboy Cat Ranch Institutional Animal Care and Use Committee (IACUC). Cowboy Cat Ranch was registered with the USDA as a research facility (registration number 74-R-0224, ID no. 502147) during the time of the study (2018 – 2020). The research facility and the cats were inspected annually by the Cowboy Cat Ranch IACUC and the USDA Animal and Plant Health Inspection Service.

### Study population

Six healthy, neutered, adult, domestic short-haired cats (**Table 1**) participated in this study that commenced in June 2018 and ended in December 2020. All cats were adopted from a local shelter, with cats N, O and V from the same litter. In December of 2018 cat H needed to be separated from the other cats for medical reasons and was therefore not exposed to *V. officinalis* and the single compounds. All cats were seen by a veterinarian (Babcock Hills Veterinary Hospital in San Antonio, TX, USA) for routine veterinary care (physical examination, blood tests, vaccinations, dental cleaning and dental X-rays) at least once a year, were treated once a month with Catego (dinotefuran, fipronil and pyriproxyfen) for flea and tick control, and received milbemycin oxime for heartworm prevention and intestinal parasite control once a month. In November 2020 cat A was diagnosed with hyperthyroidism and treated with radioiodine that same month.

**Table 1.** Age, breed, hair-color and pattern, and gender of the cats who participated in the study.

| Name (abbreviation) | Age<sup>1</sup> | Breed | Color / pattern | Gender |
| --- | --- | --- | --- | --- |
| Aguereberry (A) | 11Y 1M | domestic short-haired | calico | female |
| Harvey (H) | 1Y 4M | domestic short-haired | orange | female |
| Namibia (N) | 3Y 5M | domestic short-haired | grey tabby | female |
| Olli (O) | 3Y 5M | domestic short-haired | black | male |
| Vlinder (V) | 3Y 5M | domestic short-haired | grey tabby | female |
| Zappa (Z) | 6Y 6M | domestic short-haired | tortoiseshell | female |

<sup>1</sup> Age in years (Y) and months (M) at the start of the study

### Study environment

All experiments were done in a stress-free setting. Cowboy Cat Ranch is the permanent home of all the cats who participated in this study, as well as the authors and researchers SB and EMB. It is a one-story house with 195 m^2^ indoor living space and 51 m^2^ enclosed outdoor space on 4.1-hectare privately-owned land, with little to no distraction. The testing area consisted of a 6.3 m^2^ (3.3 × 1.9 m) piece of vinyl sheet (**Supplementary Figure 1A**) placed in the center of the floor of a 45 m^2^ (8.2 × 5.5 m^2^) open room that was recorded continuously when an olfactory test sample was made available to the cats. The cats were motivated to spend time in the open room with the testing area by temporarily restricting their living space (not allowing access to other rooms or outdoor enclosures), offering treats close to, but not at, the testing area at set times, or by being present in the room where the olfactory samples were available. The latter two strategies were only employed when the single compounds were tested. For the safety of the cats, the IACUC required at least one human to be present during the testing of these compounds to actively look for any potential adverse reactions (none were observed), since felines have not been previously exposed to several of these before. When the tested plants were available for the cats (10 days per plant, 10 hours per day), no humans were present. The minimum living space for the cats during the testing period was 120 m^2^ and contained twelve large (about 2 m tall) cat trees, more than 37 m of wall shelves, multiple cat beds and comforters to provide vertical space and hiding places. None of the cats were ever forced to be at a certain location, act, respond or behave in any way. The indoor temperature was maintained constant at 20 – 23°C and all rooms were illuminated when it was dark outside. The cats were fed canned food four to six times a day, had continuous access to running and standing water, multiple litter boxes and fresh (no older than 14 days after seeding) oat grass.

### Plants

Five different plant species were used in this study: *Nepeta cataria* or catnip, *Acalypha indica* or Indian nettle, *Actinidia polygama* or silver vine, *Lonicera tatarica* or Tatarian honeysuckle, and *Valeriana officinalis* or valerian (**Table 2, Supplementary Figure 1B-G**). *Actinidia polygama* ‘Hot Pepper’ (female) and ‘Pavel’ (male) varieties were purchased as one gallon-size plants in 2017. Before collecting leaves and woody stems in October/November 2020, the plants had been growing in Mico, Texas, USA for three and a half years. Leaves and stems from these plants were dried at room temperature and 30-50% humidity for one to two weeks. The stem used for testing was woody, 15 cm long, and had a diameter of 1 cm (similar to what is commercially available). With the exception of *A. polygama* stem, 15 g of each plant material was offered to the cats. *A. indica* roots were collected from Christmas Island, Australia as described previously (Scaffidi et al. 2016). The roots were washed free of soil material and lyophilized immediately after collection and stored in vacuum sealed bags until use. All plant materials were stored airtight at room temperature, away from direct sunlight.

**Table 2.** An overview of the plant materials that were used in this study.

| Plant species (common name) | Tissue | Source / brand |
| --- | --- | --- |
| *Acalypha indica* (Indian nettle) | roots (lyophilized, cut) | Christmas Island, Government of Western Australia |
| *Actinidia polygama* (silver vine) | fruit galls (dried, powder) | Smack (smack.co.jp) |
| *Actinidia polygama* varieties 'Hot Pepper' (female) and 'Pavel' (male) | leaves (dried, cut) and stem (lignified, dried) | One Green World (Portland, Oregon, USA) |
| *Lonicera tatarica* (Tatarian honeysuckle) | wood (sawdust) | The Cat House (Calgary, Alberta, Canada) |
| *Nepeta cataria* (catnip) | leaves (dried, cut) | Frontier /SmartyKat |
| *Valeriana officinalis* (valerian) | roots (dried, cut) | Frontier |
| *Camellia sinensis* (green tea)<sup>1</sup> | leaves (dried, cut) | Frontier |

<sup>1</sup> Used as negative control

### Single compounds

Thirteen different single compounds were used in this study to test if and how domestic cats responded to them (**Table 3**): (**1**) nepetalactone ((4a*S*,7*S*,7a*R*)-4,7-dimethyl-5,6,7,7a-tetrahydrocyclopenta[c]pyran- 1(4a*H*)-one), (**2**) epinepetalactone ((4a*S*,7*S*,7a*S*)-4,7-dimethyl-5,6,7,7a-tetrahydrocyclopenta[c]pyran- 1(4a*H*)-one), (**3**) dihydronepetalactone ((4*S*,4a*R*,7*S*,7a*R*)-4,7-dimethylhexahydrocyclopenta[c]pyran- 1(3*H*)-one), (**4**) isodihydronepetalactone ((4*R*,4a*R*,7*S*,7a*R*)-4,7-dimethylhexahydrocyclopenta[c]pyran- 1(3*H*)-one), (**5**) neonepetalactone ((4*S*,4a*R*)-4,4a,5,6-tetrahydro-4,7-dimethylcyclopenta[c]pyran-1(3*H*)- one), (**6**) isoneonepetalactone ((4*R*,4a*R*)-4,4a,5,6-tetrahydro-4,7-dimethylcyclopenta[c]pyran-1(3*H*)-one), (**7**) iridomyrmecin ((4*S*,4a*S*,7*S*,7a*R*)-4,7-dimethylhexahydrocyclopenta[c]pyran-3(1*H*)-one), (**8**) isoiridomyrmecin ((4*R*,4a*S*,7*S*,7a*R*)-4,7-dimethylhexahydrocyclopenta[c]pyran-3(1*H*)-one), (**9**) actinidine ((*S*)-4,7-dimethyl-6,7-dihydro-5*H*-cyclopenta[c]pyridine), (**10**) dihydroactinidiolide ((*R*)-4,4,7a-trimethyl- 5,6,7,7a-tetrahydrobenzofuran-2(4*H*)-one), (**11**) indole, menthol, and methyl salicylate. Dihydroactinidiolide (**10**) and indole (**11**) were purchased from AK Scientific (#J10744 and #I908, respectively). Compounds **1** and **9** (Beckett, Beckett and Hofferberth 2010), **2** – **4** (Scaffidi et al. 2016), **5** – **6** (Enders and Kaiser 1997) and **7** – **8** (Stepanov and Veselovsky 1997) were all synthesized according to literature procedures and shipped from Australia to the USA at room temperature in two clear glass vials with Teflon screw caps, each containing approximately 5 mg of material accurately weighed. Upon arrival, the vials were stored in the dark at 4°C. Immediately prior to testing the compounds were dissolved in diethyl ether (Acros #448421000) at a concentration of 1 mg/mL using a Gilson Microman E M1000E positive displacement pipet with 1 mL capillary pistons. The concentration of 1 mg/mL is equivalent to about 6 mM for the lactones, 7 mM for actinidine, 5.5 mM for dihydroactinidiolide, 6.5 mM for menthol and methyl salicylate, 4 mM for civetone and 8.5 mM for indole. While we did not normalize based on the molecular weight of the compounds, three different amounts of each compound were tested: 33 µg, 100 µg, 300 µg and 900 µg, equivalent to 33, 100, 300 and 900 µL of solution. These amounts were chosen in the absence of information about the lower level of detection in domestic cats and were somewhat arbitrary. While Nelson and Wolinsky tested 3,000 – 5,000 µg (Nelson 1968), we decided to start with the lowest amount that we could reliably pipette and increase the volume (and hence the amount of the compound) during the day or subsequent testing on another day to rule out that cats did not respond because the amount used was too low. 50 – 400 µg of iridoids was used in the study led by Masao Miyazaki (Uenoyama et al. 2021), who published the results while this manuscript was in preparation. We did not screen different amounts with the intent to establish a dose response relationship, since this would be complicated by both fluctuating activity levels of domestic cats during the day (typically less active in the afternoon). Furthermore, previous exposures may have an effect on the subsequent testing of higher amounts of the compounds, especially when different amounts are tested on the same day. Since there was no information available on the stability of these compounds when dissolved in diethyl ether, the goal was to test them immediately. This limited us to two testing days.

**Table 3.** An overview of the single compounds used in this study.

| # |  | Compound | Class<sup>1</sup> | Retention index<sup>2</sup> | Source |
| --- | --- | --- | --- | --- | --- |
| 1 | A<sup>3</sup> | nepetalactone<br>(*cis-trans*-nepetalactone) | type II lactone | 1383 | synthesized |
| 2 | A | epinepetalactone<br>(*trans-cis*-nepetalactone) | type II lactone | 1416 | synthesized |
| 3 | B | dihydronepetalactone | type II lactone | 1490 | synthesized |
| 4 | B | isodihydronepetalactone | type II lactone | 1446 | synthesized |
|  | B | *trans*-dihydronepetalactone<sup>4</sup> | type II lactone | 1505 | synthesized |
|  | B | *trans*-isodihydronepetalactone<sup>4</sup> | type II lactone | 1470 | synthesized |
| 5 | C | neonepetalactone | type II lactone | 1517 | synthesized |
| 6 | C | isoneonepetalactone | type II lactone | 1511 | synthesized |
| 7 | D | iridomyrmecin | type I lactone | 1466 | synthesized |
| 8 | D | isoiridomyrmecin | type I lactone | 1478 | synthesized |
| 9 |  | actinidine | pyridine | 1348 | synthesized |
| 10 |  | dihydroactinidiolide | furanone | 1562 | AK Scientific |
| 11 |  | indole |  |  | AK Scientific |
| 12 |  | menthol |  |  | GreenHealth |
| 13 |  | methyl salicylate<sup>5</sup> |  |  | TCI Chemicals |

<sup>1</sup> The difference between type I and II lactones is the position of the carbonyl group (Nangia, Prasuna and Bheema Rao 1997).
<sup>2</sup> Linear retention index relative to n-alkanes on a DB-5ms column
<sup>3</sup> The same letters in the second column of this table indicates these compounds are diastereoisomers: stereoisomers with one or more differing stereocenters resulting in different molecules that are not mirror images and not superimposable.
<sup>4</sup> These compounds were only prepared in small amounts and used as standards in the GC-MS analysis, but were not used in bioassays with cats.
<sup>5</sup> Liquid at room temperature

However, a limited amount of the compounds (neonepetalactone and isoneonepetalactone: one vial with 5 mg) and the need for additional testing (neonepetalactone: technical problems with recording; *trans*-*cis*- nepetalactone and actinidine: assuring that lack of response was not due to limited exposure time) required us to store compounds dissolved in diethyl ether for short periods of time. Neonepetalactone and isoneonepetalactone were stored in the dark at 4°C for 4 days before 100 µg, 300 µg and 900 µg were tested. Neonepetalactone had to be stored for a further 6 days because of technical issues while recording. Testing *trans*-*cis*-nepetalactone and actinidine on more than two days was not anticipated, but motivated by the results of the first two testing days (absence of response by multiple cats). After the second test day, *trans*-*cis*-nepetalactone dissolved in diethyl ether was stored for one and a half months at room temperature. Actinidine dissolved in diethyl ether was stored in the dark but at fluctuating temperatures, ranging from freezing to room temperature and was used once to test 900 µg after two weeks of storage. Actinidine used to test on the 4^th^ and 5^th^ day was a new shipment from Australia, which was re-purified (silica chromatography) prior to shipping because some degradation (browning) of the original stock (that was stored neat at 4⁰C) was noticed. In addition to the compounds mentioned above, a small amount of *trans*-dihydronepetalactone ((4*S*,4a*R*,7*S*,7a*S*)-4,7- dimethylhexahydrocyclopenta[c]pyran-1(3*H*)-one) and *trans*-isodihydronepetalactone ((4*R*,4a*R*,7*S*,7a*S*)-4,7-dimethylhexahydrocyclopenta[c]pyran-1(3*H*)-one) was used for quantitative analysis and prepared by reducing epinepetalactone (**2**) in a similar fashion to the preparation of dihydronepetalactone (**3**) and isodihydronepetalactone (**4**) from nepetalactone (**1**) (Scaffidi et al. 2016).

### Testing procedures

Cats were familiarized with the vinyl sheet and their altered environment for at least one month prior to the start of the study. Thin, fibrous, porous, polyester socks with 2 – 3% spandex were used as carriers of the olfactory material. The testing area accommodated a maximum of four socks / samples at the same time. Socks were mounted near each corner of the testing area using twine in a way that allowed for some movement (25 cm radius) of the sock, but eliminated cross-over contamination between samples or controls (**Supplementary Figure 1A**) and prevented cats from moving the socks outside of the 4.5 m^2^ area captured by the camera. The olfactory samples were deliberately offered for many hours and on multiple days to reduce the chance that a cat would not respond because of unawareness of the presence of the olfactory stimulus, competition for the sample with another cat, or hindrance in any other way by another cat. The vinyl sheet was cleaned with water and soap before and after every testing day. Responses where the cats displayed behavior listed in the ethogram (see the paragraph *behavioral analysis* below) were considered catnip-responses or positive responses.

#### Plants

Five different plants plus a negative control (green tea) were tested between June 2018 and May 2019 (**Table 2, Supplementary Figure 2A**). Each sock contained 15 g of plant material. Samples and negative controls were offered in duplicate to allow two cats to respond simultaneously. Samples were mounted diagonally across so responding cats had enough space and would not disturb each other. Plant materials were offered for 10 hours (between 9:30 and 19:30) on ten different days (Monday – Friday only), within a total time period of five weeks: five days in the first two weeks and five days in the last two weeks, separated by at least one full week (Monday – Sunday) of no exposure. Exposure was limited to no more than three days in one week (Monday – Friday), never more than two days of exposure in a row, but at least one consecutive two-day exposure per two-week period (**Supplementary Figure 2A**). To avoid human presence affecting the outcome of the experiments no humans were present during these tests. The schedule described above allowed for some flexibility, while keeping the testing conditions highly similar for all plants tested. Different plant samples were offered no sooner than one week after the previous testing was completed. The same socks and plant materials were used throughout the 5-week testing period. On days and nights when the olfactory stimuli were not tested, they were stored in an air- tight bag, away from direct sunlight, at room temperature. New socks were used when new plant materials were tested. Since the tested plant samples could easily be identified by their smell, the experiments with the plants were not performed blinded.

#### Single compounds

The single compounds were tested from December 2018 through to August 2020. Various volumes (33, 100, 300 or 900 µL) of compound dissolved in diethyl ether (final concentration of 1 µg/µL) were applied on the outside of a sock. Equal amounts of diethyl ether were used for the negative control socks. The diethyl ether was allowed to evaporate prior to mounting the socks. Compounds were tested on two different days: 33 and 100 µg on day A for 1 and 4 hours in the afternoon, respectively, and 100, 300 and 900 µg on day B, all for 4 hours, starting at 7:30, ending at 19:30. All samples to be tested on the same day (e.g., socks with 33 µL and 100 µL for day A) were prepared early in the morning, and were stored in an air-tight bag away from direct sunlight at room temperature prior to use. There were always 3 days between days A and B, to rule out responses on day A affecting responses on day B. While the recording area had a capacity for 4 samples, no more than two different compounds were tested at the same time. When possible, different combinations of compounds were tested on day A than on day B (e.g., *cis*-*trans*- nepetalactone and iridomyrmecin on day A, but on day B *cis*-*trans*-nepetalactone was tested together with dihydronepetalactone). We chose for this rotating setup to prevent false negative responses that were the result of a strong preference for one compound over the other. Testing of *trans*-*cis*- nepetalactone was repeated for this reason since this compound was offered in combination with actinidine on both days, and cat A was extremely attracted to actinidine. We aimed to test the single compounds as soon as possible after they were received as we were unsure about their stability.

However, not all the compounds were received at the same time and therefore some compounds were tested together on both days A and B. Each sample was always accompanied by a negative control. To comply with IACUC guidelines, at least one human was present during exposure tests with the single compounds, since at the time the tests were conducted, no safety information was available for these compounds. Socks containing higher amounts of the single compound were mounted on the same location to prevent cross-contamination of the vinyl surface area. All compounds were coded and the testing and analysis (response frequency and duration) were done blind.

### Behavioral analysis

All responses of the cats to the plants or single compounds were video recorded. Behavioral analyses of the responses were performed using the free, open-source software BORIS (version 7.9.19) (Friard and Gamba 2016). The ethogram shown in **Table 4** was used for the video coding of the cats’ behavior. The video ethogram shows the behavior listed in the ethogram of the domestic cats who participated in this study in response to the cat-attracting plants or single compounds (**Supplementary File 1**). Four recordings are shown for each behavior listed in Table 4. The analysis using BORIS software was not done blinded, since behavioral comparisons were done using recordings of responses with known duration and hence after the quantitative analysis. Body position, biting, head rubbing, holding, licking and raking were expressed as the percentage of the total response duration as determined by using BORIS. Head shaking, rippling of the back, rolling on the side and twitching of the back were reported as events per minute and plotted on a different Y axis to allow for better visualization of these behaviors and discrimination of their frequency between cats or stimuli. Small discrepancies exist between the total response time (used to calculate differences in response duration and timed with a stopwatch) and the sum of the duration of all behaviors scored in BORIS. For the former, the total time the cat was responding was used, and sometimes included aspects of the response that were not scored in BORIS (e.g., stretching out while in lateral position in the middle of the response, but not actively engaging with the test object, or, rarely seen, playfully running away from and towards the test object, swatting it when passing by). In rare situations it was difficult to determine with certainty what the cat was doing, for example, discriminating between head rubbing, licking or biting. When in doubt, the behavior was not scored in BORIS.

**Table 4:** Ethogram describing body positions and behaviors seen in domestic cats in response to cat-attracting plants or their volatile compounds.

| Body position | Description |
| --- | --- |
| standing | The cat is in an upright position with all paws on the ground and the legs extended. |
| sitting | The cat is sitting in a crouched position: the body is close to the ground, all legs are bent, and the belly is touching or raised slightly off of the ground; crouched down to get a closer look at the object, not to be mistaken with crouching because of fear. |
| lying on side | The cat lies on her or his left or right side. |
| lying on back | The cat lies on her or his back. |
| Behavior | Description |
| biting<sup>1</sup> | The cat bites the object or has the object in her or his mouth. Sometimes combined with pulling or shaking her or his head. |
| head rubbing<sup>1</sup> | The cat rubs with her chin, cheek or forehead against the object. |
| head shaking<sup>1</sup> | The cat shakes her or his head without an object in her or his mouth. Sometimes combined with shaking the rest of the body. |
| holding<sup>1</sup> | The cat holds an object with one or two paws. |
| licking | The cat passes her or his tongue over the object. |
| raking<sup>1</sup> | The cat makes kicking movements with one or both hind legs against the object. Also known as bunny kicking. Typically seen when the cat holds the object with her or his paws or in her or his mouth. |
| rippling of back<sup>1</sup> | Rippling or rolling motion of the cat's skin in the dorsal lumbosacral region as the underlying cutaneous trunci / panniculus carnosus muscles rhythmically contract and relax. Not to be confused with feline hyperesthesia syndrome. |
| rolling on side<sup>1</sup> | The cat rolls on her or his side or back, from a sternal or lateral body position, respectively. |
| twitching of back<sup>1</sup> | Short (fraction of a second), quick contractions of the cutaneous trunci / panniculus carnosus muscles. Distinct (shorter) from rippling of the back, but possibly related. |

<sup>1</sup> See **Supplementary File 1** for a video with examples of these behaviors.

### Habituation / dishabituation testing

*Actinidia polygama*, *Lonicera tatarica* and *Nepeta cataria* (Frontier) were used to study habituation / dishabituation of domestic cats to these cat-attracting plants (**Supplementary Figure 2B**). Half a gram and 2.5 g dried *A. polygama* fruit gall powder, 2.5 g *L. tatarica* sawdust and 2.5 g dried *N. cataria* leaves were available to the cats. Each plant material was tested on at least 10 consecutive days for either 2 or 12 hours (20:00 – 22:00 and 10:00 – 22:00, respectively). Two socks were available for each plant material. No negative controls were used in these experiments. Prior to testing, no olfactory stimuli were available to the cats for at least 2 weeks.

### Detection of Pseudasphondylia matatabi in A. polygama fruit galls

Forty milligrams of dried *A. polygama* fruit galls were powdered by grating, and subsequently used to isolate total DNA with Zymo’s Quick-DNA Microprep Plus kit (#D4074) as per manufacturer’s instructions. Potential PCR inhibitors were removed using Zymo’s OneStep PCR Inhibitor Removal kit (#D6030). *P. matatabi* mitochondrial cytochrome c oxidase subunit 1 (COX) (GenBank AB085873.1) DNA was amplified with AccuStart II PCR SuperMix and *P. matatabi* specific primers 5′– AGGAACTGGAACAGGATGAACA–3′ and 5′–AAAATTGGGTCTCCACCTCCT–3′ (250 nM final concentration) using the following program: 3 min. at 95°C, 35 × (30 sec. at 95°C, 30 sec. at 60°C, 30 sec. at 72°C), and 2 min. at 72°C. The 330 bp *cox1* amplicon was Sanger sequenced and BLASTn was used to identify the species.

### DNA barcoding of plants

DNA was isolated from 40 – 100 mg fresh leaves (*Actinidia* species) or 20 – 80 mg wood chips (*Lonicera* species) using Omega Bio-Tek’s E.Z.N.A. Plant DNA kit (#D2411-00) as per manufacturer’s instructions. *matK* was amplified with AccuStart II PCR SuperMix (QuantaBio #89235-018) and primers 5′– CGTACAGTACTTTTGTGTTTACGAG–3′ and 5′–ACCCAGTCCATCTGGAAATCTTGGTTC–3′ (250 nM final concentration) (Weihong, Dawei and Xinwei 2018) using the following program: 3 min. at 95°C, 40 × (30 sec. at 95°C, 40 sec. at 60°C, 60 sec. at 72°C), and 5 min. at 72°C. *rbcL* was amplified with AccuStart II and primers 5′–ATGTCACCACAAACAGAAAC–3′ and 5′–TCGCATGTACCTGCAGTAGC–3′ (Weihong et al. 2018) using the following program: 1 min. at 94°C, 35 × (10 sec. at 94°C, 20 sec. at 60°C, 45 sec. at 70°C). The *psbA – trnH* intergenic spaces was amplified using primers 5′– GTTATGCATGAACGTAATGCTC–3′ and 5′–CGCGCATGGTGGATTCACAATCC–3′ (Sun et al. 2011) as described for *rbcL*. After DNA cleanup using NEB’s Monarch PCR Cleanup kit (#T1030G) the amplicons of approximately 450 – 800 bp were Sanger sequenced using the forward and reverse primer. T-Coffee (Notredame, Higgins and Heringa 2000) was used to align the sequences and the consensus sequence was used to identified the species using NIH’s nucleotide BLAST.

### Tinctures

Tinctures were made by adding five volumes (500 mL) of absolute ethanol (Fisher Scientific, #BP2818) to approximately 100 mL volume of plant materials inside a glass bottle. This included: dried catnip leaves (Frontiers; 10 grams), Tatarian honeysuckle sawdust (20 grams) and dried valerian roots (50 grams). The bottles were closed with a screw cap and were stored at room temperature in the dark with daily mixing for 18 months. The liquid fraction was collected into a glass spray bottle by aspiration using a Pipet-Aid, without disturbing the plant material sediment. Two sprays of the tincture (about 200 µL) were applied to a fabric (empty polyester sock with 2 – 3% spandex), one on each side. All three tinctures and a control fabric were offered for 5 hours on one afternoon/evening in October of 2020. No other olfactory stimulant was offered to the cats at least two weeks prior to testing these tinctures.

### Fragrances

Fabrics (empty polyester sock with 2 – 3% spandex) containing either a fragrance (**Table 5**) or negative control (absolute ethanol) were offered to the cats on 8 different days between early September 2020 and mid-December 2020. No olfactory stimuli were offered at least five days prior to testing of the fragrances. Fabrics sprayed once on each side were made available to the cats immediately, whereas fabrics sprayed abundantly (about 10 sprays) were left to stand for 10 hours at room temperature prior to making them available to the cats (**Table 6**). Each sample was available to the cats for 15 hours (7:00 – 22:00).

**Table 5.** An overview of the fragrances used in this study.

| Company | Fragrance | Type or concentration | Place of purchase |
| --- | --- | --- | --- |
| Calvin Klein | Obsession for Men | eau de toilette | (1) Nordstrom, The Shops at La Cantera, San Antonio, Texas, USA. (2) The Fragrance Decant Boutique (decantboutique.com), Little Elm, Texas, USA. (3) USA Fragrance, Ingram Park Mall, San Antonio, Texas, USA |
| Nina Ricci | L'Air Du Temps | eau de parfum | Amazon |
| Guy Laroche | Drakkar Noir | eau de toilette | iDimino |
| Paco Rabanne | Paco Rabanne Pour Homme | eau de toilette | Natural Nutrient |
| SP Parfums | Civette Intense | eau de parfum | Indie Scents |
| Matieres Premieres Essentielles | Civet Absolute (synthetic) | 1% in absolute ethanol | Perfumer's Apprentice |
| Firmenich | Civettone | 0.1 and 1% civetone in absolute ethanol | Firmenich |

**Table 6.** An overview of the methodology used for testing the fragrances.

| **Fragrance** | **Day<sup>1</sup>** | **n** | **Amount<sup>2</sup>** | **Hours between application and availability** |
| --- | --- | --- | --- | --- |
| Ethanol (control) | 1 | 2 | 2 sprays | 0 |
| Obsession for Men (source 1) | 1 | 2 | 2 sprays | 0 |
| Drakkar Noir | 2 | 1 | 2 sprays | 0 |
| Obsession for Men (source 2) | 2 | 1 | 2 sprays | 0 |
| L'Air Du Temps | 2 | 1 | 2 sprays | 0 |
| Paco Rabanne Pour Homme | 2 | 1 | 2 sprays | 0 |
| Drakkar Noir | 3 | 2 | 2 sprays | 0 |
| Civette Intense | 3 | 1 | 2 sprays | 0 |
| Obsession for Men (source 1) | 4 | 1 | 10 sprays | 10 |
| Drakkar Noir | 4 | 1 | 10 sprays | 10 |
| L'Air Du Temps | 4 | 1 | 10 sprays | 10 |
| Paco Rabanne Pour Homme | 4 | 1 | 10 sprays | 10 |
| Obsession for Men (source 3) | 5 | 2 | 2 sprays | 0 |
| Civetone <sup>3</sup> (0.1%) | 5 | 2 | 2 sprays | 0 |
| Obsession for Men (source 3) | 6 | 1 | 2 sprays | 0 |
| Obsession for Men (source 3) | 6 | 1 | 10 sprays | 10 |
| Civetone<sup>3</sup> (1%) | 7 | 1 | 2 sprays | 0 |
| Civetone (1%) | 7 | 1 | 10 sprays | 10 |
| Civet Absolute (synthetic) (1%) | 8 | 1 | 2 sprays | 0 |
| Civet Absolute (synthetic) (1%) | 8 | 1 | 10 sprays | 10 |

<sup>1</sup> The fragrances were tested on 8 different days. The same numbers in this column means these fragrances were tested on the same day.
<sup>2</sup> Although there is some variation between different atomizers, each spray is roughly equivalent to 100 µL.
<sup>3</sup> Natural civet is believed to contain about 1% civetone (Endallew and Dagne 2020).
n, number of fabrics available to the cats

### Chemical analysis of cat-attracting compounds

The methodology published previously (Bol) to extract and quantitate known cat-attracting compounds was optimized as described in detail below. Plant materials not already in powdered form were frozen with liquid nitrogen and powdered using a mortar and pestle. Powdered samples (circa 500 mg) were accurately weighed into glass vials in triplicate and 50 µg of internal standard (50 µL of 1 mg/mL benzofuranone in ethyl acetate, Sigma #124591) was added per 500 mg tissue (10 µg per 100 mg). This standard was chosen over the previously used tridecyl acetate standard because benzofuranone was found to be more stable over time and it does not elute in regions of the chromatogram where other analytes in the samples elute (especially *A. polygama* samples). Instead of 100% dichloromethane, we used 5% (v/v) methanol / 95% dichloromethane as extraction solvent, since we found through optimization that it is more effective at extracting the maximum amount of compounds of interest in 2 days versus up to 7 days using 100% dichloromethane. Five mL of extraction solvent was added, the vials were sealed and the content was magnetically stirred at room temperature for two days. One mL aliquots were filtered and subsequently analyzed by gas chromatography–mass spectrometry (GC-MS). Samples were analyzed on an Agilent 6890 instrument with autosampler connected to a 5973 mass selective detector. The samples were separated using a DB5-ms capillary column (50 m × 0.2 mm × 0.33 µm, J&W Scientific) using a flow rate of 0.7 mL/minute of ultra-high purity helium. The column initial temperature was 40⁰C, held for 1 minute, then increased to 130⁰C at 10⁰C per minute, then increased at 2⁰C per minute until 200⁰C, then finally at 15⁰C per minute to 280⁰C. The inlet temperature and transfer line were set at 250⁰C and the mass spectrometer was set to record between 40 to 250 amu.

All compounds measured were confirmed with external standards and quantified by calibrating the instrument using the internal standard method using the Agilent Enhanced Chemstation software (version D.01.02.16). The instrument was calibrated against standard concentrations ranging from 0.1 µg/mL to 100 µg/mL of the pure compounds relative to the internal standard benzofuranone (10 µg/mL).

### Statistics

The number of samples in this study was too small to assume normality of the data and therefore we used non-parametric tests for the statistical analyses, with the exception of paired analysis with missing data. The name of the test used for an analysis is mentioned in the text, immediately after a P value is reported. Dunn’s post-hoc test was always used for pairwise comparisons after the Kruskal-Wallis and Friedman test. All P values reported from Dunn’s post-hoc test are corrected for multiple comparisons. All analyses were done using GraphPad Prism version 9. Color schemes were selected using ColorBrewer (v2.0).

## Results

### The duration of the response to cat-attracting plants differs between cats

In a previous study, we tested the response of 100 domestic cats to *N. cataria*, *A. polygama*, *L. tatarica* and *V. officinalis* (Bol et al. 2017). Results from that study indicated that cats who did not respond to *N. cataria* (catnip) often responded to at least one of the other three plants. Because plants were available to the cats for up to only one hour, we limited our analysis to scoring the absence or presence of the “catnip response” and did not study their behavior in detail. Here we studied the response of 6 cats in their familiar permanent home environment to the same 4 plants used in our previous study, plus *Acalypha indica* (Indian nettle) (**Supplementary Figure 1B-G**), which has not been tested before to our knowledge. To allow for a comprehensive analysis of cat behavior in response to the cat-attracting plants, each plant was presented to the cats for a total of 100 hours, spread over 10 days (**Supplementary Figure 2A**). This dataset was analyzed for differences in (1) response duration and (2) behavior in response to these plants between (A) the cats and (B) the plants tested.

All but one of the 6 cats responded to all 5 plants tested (**Figure 1** and **Supplementary Figure 3**) and all responses to the plants could be classified as “catnip responses”, meaning the cats showed (a combination of) behaviors listed in Table 4. We observed approximately 2 hours of responses to *A. polygama* and *L. tatarica*, 1.5 hours to *N. cataria* and *A. indica*, and 1 hour to *V. officinalis*. Since 5 of the 6 cats in this study had never responded to *N. cataria* in the past, two different brands of catnip were used to investigate whether fluctuations in the level of active compounds in different sources of catnip could account for variation in (or lack of) attractiveness. One sock contained catnip from the brand Frontier, the other from the brand SmartyKat. When comparing the daily total response duration to both catnip brands for each cat separately, we observed that cat O responded significantly longer to the catnip from Frontier (**Supplementary Figure 4**). This finding suggests there may be a difference between the two brands of catnip that were used in this study, but overall, many and robust responses were observed from all 6 cats to catnip from both brands.

**Figure 1.**
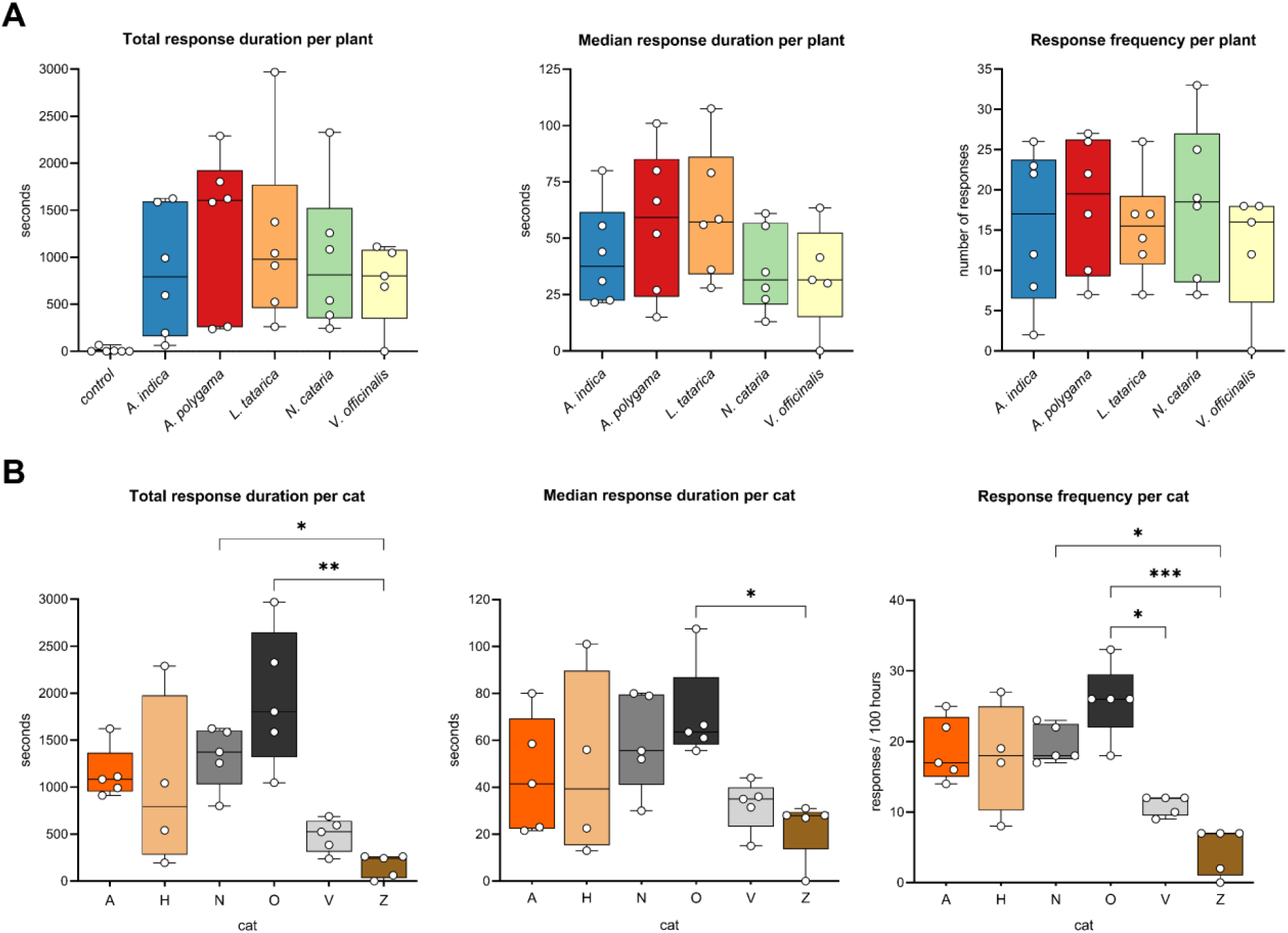
Response duration and response frequency of domestic cats to cat-attracting plants. Box and whisker plots showing the total response time, median response duration, and the total number of responses of 6 domestic cats to 5 cat-attracting plants. Each dot represents the data of one cat; the middle line in the bars shows the median value. Each cat-attracting plant was available for 100 hours, the control (green tea) was available for 500 hours (100 hours for each of the 5 plants tested). (**A**) Data shown per plant. Note the large spread of the data points, indicating large variation in response duration and frequency to the various plants between the cats. Differences between the 5 plants (total response duration, median response duration and response frequency) were not statistically significant (P > 0.05, mixed-effects repeated measures ANOVA and Tukey post-hoc test, corrected for multiple comparisons). We obtained 5 instead of 6 data points for *V. officinalis* since cat H was unable to participate due to medical reasons. For the statistical analysis of the paired data with missing data (cat H) we used a parametric test (mixed-effects repeated measures ANOVA). Therefore, for the analysis we used the average values (both the average response time to a plant for each cat and the average of the cats for each plant) instead of the median. Using either the average or median data did not affect the outcome of the statistical analysis. (**B**) Differences in total response time, median response duration, and response frequency between cats. Colors represent the fur color of the cats. Response duration and frequency differed significantly between the cats (Kruskal-Wallis). P values shown in the graph are from Dunn’s post-hoc tests. * P < 0.05; ** P < 0.01, *** P < 0.001

While previous work had suggested domestic cats respond euphorically to *A. indica* (Indian nettle) root in a similar fashion to catnip (Scaffidi et al. 2016), this plant has never been tested on cats in a controlled study. Since the cat-attracting effect of *A. indica* root quickly disappears after harvest (Scaffidi et al. 2016) and its geographical distribution does not extend to North America, roots were lyophilized immediately after collection on Christmas Island, Australia, in an attempt to preserve their effect on cats. Our data show that the response duration to the lyophilized roots of Indian nettle was similar to the other plants that were tested.

The cats only sparsely interacted with the negative controls (green tea). The total response time (any engagement with the object, not behavior specific to the “catnip response”) from all cats to the negative controls after 500 hours availability was just over 6 minutes, which is approximately 1% of the observed response time to the cat-attracting plant materials (490 minutes). Nearly all interactions with the negative control were from cat V and most of them occurred when *A. polygama* was tested. Three cats never engaged with the negative controls.

There was no statistically significant difference in total response time of the cats between the 5 plants (**Figure 1A**). Total response time is the sum of the duration of all responses, and is determined by both response frequency and response duration. We also did not find a statistically significant difference in the median response duration and response frequency of the cats between the cat-attracting plants.

However, when comparing the response duration to the 5 different plants between the 6 cats, we found these to be significantly different (**Figure 1B**). Cats O and N responded longer to the cat-attracting plants than cat Z. The differences in total response time to the cat-attracting plants between the cats could be explained by both differences in the length of the responses and the frequency of responses. These data show there are significant differences between cats in how long and frequently they respond to cat- attracting plants.

There was no statistically significant difference in response duration between the various plants, possibly because of the large variation between the cats. However, when we looked at the responses to the various plants for each cat individually, we observed that cat H responded significantly longer to *A. polygama* and cat O to *L. tatarica* and *N. cataria* than to some of the other plants (**Figure 2**). Interestingly, cat Z showed no interaction at all with the sock containing *V. officinalis* root over the full 5-week testing period.

**Figure 2.**
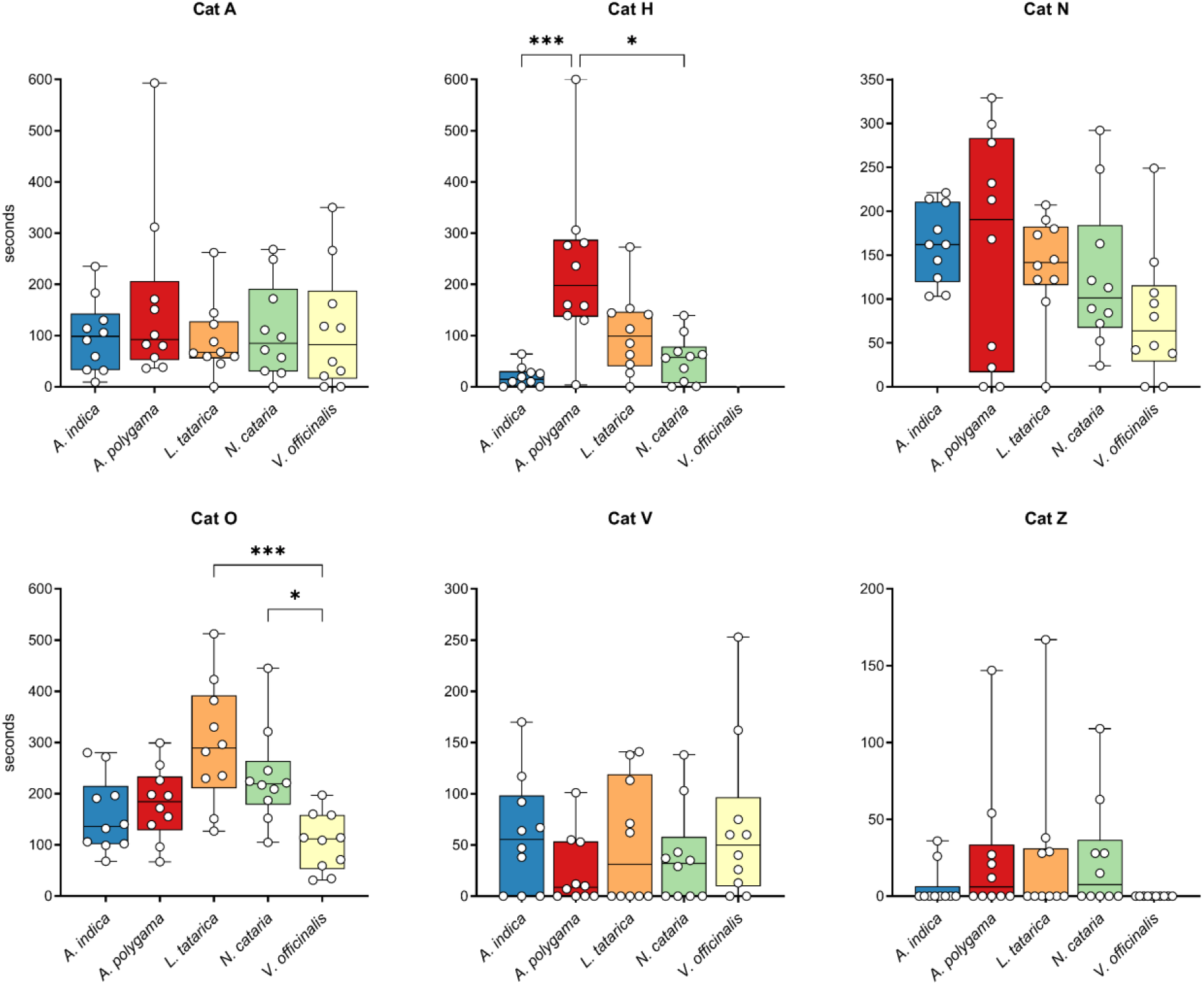
Response duration to cat-attracting plants shown for each cat individually. Each dot represents the total response duration of one day (10 hours), with the middle line in each box showing the median of these 10 days. Each plant was available for 10 days (total of 100 hours). Note that the Y axes are not the same for all graphs since the goal was to illustrate differences between the plants for each cat, not between cats. The Kruskal-Wallis test was used to test for statistically significant differences between plants. P values shown in the graph are from Dunn’s post-hoc tests. * P < 0.05; *** P < 0.001

The data also show that *N. cataria* (catnip) was not more popular than the other plants tested when comparing across the 6 domestic cats in this study. The longest total response duration after 100 hours, as well as the longest total response per day, and the longest single response was never to *N. cataria* (**Supplementary Figure 5**). These results suggest that while catnip might be the best-known cat- attracting plant among cat caregivers outside of East Asia, the other plants seem to be at least as potent. Behavior observed for cats O and V in response to the plant *Menyanthes trifoliata* (buckbean) suggests this plant is also able to elicit the “catnip response”. Fifteen grams of dried buckbean leaves (Siberian Herbals) inside a sock was offered to cats A, N, O, V and Z for a couple of hours on one day. We observed one response of cat O that lasted about half a minute and one response of cat V that lasted a little over one minute.

### The degree of attraction to cat-attracting plants differs between cats

Next, we looked at the degree of attractiveness of the plants. This was measured by the time it took a cat to respond to the plant for the first time after it was made available on each of the 10 test days. The data show no difference in attractiveness between the 5 plants we tested (**Figure 3A**). However, we did observe significant differences in how strongly individual cats were attracted to the plants (**Figure 3B**).

**Figure 3.**
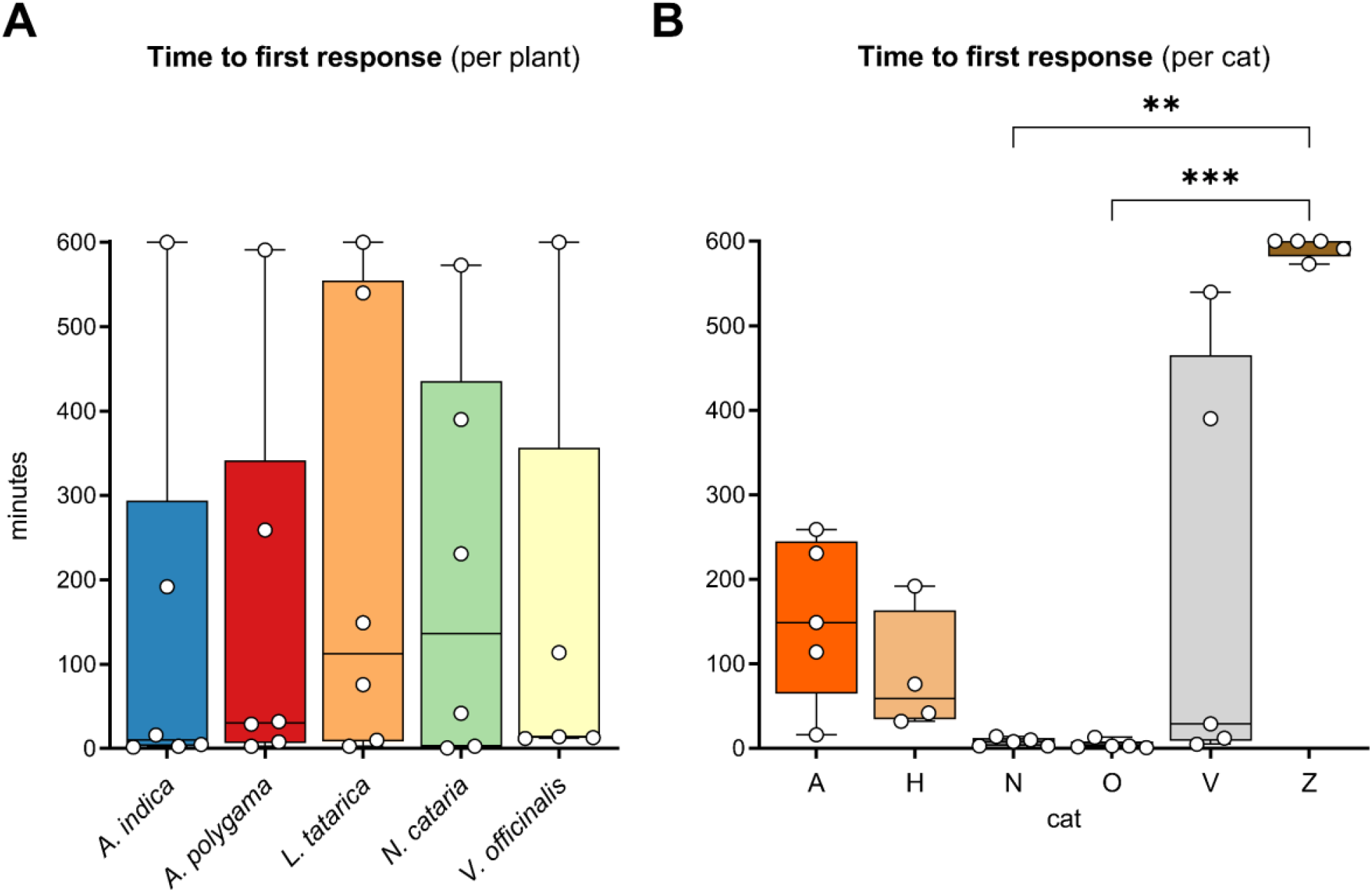
Time to first response. (**A**) The median time till the first response of 6 cats is shown for 5 cat- attracting plants. Each dot represents the median time till the first response of 10 testing days of each cat to the cat-attracting plants. Cat H did not participate in testing *V. officinalis*. There were no statistically significant differences in the time to the first response between the plants (P > 0.05, mixed-effects repeated measures ANOVA (paired test with missing data; see Figure 1A)). (**B**) The median time till the first response of 5 cat-attracting plants is shown for the 6 domestic cats. Each dot represents the median time to first response of 5 cat-attracting plants. The differences between the cats were statistically significantly different (Kruskal-Wallis). P values shown in the figure are from Dunn’s post-hoc test. ** P < 0.01, *** P < 0.001

These results suggest that the time to first response is in part determined by the cat’s personality (e.g., curiosity or fear of missing out), rather than intrinsic properties of the plant. Therefore, we also compared the times to first response to the 5 cat-attracting plants for each cat separately. Seeing differences in time to first response between the plants for individual cats may suggest differences in intrinsic properties between the plants. Similar to response duration, while we did not see differences between the time to first response when we looked at the combined data of all 6 cats, we did see statistically significant differences in time to first response between plants when we analyzed the data for each cat separately (**Figure 4**). While cat O did not have a single day out of the 50 without responding at least once, cat Z did not respond at all on about 70% of the days, including the 10 days *V. officinalis* was available. Cat O responded to *L. tatarica* and *N. cataria* almost immediately on each of the 10 test days. In contrast, the first response to *V. officinalis* of cat O was about 9 hours on three of the 10 test days. The opposite was seen for cat V, who appeared to be attracted more strongly to *V. officinalis* than to *N. cataria*. On all 9 days that cat V responded to *V. officinalis*, this was within or around half an hour. These results suggest that the level of attractiveness of a plant is not solely determined by properties of the plant, but also by how the cat perceives the plant.

**Figure 4.**
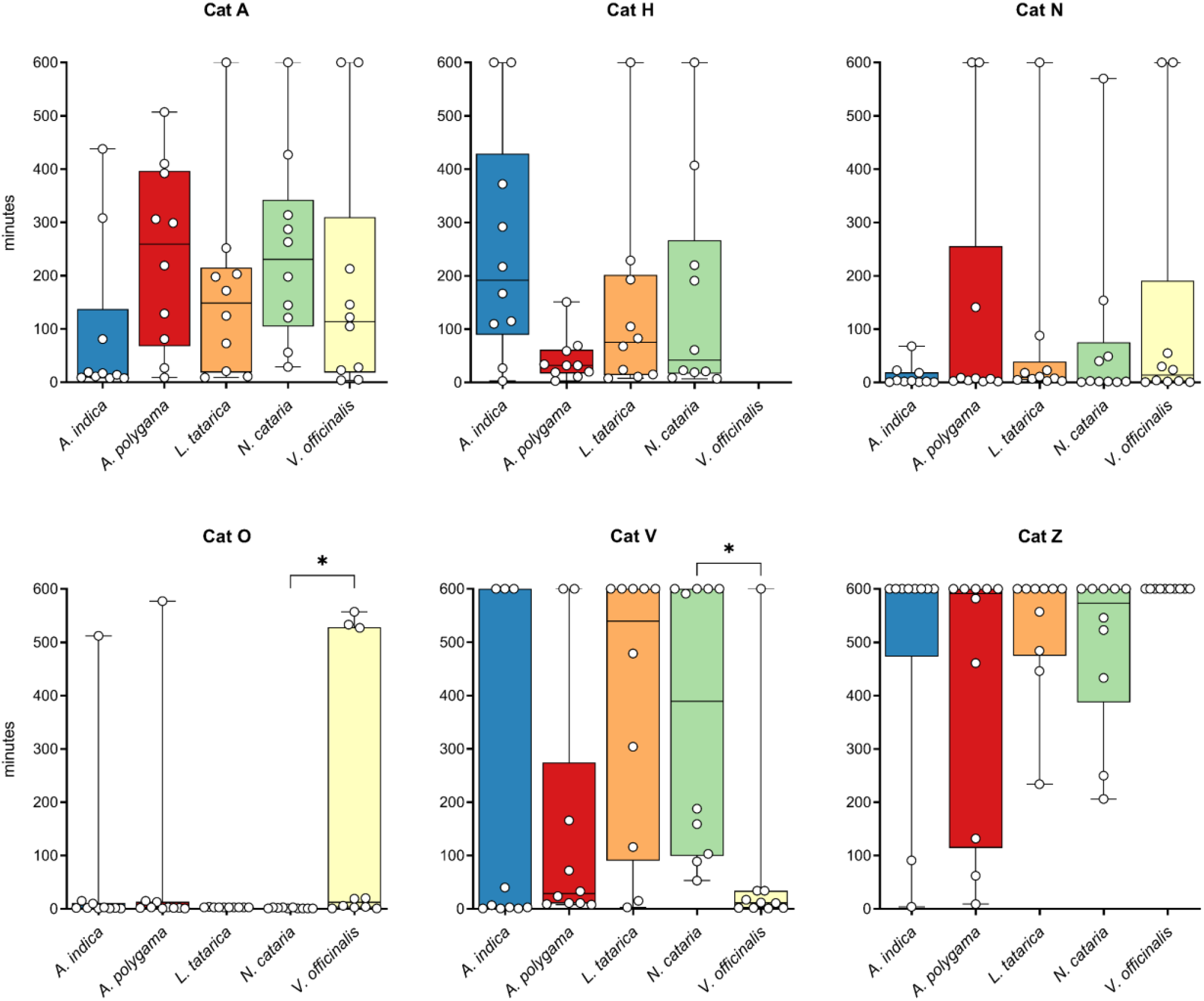
Time to first response to 5 cat-attracting plants shown for each cat separately. Each dot shows the time it took the cats for their first response on each of the 10 test days. Cat H did not participate in the testing of *V. officinalis*. * P < 0.05

Taken together, these data show that all 5 plants are equally capable of attracting domestic cats and eliciting the “catnip response”, while both response duration and how strongly individual cats are attracted to the plants can differ significantly. These differences might in part be due to variation in olfactory perception and in part to differences in the cats’ personalities.

### The “catnip response” is different between cats, but comparable among various cat-attracting plants

In addition to the quantitative analysis (i.e., duration of the response) we also studied the qualitative aspects of the responses to the various plants. We created an ethogram that is specific for the “catnip response” (**Table 4**). Some of these behaviors may be affected by how the olfactory stimulus is offered to the cat. For example, biting and pulling with the object in the cat’s mouth will be possible when the plant material or single compound is offered inside or on a fabric, respectively, but it will not be observed when powder of dried *A. polygama* fruit galls is sprinkled on the floor. In this study, all plant materials and single compounds were offered on or in a fabric and therefore allowed for comparison between cats, as well as between plants or single compounds. Behaviors not mentioned and described in the ethogram either did not occur (Flehmen, lordosis, vocalization) or were not analyzed because of limitations such as camera angle and distance (e.g., drooling). Sniffing was not included because it was considered behavior used to detect or identify an odor, not behavior in response to smelling odorants. Although not specifically studied or analyzed, no signs of stress, fear or aggression (determined by for example positioning of the ears or tail) were ever observed. In addition to previously described behavior in response to catnip, we have added “rippling of the back” and “twitching of the back” (**Supplementary File 1**). This behavior is not linked to feline hyperesthesia syndrome. There is no reaction (biting, scratching or licking of the area where the twitching or rippling occurs) of the cats to the concerning area of the back, rather, the cat seems completely unaffected by it. Twitching and rippling of the back appeared to be quite specific for the “catnip response” since it was only rarely observed on other occasions. “Rolling on the side” reflects the frequency of changes in body position (standing/sitting to lying on the side or lying on the side to lying on the back). Rippling and twitching of the back, as well as rolling on the side and head shaking are extremely short events and are therefore reported and shown as events per minute response, whereas all other behaviors are reported and shown as the percentage of the total response time. The percentages can exceed 100% since some behaviors can be displayed by the cats simultaneously (e.g., holding and rubbing, or, holding and raking).

To compare behavior between the cats, we analyzed 5 responses to *N. cataria* nearest to 60 seconds of each cat using BORIS behavioral analysis software. Catnip was chosen because the variation in frequency and length of the responses of the 6 cats was least for this plant. During the response, the cats were mostly either sitting or lying on their side. Time spent while standing or lying on their back during the response was also observed, but not frequently (**Figure 5**). Body position during the response varied enormously between the cats. Cat O predominantly lay on his side while engaging with the filled sock, cats A, H and Z responded predominantly in a sitting position, and cats N and V showed an equal mix of sitting and lying on their side (**Figure 6**).

**Figure 5.**
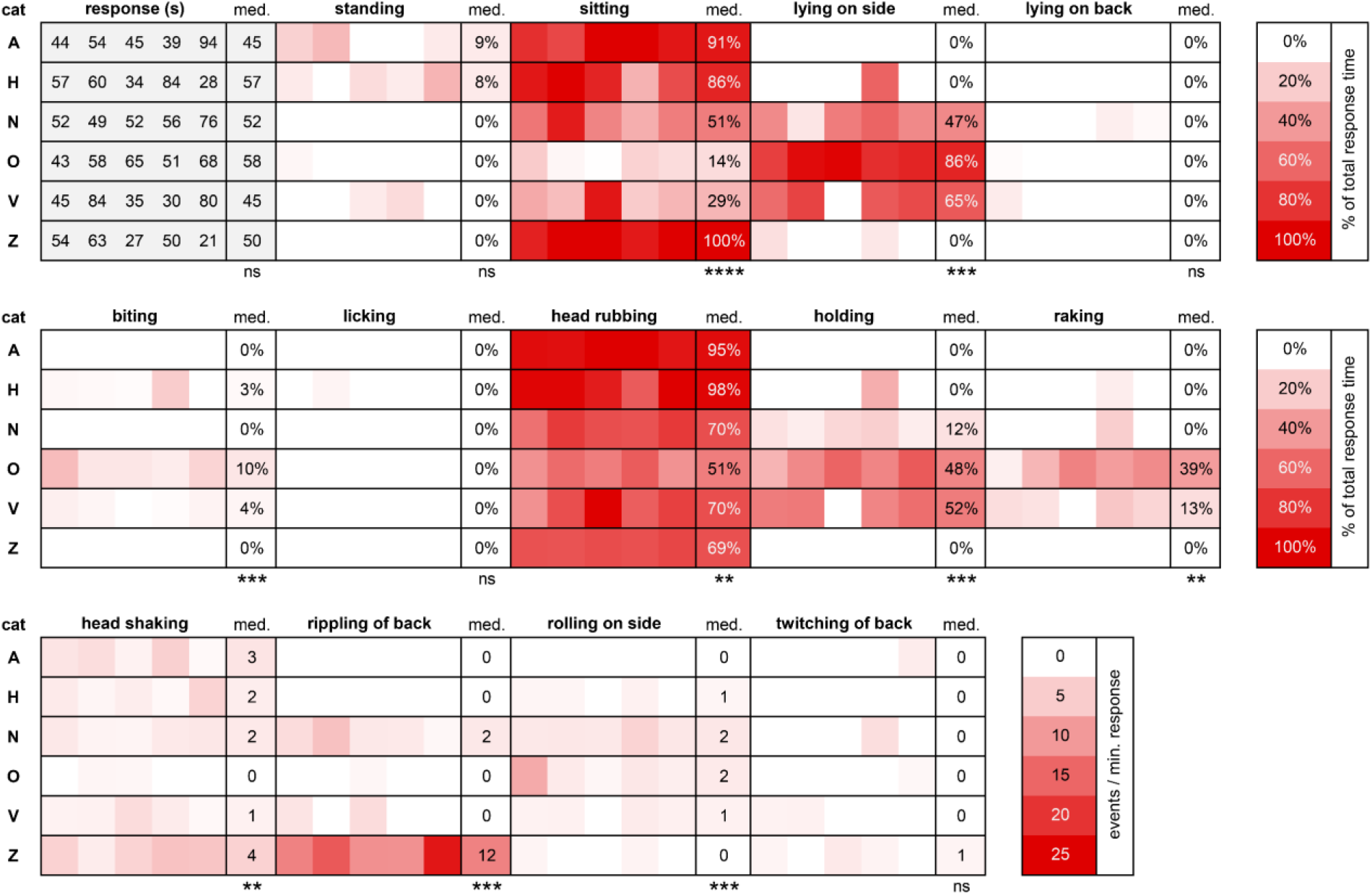
Heatmap showing similarities and differences in behavior between 6 domestic cats in response to *N. cataria* (catnip). For each cat, the five responses nearest to 60 seconds were analyzed using BORIS behavioral analysis software. All P values shown are from the Kruskal-Wallis test. med, median; ns, not statistically significantly different; ** P < 0.01; *** P < 0.001; **** P < 0.0001

**Figure 6.**
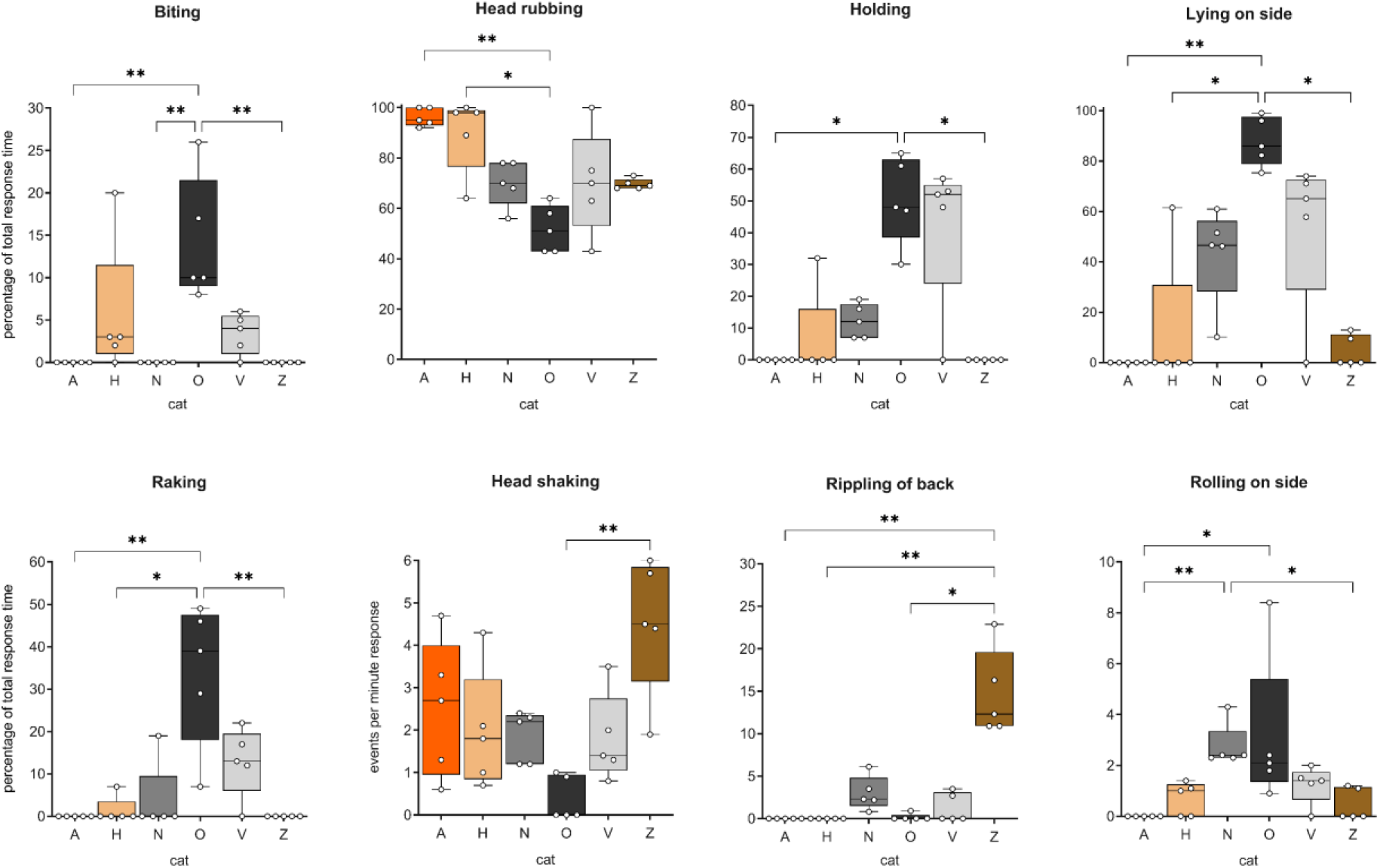
Body position and behavior of 6 domestic cats observed in response to *N. cataria* (catnip). Results for “biting”, “head rubbing”, “holding”, “lying on side”, and “raking” are shown as time spent relative to the total response duration (percentage), whereas results for “head shaking”, “rippling of back” and “rolling on side” are depicted as the number of events per minute of response. Data for the body position “sitting” is not shown because sitting and lying down were mutually inclusive and inversely correlated (Figure 5). All P values shown are from Dunn’s post-hoc tests. * P < 0.05; ** P < 0.01

Our data also suggest there is large variation between cats in most behaviors that are typical for the “catnip response”. Head rubbing the olfactory object was the behavior observed most frequently, and although it was seen for all 6 cats, there were significant differences between the cats (**Figure 6, Supplementary File 4**). The response to *N. cataria* for cats A and H consisted almost exclusively of head rubbing, significantly more than for cat O. In addition to head rubbing, cat O showed other behaviors such as raking or biting while holding the object. The amount of time spent holding the sock, raking and biting was significantly greater for cat O than for several of the other cats (**Figure 6**). Rippling of the back was not seen for cats A and H but was a characteristic feature of cat Z’s response, where it was seen at high frequency (**Figure 6**). In fact, about 15% of her response time was rippling of the back. Head shaking, rolling on the side, and twitching of the back was seen for most or all cats, with no differences between cats for the latter. The frequency of head shaking was significantly different between the cats O and Z (**Figure 6**). This behavior seemed to be rather specific for the “catnip response” since it was not seen during their normal daily activity. None of the cats had medical problems with their ears, nor did we observe any buildup of wax in their ear canal to account for head shaking. We also did not see any scratching or pawing aimed at the head or ears, which would be indicative of medical problems with the ears. Perhaps this head shaking behavior is similar to “shake-off” behavior seen in dogs where it can serve as a “reset button” after excitement, although there is no literature that would support this hypothesis. Alternatively, it might be a way for the cats to shed excess saliva, since it is known that these cat-attracting plants can induce drooling (Bol et al. 2017).

Overall, the frequency of rolling on the side was low. The responses of cats N and O seemed more dynamic than the response of cats A and Z since rolling on the side from a sternal position, or onto the back from a lateral position, was seen more frequently with cats N and O (**Figure 6**).Collectively, these data demonstrate that the behavior seen in the “catnip response” is quite consistent for each cat, but show enormous variability between cats.

Having observed large variation in response traits of domestic cats towards catnip, we wondered if their idiosyncratic behavioral pattern would be the same for all the various cat-attracting plants used in this study. As can be seen in **Figure 5**, the behavioral pattern in response to *N. cataria* is quite distinct between cats A, O and Z. Cats A and Z have a fairly simple behavioral response where they predominantly sat and head rubbed the object, with cat Z also frequently demonstrating rippling of her back. On the contrary, cat O spent much more time lying on his side, raking, biting, and holding the object, and rolled on his side much more frequently than the other two cats. To test if there is a difference in behavioral patterns of cats towards different cat-attracting plants, we analyzed the behavior of cats A, O and Z in response to all plants tested in this study.

During the response of cat A to any of the 5 plants, she predominantly sat and head rubbed the filled socks (**Figure 7A**). While some licking was seen during some of her response to *A. polygama* and *V. officinalis*, the body position and behaviors of cat A were highly similar between catnip and the 4 other plants.

**Figure 7.**
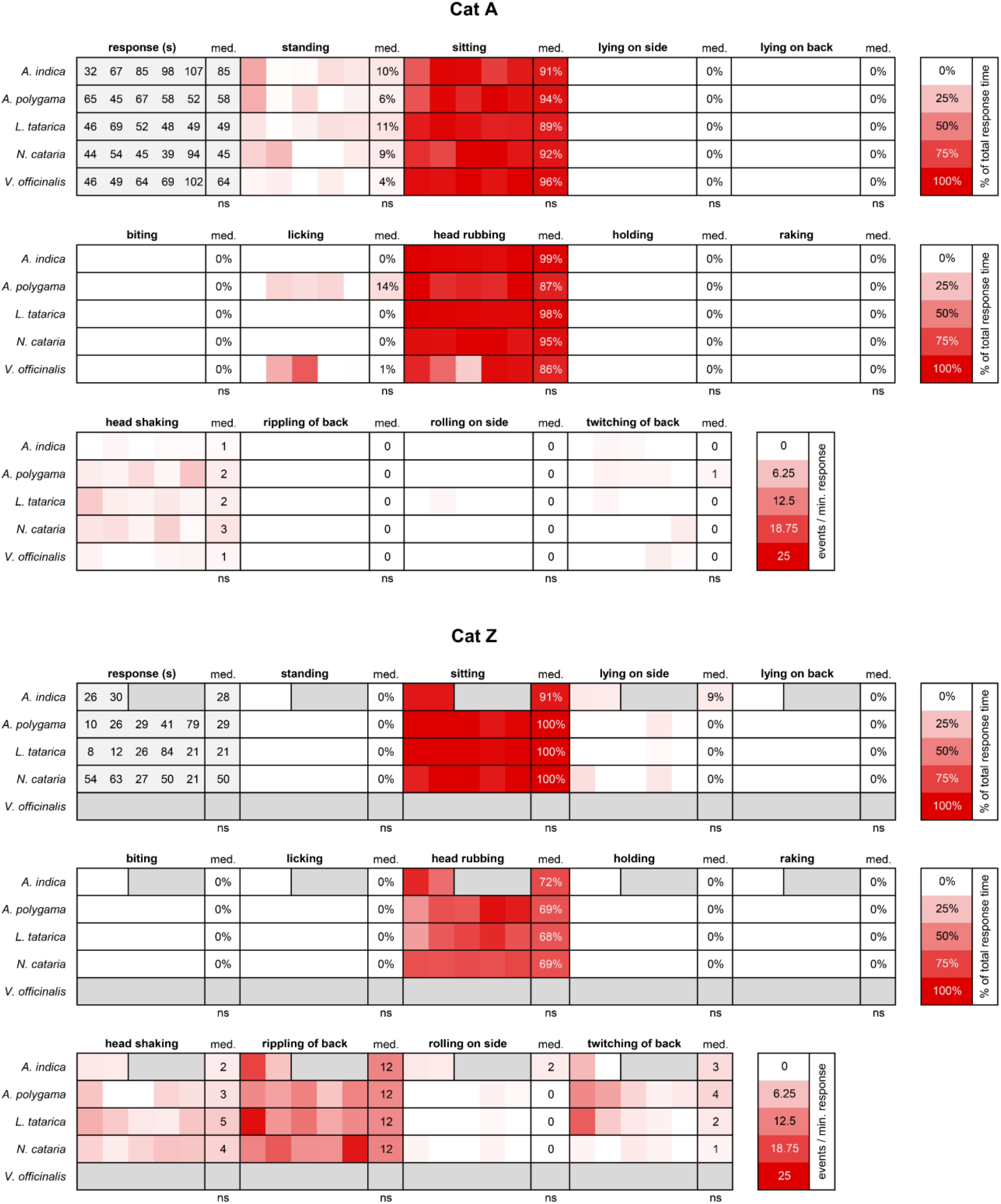
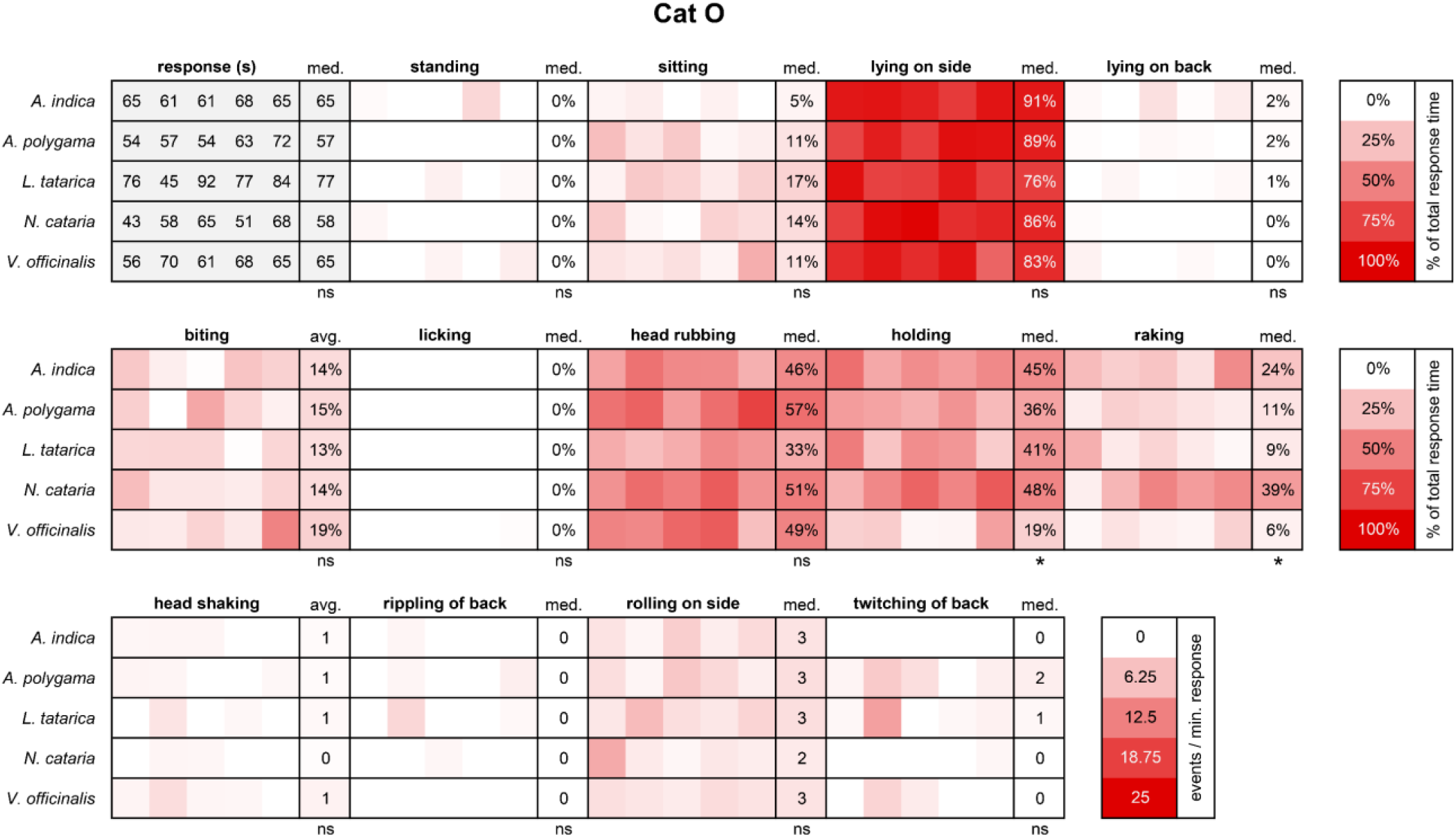
Body position and behavior observed during the response to various cat-attracting plants. For cats A, Z and O (Figure 7A, **B** and **C**, respectively), the five responses nearest to 60 seconds were analyzed using BORIS behavioral analysis software. We observed only two responses from cat Z to *A. indica*. Therefore, two responses instead of 5 were analyzed. P values shown are from the Kruskal- Wallis test. med, median; ns, not statistically significantly different; * P < 0.05

We observed lots of rippling of the back for cat Z in response to *N. cataria*. Behavioral analysis revealed that rippling of the back was not specific for catnip, but rather part of her general response since it was observed in response to all cat-attracting plants (except *V. officinalis* to which she never responded) (**Figure 7B**). In addition to rippling of the back, we also observed twitching of the back in response to all the other plants tested. It is unknown whether rippling of the back (wavelike motion) and twitching of the back (single contraction on one location lasting a fraction of a second) are related. Her body position and behavior during the responses to the other cat-attracting plants were highly similar in proportion and frequency when compared to catnip.

Finally, we compared the behaviors of cat O between the 5 different plants. His response to *N. cataria* was the most diverse and complex out of all the 6 cats with him predominantly in a lateral position (∼85% of the response time) when head rubbing (∼50%), raking (∼35%) and biting occasionally (∼15%) while holding the object (∼50%). Cat O rolled on his side from a sternal position 2 – 3 times per minute response duration, and we rarely observed headshaking (without the sock in his mouth), and rippling or twitching of his back. In line with what we observed for cats A and Z, his behavioral pattern was near identical for all cat-attracting plants (**Figure 7C**). The data also suggest however, that holding and raking was seen less frequently for cat O when responding to *V. officinalis*, especially when compared to *N. cataria* (**Figure 7C** and **Supplementary Figure 6**). These findings are interesting when considering the previous observations that cat O was significantly less attracted to *V. officinalis* root than to *N. cataria* (**Figure 4**), and that his total response duration to valerian root was also less than to other cat-attracting plants (**Figure 2**).

Taken together, these data suggest that while responses between cats vary, the behavior of individual domestic cats to diverse cat-attracting plants is highly similar, although the effect of *V. officinalis* root on cats seems to be slightly different.

### Response duration to cat-attracting plants decreases with repeated exposure

The setup of the experiments, with its repeated presentation, allowed us to learn more about possible habituation (reduced response duration over time to the same stimulus) to the cat-attracting plants.

Information about possible habituation will be useful when giving advice to cat caregivers on how to use olfactory stimuli for environmental enrichment. Furthermore, seeing differences in habituation between plants might suggest the presence of different compounds or quantities of these compounds in the cat- attracting plants.

The olfactory stimuli were offered 2 – 3 days a week, for 10 hours a day, for two periods of two weeks (weeks 1 – 2 and 4 – 5), with an interstimulus interval of at least 9 days between weeks 2 and 4 (**Supplementary Figure 2A**). First, we compared the total response time (median of 6 cats) during the first two-week testing period (weeks 1 and 2) with the second two-week testing period (weeks 4 and 5). When we analyzed all 5 cat-attracting plants together, we found that the median response time was the same (**Figure 8A**). We observed a similar pattern when we looked at the plants individually, suggesting that either no habituation occurred within the 5-week testing period, or that the one-week interstimulus interval was sufficient to reverse any habituation that may have occurred during the first two-week testing period.

**Figure 8.**
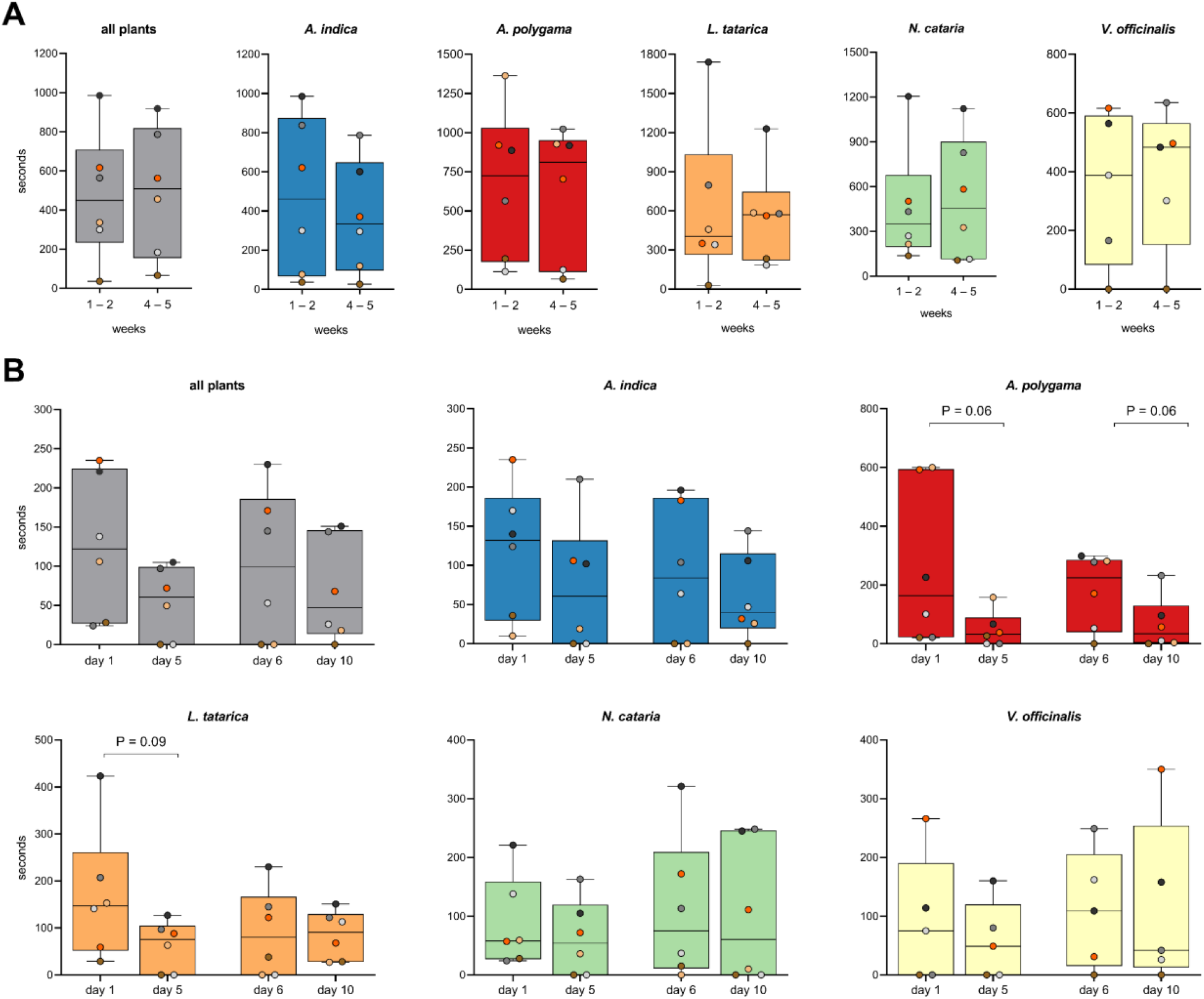
Response duration to cat-attracting plants over time. Each dot represents data (total response time) of one cat. When all plants were compared, each dot shows the median value of the total response durations to the 5 cat-attracting plants. (**A**) The total response duration of 6 cats to 5 cat- attracting plants during the first 5 testing days (50 hours; weeks 1 – 2) was compared to the total response time during the 5 testing days (50 hours) during weeks 4 – 5 (**Supplementary** Figure 2A). The test periods of two weeks were separated by a 9-day interstimulus interval. (**B**) Total daily response time of 6 cats during the first (day 1 and 6) and last day (days 5 and 10) of both two-week testing periods. Cat H did not participate in testing *V. officinalis*. For all statistical analyses the paired, non-parametric Wilcoxon matched-pairs signed rank test was used. All P values were > 0.05. Only P values < 0.1 are shown.

To test the latter, we compared the response duration between day 1 and day 5, as well as between day 6 and day 10. While none of the observed differences were statistically significant, we did see a decline in response time to *A. polygama* within both the first and the second two-week testing period (**Figure 8B**).

The response duration on the last day of both 5-day testing periods (days 5 and 10) was shorter for nearly all cats, suggesting that some habituation may have occurred. The response duration to this plant was the highest of all plants tested on the first day of both 5-day testing periods.

To learn more about possible habituation to the various stimuli, we performed additional experiments where the plant material was offered 10 days in a row for 2 or 12 hours per day. To rule out the effects of potential degradation or complete volatilization of the active compounds over time, two new socks with fresh plant material were offered every day. Habituation was observed for *A. polygama* (dried fruit gall powder) and *L. tatarica* (sawdust) (**Figure 9**, days 1 – 10). A similar pattern was seen for *N. cataria* (dried, cut leaves), but the difference between day 1 and day 10 was not statistically significant. We did not have enough material to also test *A. indica*. For all plants tested, after 1 to 1.5 weeks of daily, voluntary exposure (2 or 12 hours a day), the response duration of each cat was reduced to (close to) zero. After the 10-day testing period and possible habituation to the plant materials, a different cat-attracting plant was offered to learn if the scent from this stimulus would result in the reappearance of the response. This dishabituation would suggest the presence of other active compounds or higher levels of similar compounds in the newly offered stimulus. After habituation of the cats to either *L. tatarica*, *A. polygama* or *N. cataria*, no dishabituation was seen when the cats were offered different cat-attracting plant material (**Figure 9**). The only exception was cat O, who showed a longer response to *L. tatarica* than his first and longest response to *A. polygama* and *N. cataria* (**Figure 9A+D**), underscoring the idiosyncrasy between cats. Furthermore, these results suggest that *L. tatarica* may contain compounds not present, or at significantly lower amounts, in catnip and silver vine. Another interesting finding was the observation that offering *N. cataria* to the cats who were habituated to *A. polygama* and *L. tatarica* did not significantly increase response duration. This might suggest that nepetalactone binds to (some of) the same olfactory receptor(s) as some of the active compounds present in *A. polygama* and *L. tatarica*. These findings also indicate that offering cat-attracting plants on a non-continual basis or alternating between the various cat- attracting plants could prevent or reduce habituation in cats.

**Figure 9.**
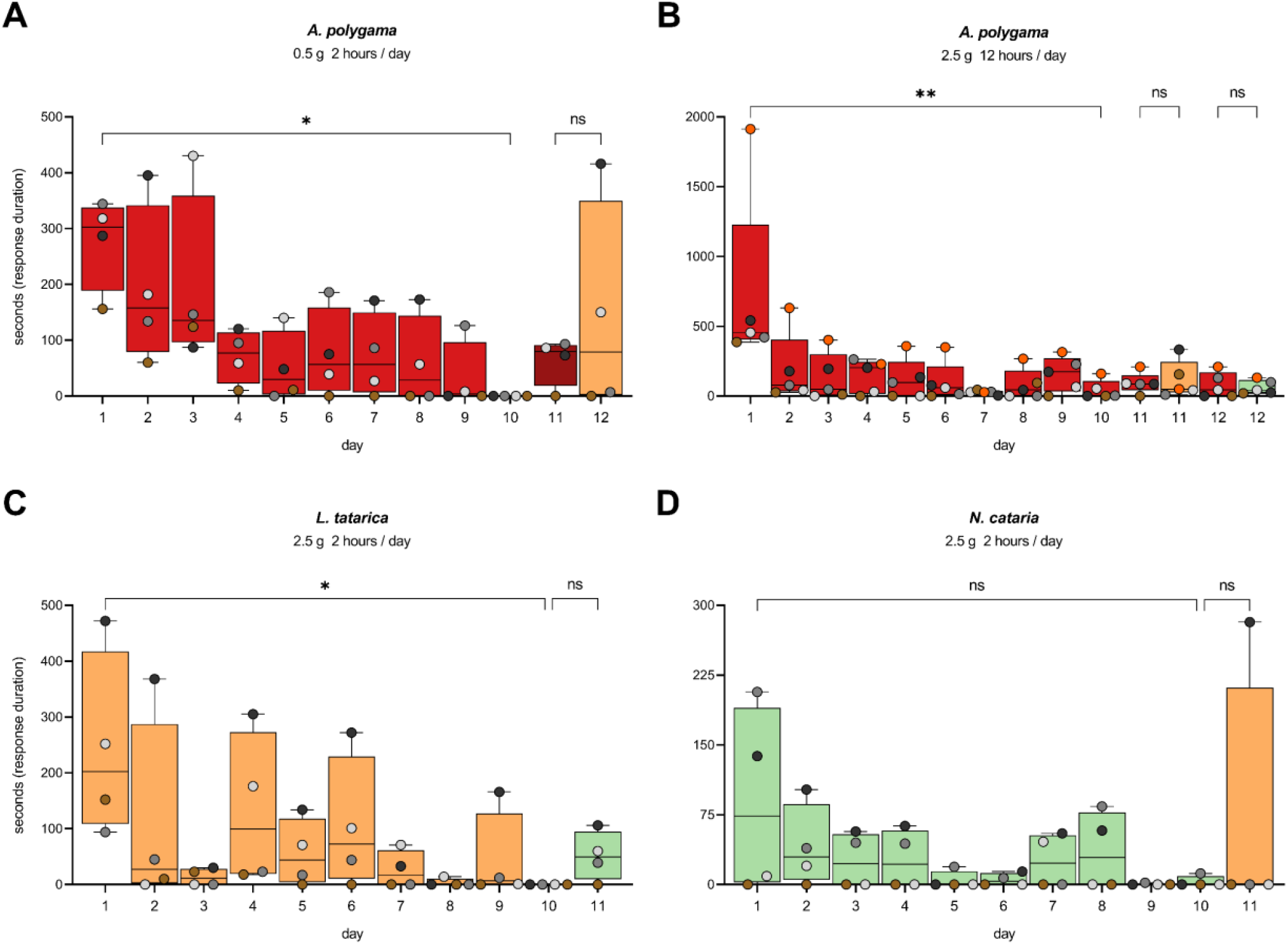
Habituation and dishabituation to cat-attracting plants. The response duration of 4 – 5 domestic cats to three different cat-attracting plants is shown for 10 consecutive days. With habituation a gradual decrease in response duration over time is seen. Dishabituation is the reappearance or increased duration of a response to a different stimulus that is offered to the cats after habituation has occurred (day 11 or 12, or both) and its duration is similar or higher to what was seen on day one. Results for *A. polygama* (**A** and **B**) are shown in red, for *L. tatarica* (**C**) in orange, and for *N. cataria* (**D**) in green. See **Supplementary** Figure 2B for more details. The differences between day 1 and 10 for *A. polygama* and *L. tatarica* were statistically significant (Friedman test). P values shown in the figure are from Dunn’s post- hoc test. * P < 0.05; ** P < 0.01

### Cat-attracting compounds in *A. polygama* are not exclusively produced in response to the parasitic attack of the gall midge *P. matatabi*

Both normal *A. polygama* fruit and fruit galls used in our previous study (Bol et al. 2017) were collected from vines growing in East Asia. In this natural habitat of the plant, gall midge *Pseudasphondylia matatabi* females can lay their eggs in the plant’s flower buds. As a result of this parasitic invasion fruit galls develop. It seems that the presence of *P. matatabi* larvae in the developing kiwi fruit is critical for the synthesis of compounds that serendipitously attract cats, since we have previously shown that domestic cats respond to dried *A. polygama* fruit galls, but not to dried normal fruit (Bol et al. 2017). Indeed, we were able to detect *P. matatabi* DNA in dried fruit galls that we used in our preceding study (**Figure 10A**). Sequencings results confirmed, unequivocally, that *P. matatabi* DNA was present in the *A. polygama* fruit galls (100% percent identity and query coverage; **Supplementary File 2**).

**Figure 10.**
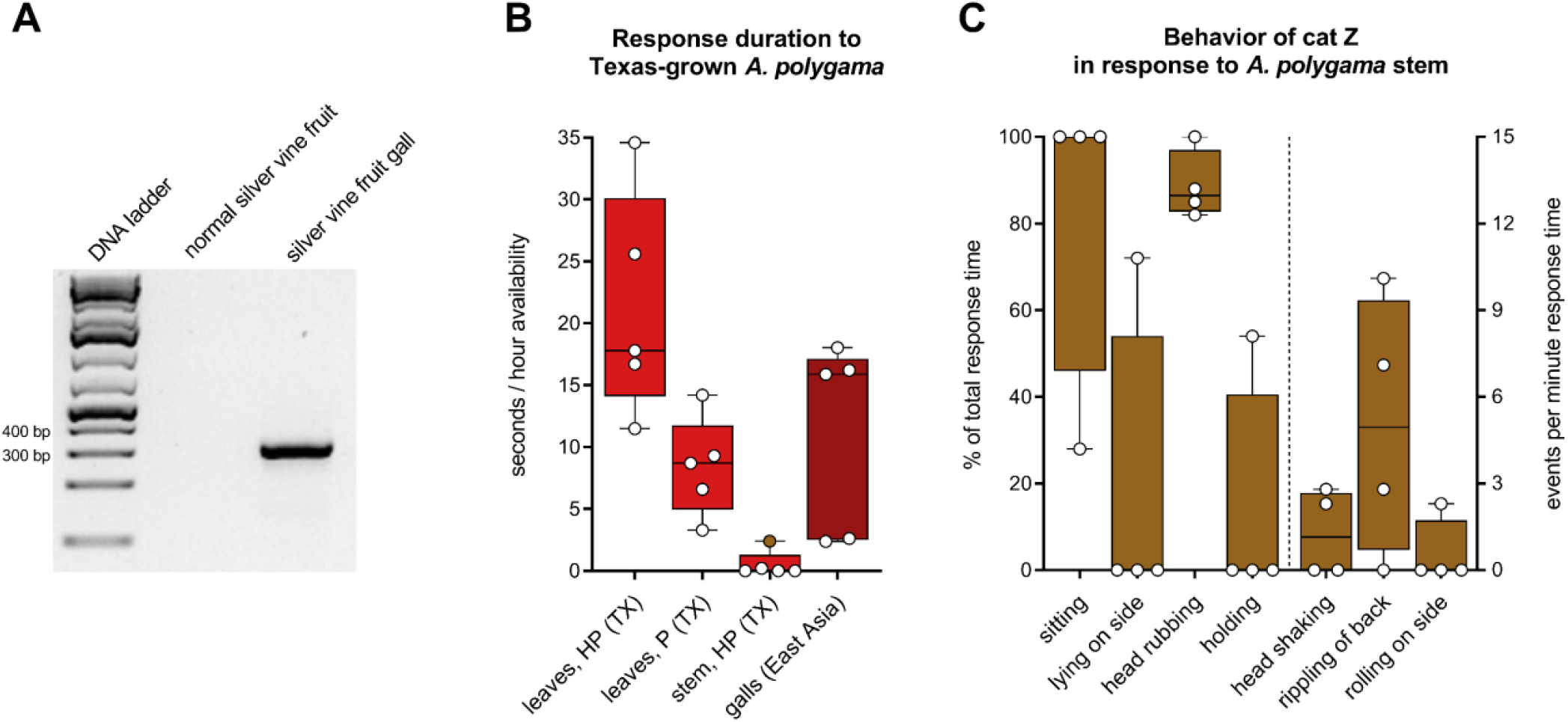
Response of domestic cats to Texas-grown *A. polygama*. (**A**) Detection of *P. matatabi* DNA in dried *A. polygama* fruit galls from East Asia. Species-specific primers were used to amplify a 330 bp fragment of the mitochondrial cytochrome oxidase subunit 1 gene. Sanger sequencing and nucleotide BLAST confirmed the DNA was from the gall midge *P. matatabi*. (**B**) Response time, shown in seconds per hour availability, of 5 cats to Texas-grown silver vine plant material. The cats were offered dried leaves from a female and male silver vine variety (Hot Pepper and Pavel, respectively), as well as dried, lignified stem. The response time to dried, powdered *A. polygama* fruit galls originating from East Asia is shown in dark red. Hot Pepper and Pavel leaves were available to the cats for 15 and 16 hours, respectively. Stem was available 2 × 15 hours. Powdered silver vine galls were available for 100 hours total (10 days, 10 hours per day). (**C**) Observed behavior of cat Z in response to Texas-grown *A. polygama* stem (brown dot in panel B). Bars show either behavior expressed as the percentage of the total response time (left Y axis) or the number of events per minute response time (right Y axis; “head shaking”, “rippling of back”, and “rolling on side”). Cat Z responded 4 times to the locally grown silver vine stem, with a total response time of 74 seconds. Only observed behavior is shown. HP, *A. polygama* Hot Pepper variety; P, *A. polygama* Pavel variety; TX, Texas

We wondered if the gall midge induces the synthesis of these compounds only locally (fruit) or systemically (stem, leaves, fruit). It is known that some domestic cats do respond to dried *A. polygama* stem (Bol et al. 2017). However, we do not know if these tissues were obtained from silver vine plants in East Asia that were bearing fruit galls at the time of harvest. Since *A. polygama* is dioecious and *P. matatabi* females deposit their eggs in the flower buds, not the fruit, one could argue that in response to oviposition in a male flower bud the plant might also systemically induce synthesis of cat-attracting compounds. However, *P. matatabi* oviposition in male flower buds or male flower bud galls have never been observed (Dr. Junichi Yukawa, Kyushu University, Fukuoka, Japan, personal communication, June 2021). To test whether the presence of the gall midge is required for the synthesis of the cat-attracting compounds, we grew *A. polygama* locally (Mico, Texas, USA), where *P. matatabi* does not occur. The cats were offered dried leaves from the female Hot Pepper variety and the male Pavel variety, each for almost a full day. Seeing cats respond to leaves from male plants, even when grown in their natural habitat and hence in the presence of *P. matatabi*, would suggest that the gall midge is not required for the production of these compounds. All five cats responded to the locally grown *A. polygama* leaves, both from the male and female plant (**Figure 10B**). Although the data are limited, they strongly suggest the leaves were at least as popular among the domestic cats as the dried gall material from East Asia. The shorter response to the leaves from the Pavel variety may be explained by harvesting later or the longer drying time of the leaves. Harvest time for those leaves was later in the fall when the leaves would soon be shed by the plant. Testing these already collected leaves was postponed because we wanted cat A, who had recently received radioactive iodine treatment for hyperthyroidism, to also participate. Stem from the female silver vine Hot Pepper variety was made available to the cats on two different days. In agreement with our previous findings (Bol et al. 2017), only a small percentage (20%) of the cats responded to the silver vine stem. Cat Z responded 4 times: 26, 8, 18 and 22 seconds, with a total response time of 74 seconds, and analysis of her behavior showed that the response was similar to the behavior observed when exposed to the other cat-attracting plants: mostly head rubbing in a sitting position with her back rippling and an occasional head shake (**Figure 10C**). No responses were seen to the control stem (lignified *Juniperus ashei*). Interestingly, while cat Z responded for a total time of 4 minutes and 15 seconds to the dried leaves of the Hot Pepper variety, she did not touch the sock containing the leaves for approximately half of that time. No other responses where there was no contact with the test object by cat Z or any other cat to any plant material were seen. Instead of contact with the object, she rubbed her head on the floor, rolled on her side, and her back rippled, all in close proximity (approximately 20 cm) to the olfactory object. This observed behavior in response to the dried silver vine leaves was characteristic for her and highly similar to her responses to other plants. This cat never demonstrated this behavior in response to any of the controls, which were available for hundreds of hours, and her most recent response prior to these responses was 3.5 weeks earlier. Therefore, we concluded this response was specific to the *A. polygama* leaves.

We previously concluded that domestic cats do not respond to *A. polygama* leaves grown in the USA (Bol et al. 2017). However, subsequent DNA barcoding (*matK*) revealed that the leaves previously used for testing were from the closely related species *Actinidia arguta* instead of *Actinidia polygama*. These *A. arguta* leaves were only used for one small experiment in our previous study, and this finding does not change any of the main or other conclusions of the published work. DNA barcoding (*matK*, *rbcL* and *psbA* – *trnH*) results strongly suggest we have used *A. polygama* for all experiments in this study, although we could not rule out the closely related *A. valvata*. Since the use of Tatarian honeysuckle wood as olfactory enrichment for cats is still uncommon and, as far as we know, is only available from one source (The Cat House in Calgary, Alberta, Canada), we also used DNA barcoding (*matK*, *rbcL* and *psbA* – *trnH*) to confirm that what we used in this study was indeed *Lonicera tatarica*. All sequences can be found in **Supplementary File 2**.

In conclusion, these findings show that while the gall midge *P. matatabi* seems to induce a change in the plant’s volatile pattern in the kiwi fruit gall, oviposition in the flower buds does not seem to be required to develop the cat-attracting characteristics of the stem and leaf tissues in either male or female silver vine plants.

### Active compounds in plants can be extracted using ethanol

We created *N. cataria*, *L. tatarica* and *V. officinalis* tinctures to determine whether this easy extraction method would result in a product that could attract and stimulate domestic cats. A liquid (ethanol) form would offer several possible advantages over the plant form since it can be applied to any object. *A. indica* and *A. polygama* tinctures were not created because of limited availability of plant material. We were also curious to see if we could extract any active compounds of dried *V. officinalis* root with absolute ethanol, and possibly avoid co-extracting any compounds that may have had an inhibitory effect on cat Z. She was the only cat who did not respond to *V. officinalis*, despite the plant being available for 10 days, 10 hours a day. We hypothesized that cat Z did not respond because dried *V. officinalis* roots have, at least to most humans, a strong, unpleasant or repulsive smell.

We applied two sprays of the tincture and two sprays of ethanol only (negative control) on a piece of fabric which were subsequently made available to five cats for a total of 5 hours in the afternoon / evening. We observed positive responses of two to four cats to each tincture (**Figure 11**), and the responses to them matched the “catnip response” behavior that was characteristic for each cat.

**Figure 11.**
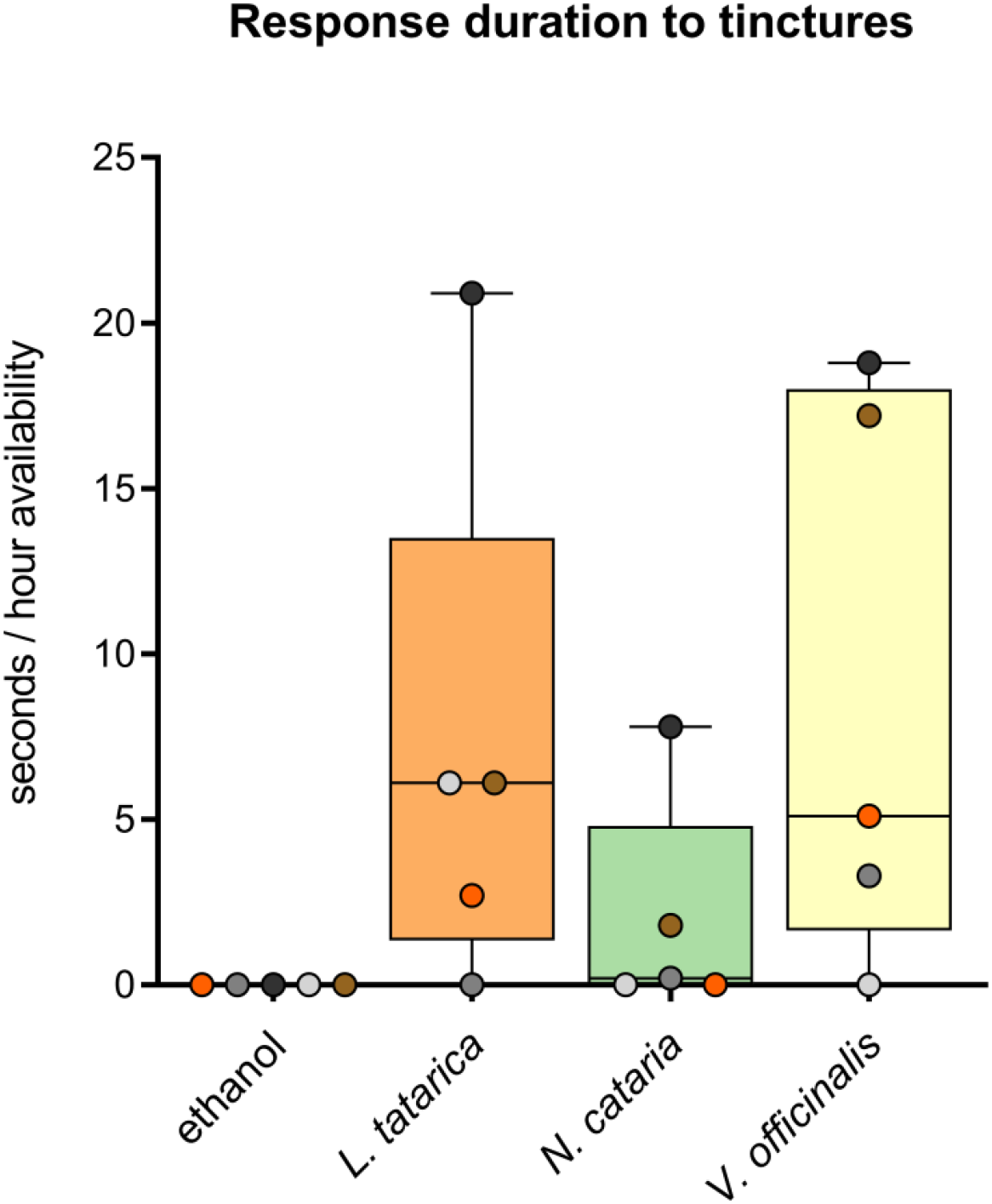
Response time of domestic cats to tinctures made from cat-attracting plants. Box and whisker plot showing the median response time of 5 cats (horizontal line) and median response time of each cat (dots). The response time is shown as time per hours availability of the tinctures. Each tincture was available for 5 hours. Ethanol was used as a negative control. The response duration of cat Z to the *V. officinalis* tincture is shown as a brown dot (18 seconds/hour availability). This cat did not respond at all to 15 g dried valerian root that was available for 10 days, 10 hours per day.

Interestingly, despite the characteristic valerian root smell still being present, cat Z did respond to the *V. officinalis* root tincture, and this single response of nearly one and a half minutes was longer than 90% of all her responses to the plants tested. Furthermore, while cat Z also responded to the catnip and Tatarian honeysuckle tinctures, her response to the valerian root tincture was the longest. Although we only applied two sprays of each tincture, we still observed responses of all 5 cats 3.5 hours after we application (cats A and N to the *V. officinalis* tincture, cats O, V and Z to the *L. tatarica* tincture, and cat Z to the *N. cataria* tincture).

The results from this experiment suggest that at least some of the active compounds found in the cat- attracting plants can be effectively extracted simply by soaking the plant materials in absolute ethanol. Although cat Z did not respond to dried valerian root, she did respond to the tincture, suggesting compounds responsible for inhibiting her attraction were not coextracted with the active compounds. However, it is also possible that she preferred different amounts or ratios of the compounds in the tincture.

### Domestic cats respond to all iridoids, including dihydroactinidiolide, but response to actinidine is rare

We have previously shown that cats respond to cat-attracting plants known to contain little to no nepetalactone (Bol et al. 2017). While we detected iridomyrmecin, isodihydronepetalactone and actinidine in these plants, we did not confirm whether these compounds are responsible for the cat-attracting properties of these plants. The main goal of this experiment was to determine to which compounds identified in cat-attracting plants domestic cats would respond. Furthermore, we were interested to see if the differences in response between various cats to the individual cat-attracting plants (e.g., cat O responding significantly longer to *L. tatarica* than cat Z) and the differences in response of individual cats to the various plants (e.g. cat O responding significantly longer to *L. tatarica* than to *V. officinalis*) could be explained by different responses to the single compounds.

In these bioassays, performed with the same cats who also tested the plant materials, we tested not only the lactones *cis*-*trans*-nepetalactone (**1**), *trans*-*cis*-nepetalactone (**2**), isodihydronepetalactone (**4**), iridomyrmecin (**7**) and actinidine (**9**), but extended the repertoire by adding the lactones dihydronepetalactone (**3**), neonepetalactone (**5**), isoneonepetalactone (**6**), isoiridomyrmecin (**8**), the pyridine actinidine (**9**), the furanone dihydroactinidiolide (**10**), and indole (**11**) (**Figure 12, Table 3**). This selection (compounds **1** – **10**) was based on previous reports in the literature and summarized in the review by Arthur and Sharon Tucker (Tucker and Tucker 1988). We attempted to obtain or synthesize several other compounds mentioned in the work of Tucker and Tucker, such as boschniakine, but they were either not commercially available or unstable. In our hands, boschniakine was found to be particularly unstable when prepared through chemical synthesis. One hypothesis as to why cats respond to these molecules is that they resemble cat pheromones found in cat urine, feces, and glandular secretions. We identified indole as the only known compound in feline excretions that showed structural resemblance to the known cat-attracting compounds (Starkenmann et al. 2015, Miyazaki et al. 2018, Uetake et al. 2018) and therefore we also tested this compound as a cat-attractant. Thirty-three, 100, 300 and 900 µg of each compound was made available to the cats on two different days, for a total of at least 17 hours per compound.

**Figure 12.**
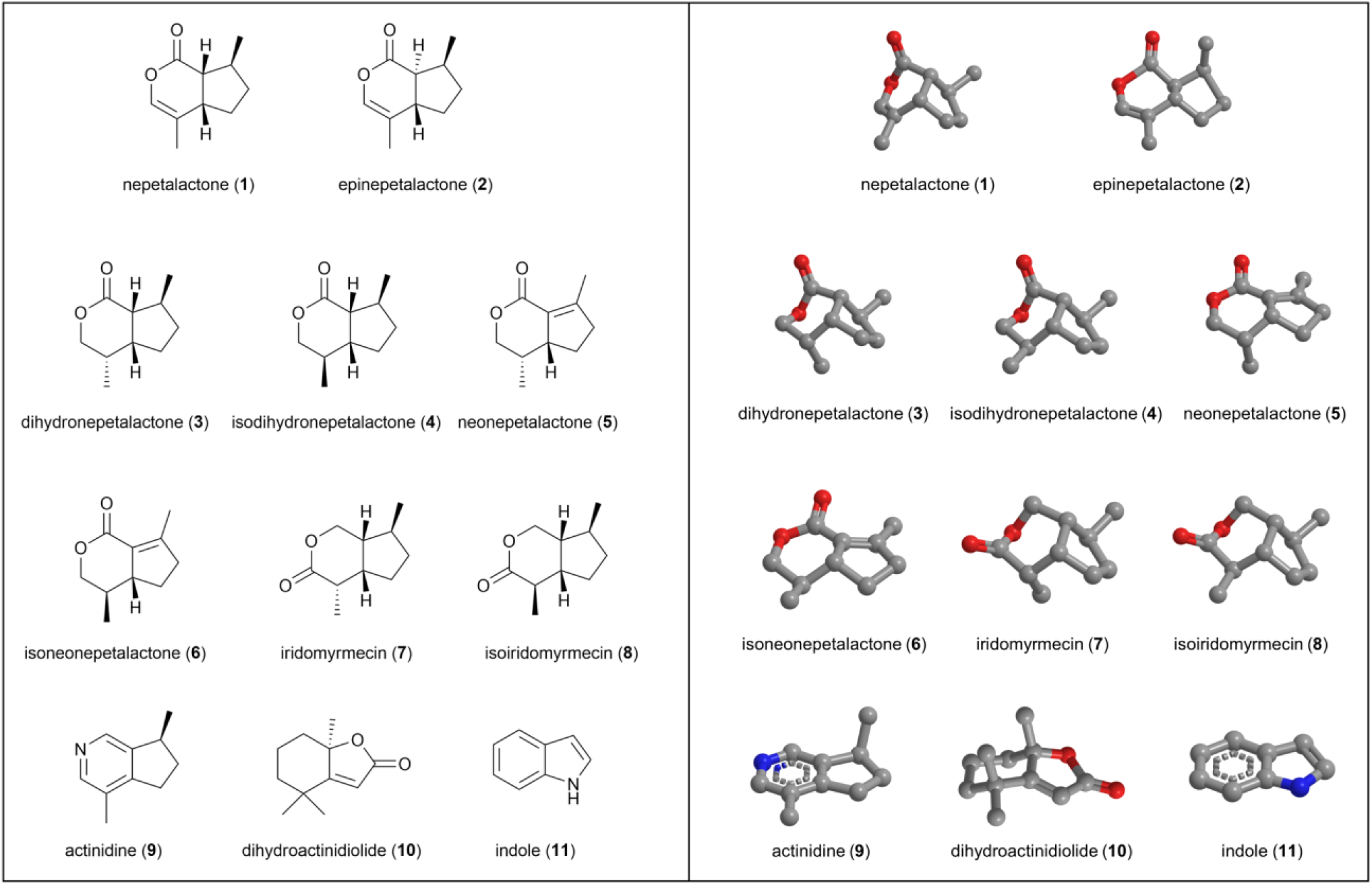
Structures of the single compounds used for bioassays with domestic cats. Two dimensional structures are shown on the left, 3D structures are shown on the right. Oxygen atoms are shown in red, nitrogen in blue. Nepetalactone (**1**) and epinepetalactone (**2**) are also referred to as *cis*- *trans*-nepetalactone and *trans*-*cis*-nepetalactone, respectively. Note how the location of the carbonyl group is different between the type I lactones **7** – **8** and the type II lactones **1** – **6**.

We found that all of the plant-derived compounds (**1 – 10**) elicited a positive response in domestic cats, but not the negative control (evaporated diethyl ether) nor indole (**Figure 13A**, **Supplementary Figure 7**). All responses could be classified as “catnip responses”. There was no statistically significant difference in median response duration of the 5 cats between the active compounds (P > 0.05, Friedman test). The response time among cats to actinidine had a larger range and more uneven distribution than any of the other compounds (**Figure 13A**). Three out of the 5 cats showed no or little interest in this compound.

**Figure 13.**
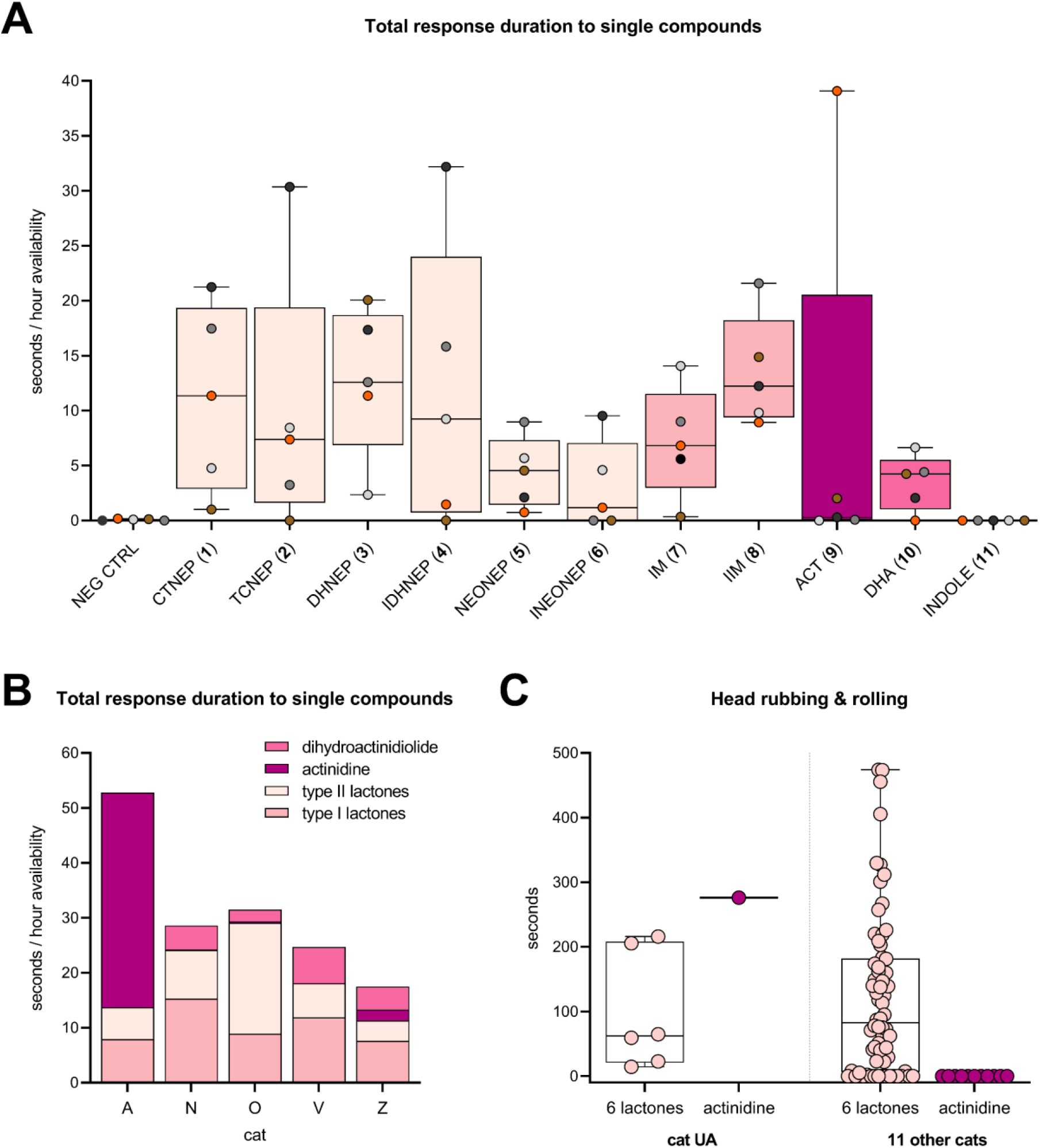
Response time of domestic cats to single compounds. (**A**) Response time, shown as seconds per hour each compound was available, per compound. Note the large range and uneven distribution of the data for actinidine. Each compound was available for at least two days; 5 hours on the first day and 12 hours on the subsequent test day. Negative controls (fabric with evaporated diethyl ether) were always tested alongside the single compounds. (**B**) Response time to single compounds, grouped by their chemical structure, shown per cat. For a cat responding equally long to every class of compounds (assuming there is no variation in response to the compounds within each group), one would see equal heights for each of the 4 portions of the bar. Type I and II lactones were available for 34 and 120.5 hours, respectively. Actinidine was tested for 53 hours on 5 days and dihydroactinidiolide was available for the cats for a total of 17 hours (2 days). (**C**) Duration of head rubbing and rolling of 12 domestic cats in response to iridoids. The data plotted here was obtained from the supplementary online material recently published by Uenoyama *et al*. (Uenoyama et al. 2021). The authors did not analyze or discuss these data in their article. The name of the only cat responding to actinidine in the study of Uenoyama *et al*. coincidentally is also cat A and is not the same cat as cat A in our study. To avoid confusion, we renamed this cat UA.

Therefore, we tested actinidine on three additional days. All actinidine data shown is from 5 days of testing, between January and May 2019, totaling 53 hours of exposure (**Supplementary Figure 7**). We also made fabric with a higher amount of actinidine (2700 µg) available for 4 hours to compensate for potential variation between the cats in their detection threshold for this single compound. The three cats who did not respond to actinidine were cats O, N and V. Interestingly, these cats had the longest total response time to all other compounds (**Figure 13B**). While most cats did not respond to actinidine, cat A responded longer to actinidine than to any of the other compounds that were tested (**Figure 13B**). The response duration of cat A to actinidine was 5 – 9 × longer than her response to the lactones. These data did not provide information on how common the response to actinidine is among domestic cats, especially since the three non-responders are suspected to be genetically related. However, recently published supplementary data by Reiko Uenoyama and her colleagues that was not analyzed or discussed in their article (Uenoyama et al. 2021) strongly suggest that a response to actinidine is less common given only one of 12 cats in their study responded to actinidine (**Figure 13C**). Furthermore, all 11 other cats who did not respond to actinidine responded to most (approximately 5 out of 6) of the lactones that were also tested (**Figure 13C**). These results from the study of Uenoyama *et al*. reinforce our findings. Uenoyama *et al*. tested 50 µg of the single compounds. Since we observed more than half of the total response time of cat A to actinidine when we used 33 and 100 µg, it is unlikely that the absence of a response of those 11 cats would be due to the amount of actinidine used in their experiments.

The longer response time of cat A to actinidine compared to the lactones could be explained by both an increased response frequency and duration of the individual responses. Cat A responded to actinidine once every 1.75 hours, compared to roughly once every 6 hours for the lactones, which is almost 3.5 × more frequent. The median response duration to actinidine of cat A was statistically significantly longer than to the lactones (42 and 18 seconds, respectively; **Supplementary Figure 8**).

For the analyses described above, data were pooled from tests with various quantities of the single compounds (33, 100, 300, 900 and for actinidine even 2700 µg) performed on different times of the day (morning, afternoon, evening). We used data from the compounds for which we observed at least 10 responses of an individual cat to look for possible correlation between quantity of the compound and response duration/frequency. Cat A responded 30 times to actinidine (**9**), and cat O responded 14, 10 and 10 times to compounds (**2**), (**3**) and (**4**), respectively. The data show absence of a dose-response relationship at quantities ranging from 33 to 2700 µg (**Supplementary Figure 9A-B**). Furthermore, we found that the distribution of responses matched the distribution of the hours the olfactory test objects were available to the cats through the day (**Supplementary Figure 9C**). This result indicates the cats were not less active in the afternoon, which may have resulted in fewer responses during this part of the day. Taken together, these data suggest that pooling data (different quantities and tests performed at different times of the day) did not affect the results and conclusions.

When we compared the cats’ response duration to the plants with the response duration to the single compounds, we found a very strong positive correlation (**Figure 14**). The response duration to the cat- attracting plants was approximately 33% longer than to the single compounds. This might be explained by higher quantities of compounds in the plants, the presence of multiple compounds, slower and more sustained release of compounds, larger volume of the test object, or a combination of these.

**Figure 14.**
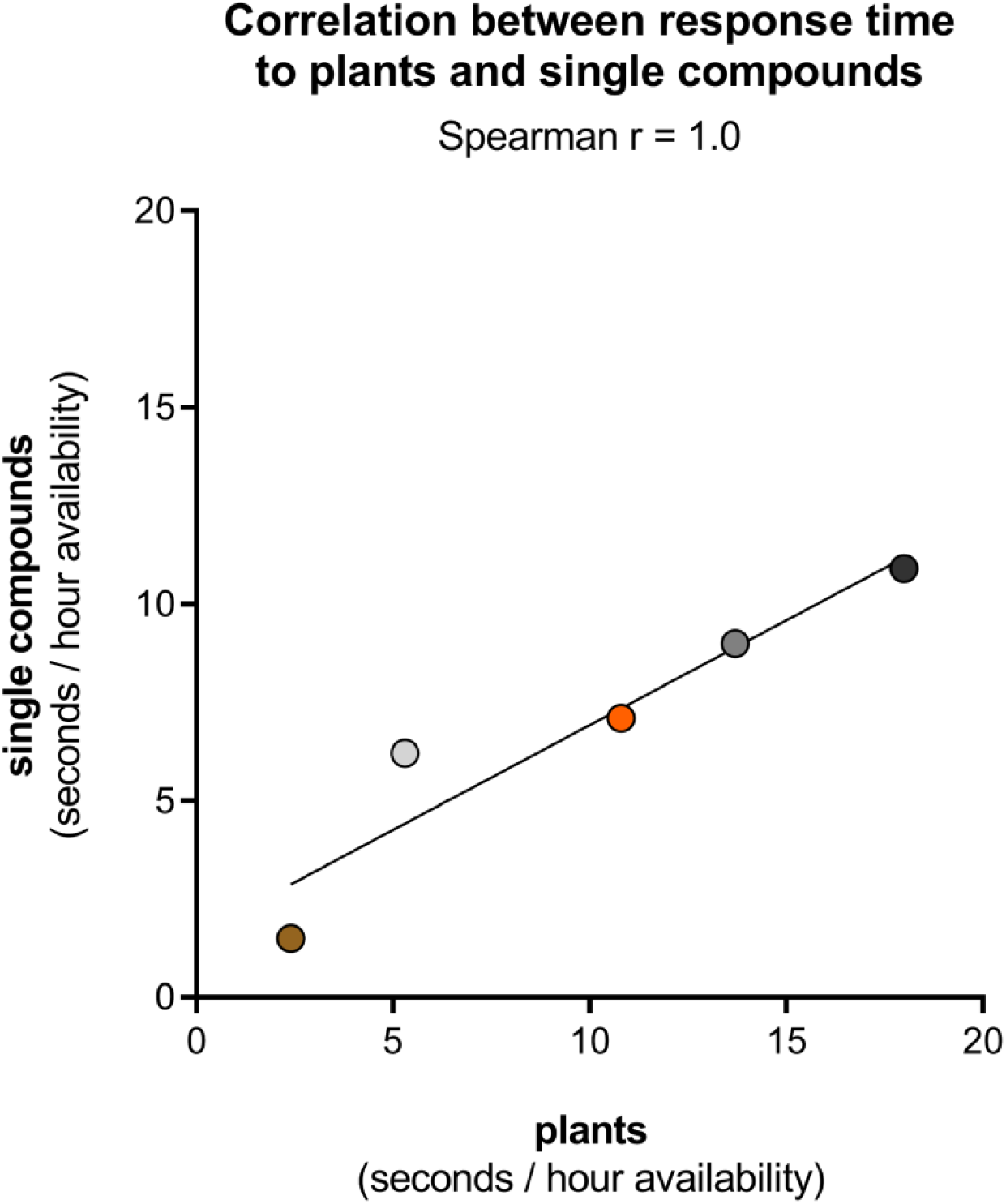
Correlation between response duration to cat-attracting plants and single compounds. For each cat the median of the 5 response times to the 5 cat-attracting plants (X axis) and the median of the 10 response times to 10 single compounds (**1** – **10**) (Y axis) are shown.

### The degree of attraction to the single compounds differs between cats

Similar to what we observed for the plants (**Figure 3B**), we found that the time to first response to the single compounds was significantly different between cats (**Figure 15**). When we looked at the data for each cat separately, we also found significant differences in time to first response between the different classes of single compounds (lactones, actinidine, dihydroactinidiolide). As expected, cat A was significantly more attracted to actinidine than to the lactones or dihydroactinidiolide, whereas the opposite was seen for cat V. The time to first response to actinidine of Cats N and O was also longer compared to the lactones, but the difference was not statistically significant because of an outlier. The responses of cats N (n=1) and O (n=2) to actinidine lasted only a few seconds and might be considered ‘false positives” (see below).

**Figure 15.**
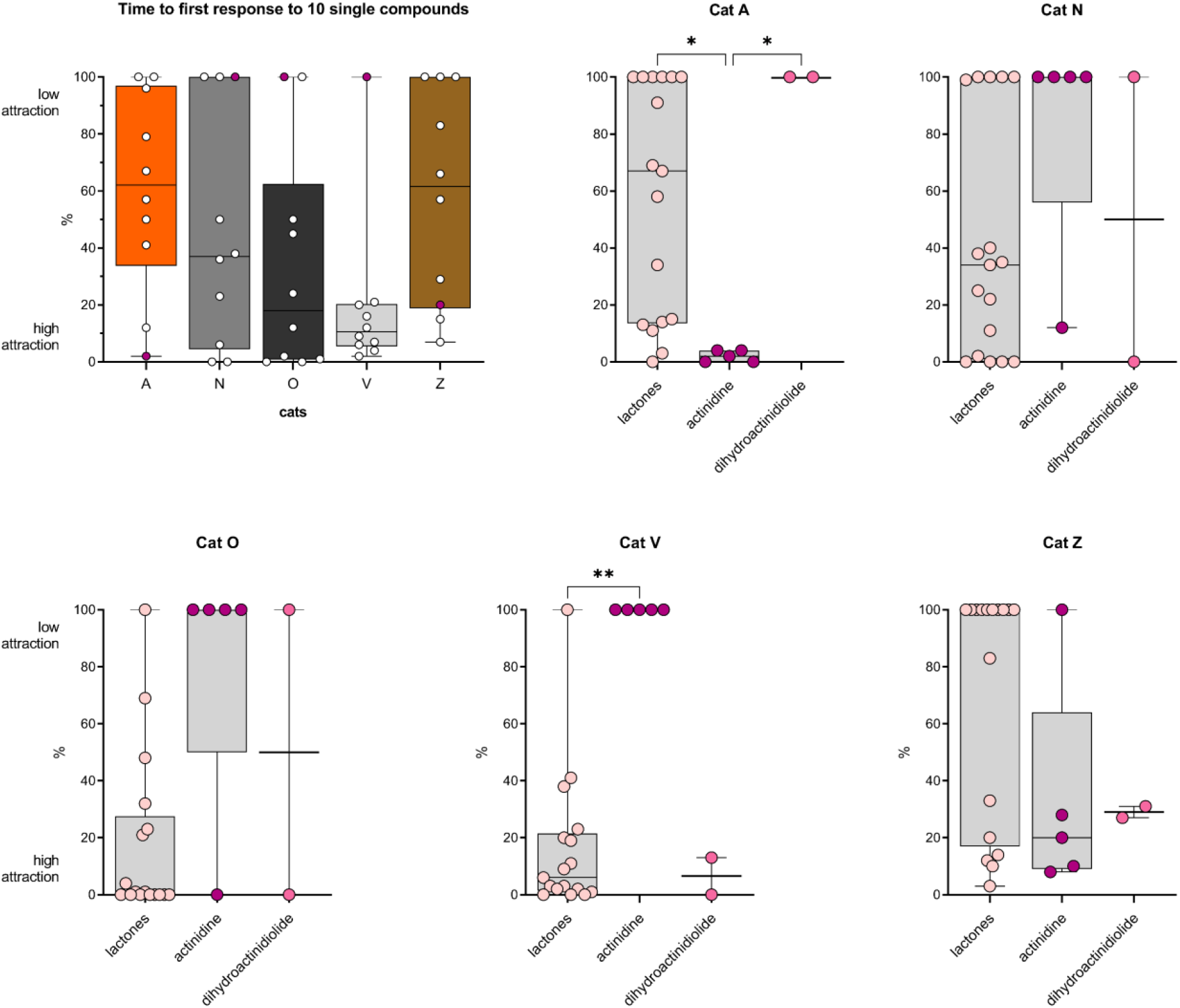
Time to first response of 5 domestic cats to single cat-attracting compounds. The time to first response was determined for every cat, for every day that a single compound (**1** – **10**) was tested (n=24). When a cat did not respond to a compound on a test day, the time the stimulus was available that day was used as time to first response. Since the compounds were available for different durations, typically 5 and 12 hours, the time to first response was expressed as a percentage of the time the compound was available, with 0% being an immediate response and 100% no response at all that day. For each compound (10 per cat) the median percentage is shown. The second test day of neonepetalactone was not included because the recording stopped about 40 minutes after the start of the experiment. The differences in time to first response between the 5 cats was statistically significant (P < 0.05, Friedman test). In addition, the differences in time to first response between actinidine and other compounds for cat A, as well as the difference between the lactones and actinidine for cat V were statistically significant (Kruskal-Wallis test). P values shown in the figure are from Dunn’s post-hoc test. * P < 0.05; ** P < 0.01

These findings support the previous observation that there is variation between cats in how attracted they are to certain cat-attracting scents. These data also strengthen the hypothesis that actinidine is distinct, not only in structure, but also in the effect it elicits in domestic cats. The near immediate (seconds after it was made available) “response” from cat O to actinidine supports the hypothesis that the time to first response is at least in part determined by the cat’s personality (i.e., curiosity, fear of missing out).

### Behavioral response to actinidine is different from responses to lactones and cat- attracting plants

Next, we analyzed the behavior of cat A using BORIS software to determine if there was a difference in her behavior when exposed to plants, lactones and actinidine. Since the responses of cat A to the various plants (n=5) was highly similar (**Figure 7A**), we only used the *N. cataria* data for the comparison to the single compounds. For the plants, five responses nearest to one minute were analyzed. To keep the median response time similar, we only analyzed responses of cat A to the lactones and actinidine with a duration between 30 – 90 seconds (n=9 and n=16, respectively). Interestingly, cat A spent significantly more time licking the object with actinidine and less time head rubbing, when compared to the responses to the lactones or *N. cataria* (**Figure 16** and **Figure 17**). The same statistically significant differences were seen when all responses to actinidine and the lactones longer than 30 seconds were analyzed (n=11 and n=24, respectively), capturing 95% and 83% of the total response duration to these compounds, respectively. The percentage head rubbing was lower for actinidine as the result of more time spent licking. Other than a difference in the frequency of head shaking, no differences were seen in any of the other behaviors.

**Figure 16.**
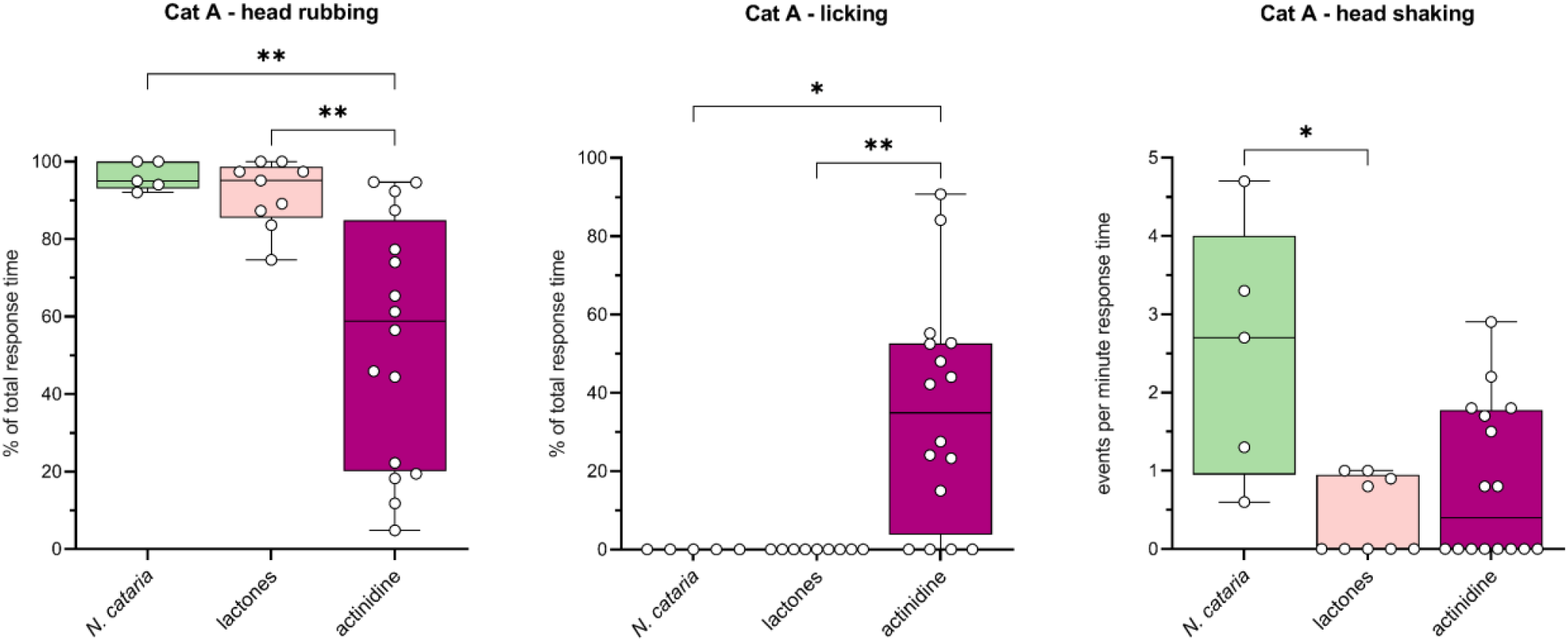
Differences in behavior of cat A between responses to actinidine, lactones and *N. cataria*. Nine responses to the lactones and 16 responses to actinidine with a response duration 30 – 90 seconds were analyzed using BORIS behavioral analysis software. Results were compared to the behavior seen in response to catnip (Figure 7A). The Kruskal-Wallis test was used to test for differences. P values shown are from Dunn’s post-hoc test. * P < 0.05; ** P < 0.01

**Figure 17.**
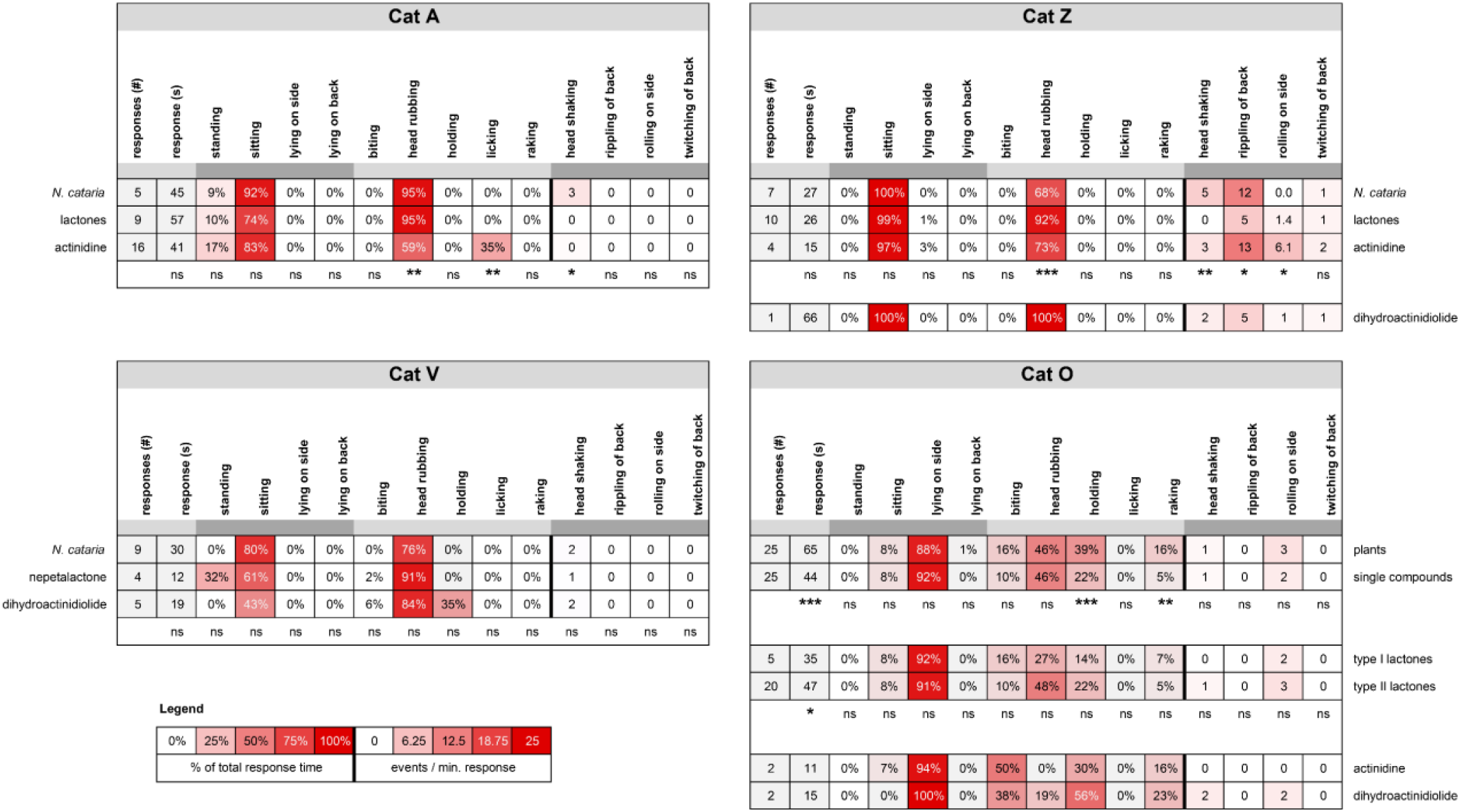
Heatmaps showing similarities and differences in body position and behaviors of 4 cats in response to cat-attracting plants and single compounds. Not all cats responded to all classes of single compounds and therefore comparisons differ between cats. Responses to actinidine and dihydroactinidiolide of cat O and to dihydroactinidiolide of cat Z are shown but were not included in the statistical analysis because the number of responses were two or less. Unless otherwise indicated, numbers represent the median. The Kruskal-Wallis test followed by Dunn’s post-hoc test or the Mann Whitney test was done to test for statistically significant differences. #, frequency; s, seconds; ns, not statistically significantly different; * P < 0.05, ** P < 0.01, *** P < 0.001

It seems that the observed licking of cat A is a true feature of her response to actinidine and not the result of longer response durations that we have seen for actinidine compared to the lactones (**Supplementary Figure 8**). Indeed, we found no correlation between the percentage of response time licking and response duration (**Supplementary Figure 10A**). Although licking was the dominant behavior observed for the two responses to the fabric with the highest amount of actinidine (2700 µg), the correlation between the amount of actinidine and the percentage of response time spent licking was weak (**Supplementary Figure 10B**).

Cat Z also responded to actinidine, but the responses were much less frequent and shorter in duration compared to cat A. Three short responses (10 – 20 seconds) and one response of almost one minute were observed. While active engagement (contact) with the object was a requirement for any feline activity to be considered a response, about 90% of the time that cat Z responded to actinidine she did not touch the object. This lack of contact during the response was also seen for freshly harvested, locally-grown *A. polygama* leaves, plant material known to contain relatively large amounts of actinidine (Bol et al. 2017). However, the response occurred in close proximity to the test object and her behavior was characteristic of what was seen with the other plants and compounds: head rubbing (the floor near the object) in a sitting position, rippling and twitching of her back and occasionally rolling on her side. Only cat Z demonstrated responses without touching the olfactory object. Since cat Z did not respond to any of the negative controls that were available for hundreds of hours, and given her most recent response to any olfactory stimulus prior to actinidine was 3 months earlier, we believe this response was specific.

The median response duration to the lactones (n=10) and actinidine (n=4) of cat Z was 26 and 15 seconds, respectively. Therefore, we included her two shorter responses to *N. cataria* in the qualitative and statistical analysis. As a result, we compared all her responses to catnip (n=7), all responses to actinidine and all responses < 60 seconds to the lactones. We also observed some differences between her responses to catnip, the lactones, and actinidine. It appeared that the response of cat Z to actinidine was more dynamic. Cat Z rolled on her side more frequently in response to actinidine than in response to *N. cataria* or the lactones (**Figure 17** and **Supplementary Figure 11**). Rippling of the back was seen less in response to the lactones as compared to catnip and actinidine, and for this reason the contribution of head rubbing to the total response duration of the response increased. Head shaking was also seen less frequently during responses to the lactones compared to catnip. When the 12 responses of cat Z to the other plants (*A. indica*, *A. polygama* and *L. tatarica*) (**Figure 7B**) were included in the statistical analysis, the results remained unaffected, except that the difference in the frequency of rippling of the back between actinidine and the lactones also became statistically significant (P < 0.05; data not shown).

The response of cat O to actinidine was uncharacteristic for him and did not resemble the “catnip response”. Both of his extremely short responses to actinidine (each about 10 seconds) lacked rubbing of the object, which was seen in all his responses to the plants and lactones (**Figure 17**). While the behavior of cat O to type I and II lactones (this discrimination is made based on the position of the carbonyl group; see **Table 3** and **Figure 12**) was near identical, less holding and raking of the object was seen for the lactones compared to the 15 grams of plant material (**Figure 17**), possibly due to lack of volume of the object.

Collectively, these data suggest that while the responses to the single compounds are in general similar to the behavior seen in response to the cat-attracting plants, there appear to be biologically significant differences between actinidine and the lactones.

### Behavioral response to dihydroactinidiolide is similar to behavior in response to lactones

Another molecule that is structurally different from type I and II lactones (as well as actinidine) is dihydroactinidiolide (**Figure 12**), which contains a furanone ring (5 membered lactone) compared to pyranone rings (6 membered lactone). Interestingly, unlike the compounds **1** – **9** tested in this study that have only been detected in plants or insects, dihydroactinidiolide has additionally been detected in glandular secretions and urine of the red fox (McLean, Nichols and Davies 2021, Albone 1975, McLean, Davies and Nichols 2017). None of the other iridoids tested here are produced or secreted by a mammal to our knowledge. We wanted to determine if the behavior of cats triggered by this compound was similar to the behavior seen in response to the cat-attracting plants and the other single compounds. Four out of 5 cats responded to this compound; however, the number and duration of the responses were low (13 responses in total for all 4 cats with a median response duration of 20 seconds) (**Figure 13A**). Of the cats exposed to dihydroactinidiolide, cat V responded most frequently (n=5) and therefore the behavior she demonstrated during those 5 interactions was analyzed using BORIS to compare to her behavior to *cis-trans* nepetalactone and *N. cataria*. The behavior seen in response to *N. cataria* and nepetalactone – sitting and head rubbing the object, holding the object while on her side, raking, and biting it, rolling on her side, and shaking her head – was also observed for dihydroactinidiolide (**Figure 17** and **Figure 18**). Head rubbing was again the dominant behavior, making up about 85% of the response time. There were no significant differences in behavior between catnip, nepetalactone and dihydroactinidiolide.

**Figure 18.**
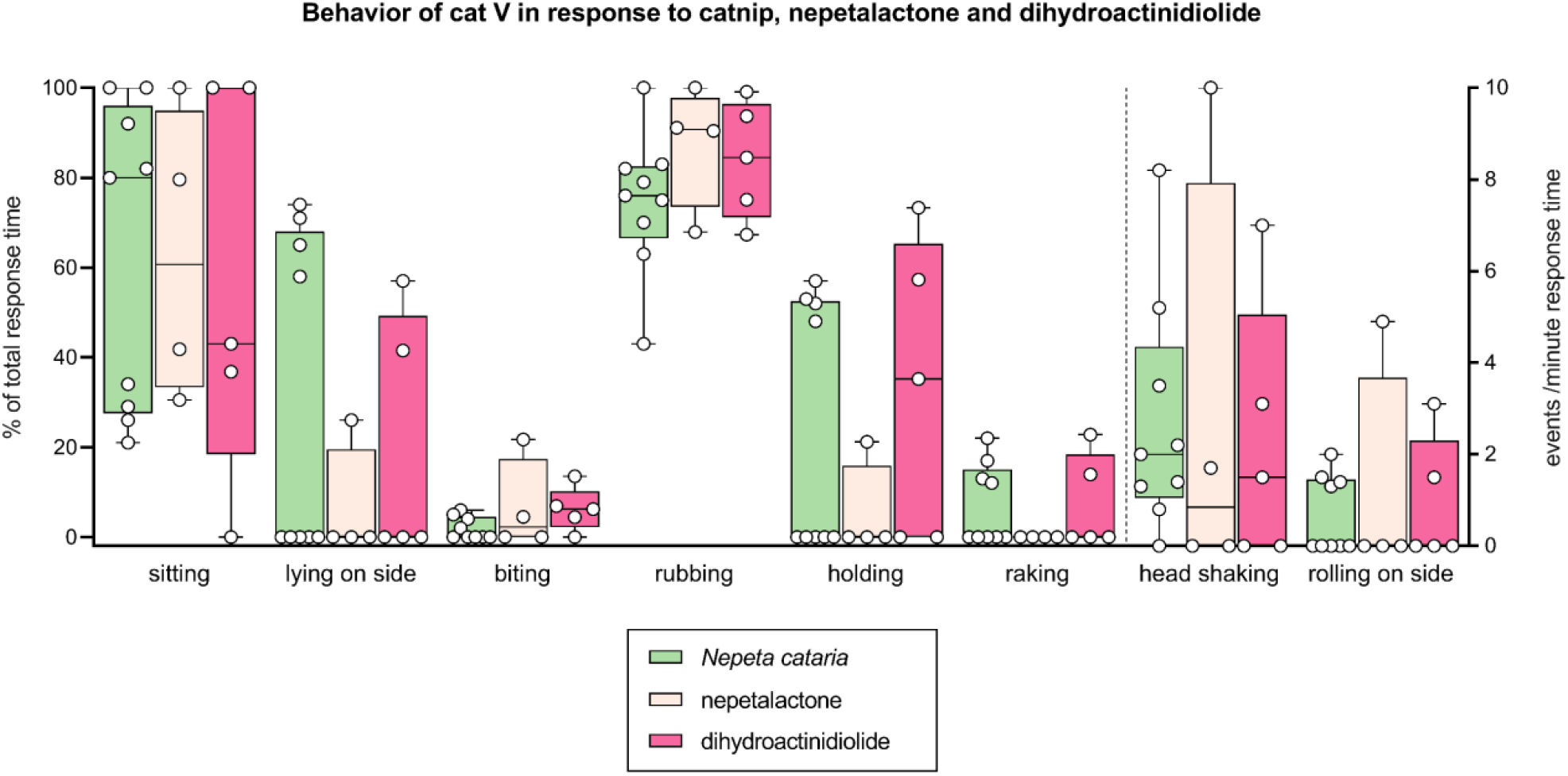
Comparison of behavior between responses to *N. cataria*, nepetalactone and dihydroactinidiolide. Results from behavioral analysis in BORIS of responses of cat V to *N. cataria* (n=9), *cis*-*trans*-nepetalactone (n=4) and dihydroactinidiolide (n=5) are shown. Some of the responses were short and this may have contributed to some outliers. Head shaking and rolling on the side are plotted on the right Y axis. There were no significant differences in behavior between catnip, nepetalactone and dihydroactinidiolide (Kruskal-Wallis and Dunn’s post-hoc test).

Cats N and O both responded only twice to dihydroactinidiolide and therefore we did not perform statistical analysis to test for differences. Responses of cat N to the plants were typically in a sternal or lateral position and included mostly head rubbing (60 – 80% of response duration), sometimes while holding the object. She also rolled on her side or back, about 2 – 3 times per minute of response duration and rippling of the back was also seen, about 3 times per minute. When we compared this with her behavior in response to dihydroactinidiolide, we noticed that head rubbing was still the most dominant behavior (about 80% of the response time) with rippling of the back making up the majority of the balance (about 2 times per minute). However, no holding of the object and no rolling on the side, and hence no response in a lateral position, were observed (data not shown; **Supplementary File 3**).

The two responses of cat O to dihydroactinidiolide were short (10 and 20 seconds), but resembled his responses to the plants: head rubbing, biting, holding the object, and raking were all seen, while in lateral position (**Figure 17**).

The behavior of cat Z in response to dihydroactinidiolide matched her typical behavior when exposed to plants and single compounds. She responded in a sitting position, head rubbing the object while her back rippled. Since we observed only a single response from her, we could not test for statistical differences between dihydroactinidiolide or other cat-attracting plants or compounds. However, the behavior of her 66 seconds response was near identical to the behavior seen during her responses to the lactones (**Figure 17**).

While the number of observations were limited and the duration of the responses was often short, we believe these data show that the behavioral response to the structurally distinct dihydroactinidiolide is highly comparable to the behavior seen in response to the other single compounds and the cat-attracting plants.

### Stability of the single compounds

We chose diethyl ether as solvent because of its inert nature and volatility, meaning it would evaporate quickly and leave only pure compounds behind. The compounds were tested immediately after they were dissolved in diethyl ether because information about their stability is lacking. Some of the *trans*-*cis*- nepetalactone, neonepetalactone, isoneonepetalactone and actinidine dissolved in diethyl ether was stored between experiments for a variety of reasons, as explained in detail in the “Materials and methods” section. The results obtained with these compounds gave us some insight into the stability of these compounds in diethyl ether under various conditions. When comparing the results between compounds that were used immediately after dissolving in diethyl ether and those that were stored after dissolving, we did not find any clear evidence of reduced activity, suggesting they were stable. When *trans*-*cis*- nepetalactone was tested on two additional days, after being stored at room temperature for 1.5 months, both the response frequency and total response time for all cats combined was higher on these days (C and D) (18 responses, 17.6 minutes) compared to days A and B (9 responses, 10.5 minutes). We also did not observe reduced response duration to neonepetalactone on day B after the dissolved compound had been stored at 4°C for 4 days. 75% of the responses to neonepetalactone occurred on day B, whereas this was 50 – 85% for the other lactones. While dissolved actinidine was stored for two weeks at various temperatures ranging from freezing to room temperature, we still observed minutes of response to this compound, albeit only during the first 4 hours of a 15-hour testing day (**Supplementary Figure 7**, actinidine, day C). The absence of responses in the afternoon and evening were in contrast with what was observed on days when actinidine was used immediately after dissolving in diethyl ether (days B and E). Any possible degradation of actinidine would not affect the conclusions drawn in this manuscript since this only would underestimate the true response of cat A.

### Any cat-attracting property of (pepper)mint is not caused by structural resemblance of the active compound(s) to molecules like nepetalactone

In addition to *Nepeta cataria*, there are several other plants from the genus *Nepeta* that contain cat- attracting type I and II lactones (Formisano, Rigano and Senatore 2011, Regnier, Waller and Eisenbraun 1967, Bicchi, Mashaly and Sandra 1984, Eisenbraun et al. 1980), of which *Nepeta mussinii* or catmint is arguably the best-known. Although all these plants are members of the Lamiaceae family, commonly referred to as the mint family, plants in the *Nepeta* genus are not closely related to plants in the *Mentha* genus, such as peppermint. There are numerous anecdotes of cats being attracted to peppermint (*Mentha piperita*) and topical analgesics such as Bengay, IcyHot and Vicks VapoRub (which should all be kept away from cats). Interestingly, *L. tatarica* (Tatarian honeysuckle) wood has a minty smell. Therefore, we studied how domestic cats respond to the odiferous molecules menthol and methyl salicylate that are responsible for the characteristic mint fragrance. Fabrics containing 33, 100, 300 and 900 µg menthol or methyl salicylate were tested separately, and each was available on two different days for a total of 17 hours (5 hours on the first day, 12 hours on the second day). None of the cats responded to either of the two compounds.

### Fragrances

Anecdotal evidence from the past decade suggests that big cats (cheetahs and cats of the *Panthera* genus: lion, tiger, leopard, snow leopard and jaguar) respond to certain fragrances (e.g. perfume, eau de toilette), Calvin Klein’s Obsession for Men in particular, in similar fashion to catnip (Banham Zoo, Norfolk, England: Time 2020, BBC 2020 and The Washington Post 2020; Taronga Zoo, Sidney, Australia: Scientific American 2014; Brookfield Zoo, Chicago, IL: CBS 2010; Bronx Zoo, New York, NY: Wall Street Journal 2010 and National Geographic 2010). Patrick Thomas and his colleagues published the results of a scent study in the Bronx Zoo where the responses of two adult cheetahs to 24 different fragrances were studied (Thomas et al. 2005). The researchers applied three sprays of each fragrance to an object in their 1,000 m^2^ outdoor naturalistic enclosure on three different days and reported the mean latency to inspect the scent, the mean number of visits to the scent, the mean contact time to the scent and if head rubbing was observed. All but one of the 24 fragrances were investigated by at least one of the two cheetahs, demonstrating how scents can be used for environmental enrichment. However, head rubbing was only seen in response to seven fragrances, and the median contact time to these was significantly higher than the contact time to the other fragrances.

We were interested to see if the response of domestic cats to fragrances is similar to the response of big cats. Furthermore, since many fragrances contain essential oils obtained from plants, we wondered if the presence of compounds such as nepetalactone, iridomyrmecin, actinidine or dihydroactinidiolide in the fragrances could be responsible for the observed behavior of the cats. It is known that lions, jaguars, leopards, snow leopards and bobcats respond to plant material containing these compounds (*N. cataria* or catnip and *A. polygama* or silver vine) (Todd 1963, Bol et al. 2017). If these compounds are present in perfumes, colognes or eau de toilettes, then we would expect domestic cats who respond to plant materials containing these stimulants to also be attracted to these fragrances. To test this hypothesis, we selected the four most popular fragrances (head rubbing by both cheetahs and longest average contact time: 668, 662, 207 and 185 seconds) of the 24 used in the study by Thomas and colleagues. We applied them to a polyester fabric and made the fragrances available to domestic cats A, N, O, V and Z who all responded to most or all of the tested plants and single compounds. The fabrics were sprayed twice (approximately 200 µL) with either Obsession for Men, L’Air Du Temps, Paco Rabanne Pour Homme, Drakkar Noir, or ethanol as a negative control (**Table 5** and **Table 6**). The fabrics were then made available to the cats for 15 hours, from 7:00 till 22:00 upon which all cats investigated the fabrics several times. Cat A responded to the Drakkar Noir for a duration of 3 minutes and 50 seconds (**Figure 19A**) and this response resembled the behavior seen in her responses to the cat-attracting plants and single compounds: head rubbing and licking of the fabric with the fragrance while in a sitting position, shaking of the head and occasional twitching of the back (**Figure 19B**). None of the other cats responded to any of the other three fragrances tested, including the popular (among big cats) Calvin Klein’s Obsession for Men. Therefore, we tested Obsession for Men a second time, one week later, again for 15 hours, and used a bottle obtained from a different source. Drakkar Noir was also made available for a second time. Again, only cat A responded to Drakkar Noir, this time for 3 minutes and 40 seconds (**Figure 19A**), while none of the cats responded to the fabric with Obsession for Men. In both cases, the response of cat A to Drakkar Noir occurred approximately 14 hours after the fabric was made available. After this amount of time in a well ventilated, open area, it is expected that only larger, less volatile molecules will remain on the fabric, such as nepetalactone, iridomyrmecin, actinidine and dihydroactinidiolide that are found in the essential oils of cat-attracting plants. In addition to base notes (larger, less volatile molecules), fragrances have what is referred to in the fragrance industry as top and middle (heart) notes consisting of molecules that can be detected more quickly. Some of these smaller, more volatile compounds may interfere with the detection or perception of other, potentially cat-attracting molecules. To increase the exposure time to the larger single compounds only, fabrics sprayed about 10 times were left to stand overnight at room temperature (at a location where the cats could not smell them) before making them available to the cats. While cat O briefly (a few seconds) interacted with the fabric sprayed with Obsession for Men, his behavior was contrary to his responses to the plants and single compounds (sitting, no head rubbing and no raking). This alternative methodology did not lead to any other responses of the cats to the fragrances. To decrease the chance that the lack of response to Calvin Klein’s Obsession for Men were false negative results, a third source of this fragrance was made available to the cats a fourth and fifth time, but no responses were observed.

**Figure 19.**
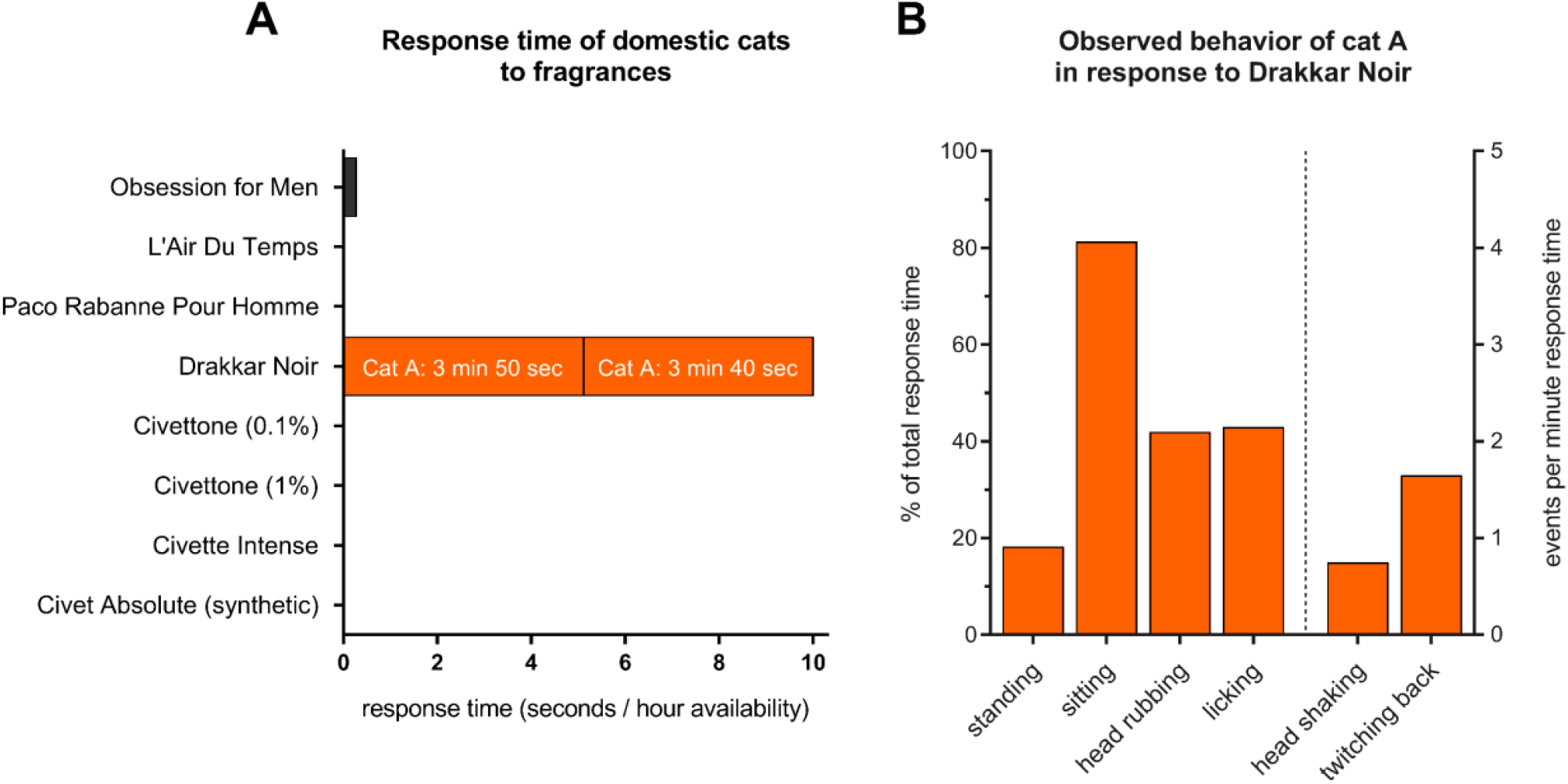
Responses of five domestic cats (*Felis catus*) to fragrances attractive to *Panthera* (jaguar, leopard, snow leopard, lion and tiger) and *Acinonyx jubatus* (cheetah). (A) Response duration plotted as time per hour the fragrances were available to the cats, with the response time of two responses by cat A to Drakkar Noir shown within the bar. Obsession for Men was available for 75 hours (5 days), Drakkar Noir 45 hours (3 days), L’Air Du Temps and Paco Rabanne 30 hours (2 days), and the other fragrances for 15 hours (1 day). (B) Analysis of the observed behavior of cat A in response to Drakkar Noir. The average of the two responses is shown. Body position, head rubbing, and licking are shown as the percentage of the response duration, while head shaking and twitching of the back are shown as events per minute of response and are plotted on the right Y axis.

Many have speculated that the response of cats to Calvin Klein’s Obsession for Men is due to the presence of the molecule civetone (Mandy Aftel, The Washington Post 2020 and NPR 2018; Miguel Ordeñana, Scientific American 2013; Ann Gottlieb, Wall Street Journal 2010). Civetone is an odiferous ketone found in civet, a glandular secretion of the civet cat (Anonis 1997) but it can also be synthesized (Tanabe 2002). To test if domestic cats would respond to civetone or civet we sprayed 0.1 and 1% civetone (a kind gift from Fred Keifer at Firmenich), a fragrance known to contain civetone (Civette Intense), and 1% absolute civet (a synthetic recreation of natural civet) on a fabric and made each available to the domestic cats for 15 hours (**Table 5** and **Table 6**). For ethical reasons, we decided not to obtain and test natural civet. None of the five cats showed any interest in the fabrics containing these scents (**Figure 19A**).

These data suggest that domestic cats do not respond to fragrances like big cats do and that the response of big cats to fragrances such as Obsession for Men is unlikely triggered by the presence of compounds similar to nepetalactone, iridomyrmecin, actinidine or dihydroactinidiolide. Indeed, GC-MS analysis of Obsession for Men revealed that no cat-attracting single compounds were detected in this fragrance. The similar negative result for Drakkar Noir suggests the presence of (an) other, unidentified cat-attracting compound(s) in this fragrance.

While domestic cats do not seem to respond to civetone, this conclusion does not exclude the possibility that the big cats do, since the fragrances that were highly popular among cheetahs were not very popular among the domestic cats.

### Cat-attracting plants contain a wide array of nepetalactone-like molecules

Previously, we quantified 5 cat-attracting molecules (nepetalactone, epinepetalactone, isodihydronepetalactone, iridomyrmecin and actinidine) in the plant materials that we used in our preceding study (Bol et al. 2017) using tridecyl acetate as an internal standard. Here, we were able to use the synthesized single compounds as standards, which were previously not available. Therefore, we were able to quantitate these compounds more accurately and quantitate additional compounds in the plant tissues. We again analyzed catnip leaves, silver vine fruit gall, Tatarian honeysuckle wood, and valerian root. We now also included Indian nettle root, Texas-grown silver vine leaves, and lignified silver vine stem (**Table 2**). All these plant tissues were from the same batches that were used in the experiments described in this article. We also performed GC-MS analysis on samples from inside the socks that were used for testing. We did not find any evidence that would suggest significant loss of active compounds as the result of interactions with the cats (e.g., contact with saliva) over the duration of the experiments (**Supplementary Figure 12**).

As expected, nepetalactone and a 5 – 10-fold lower amount of epinepetalactone were only detected in the *Nepeta cataria* samples (**Figure 20**). In addition to these two compounds, nearly all other known cat- attracting compounds were detected in catnip, except for neonepetalactone and isoneonepetalactone.

**Figure 20.**
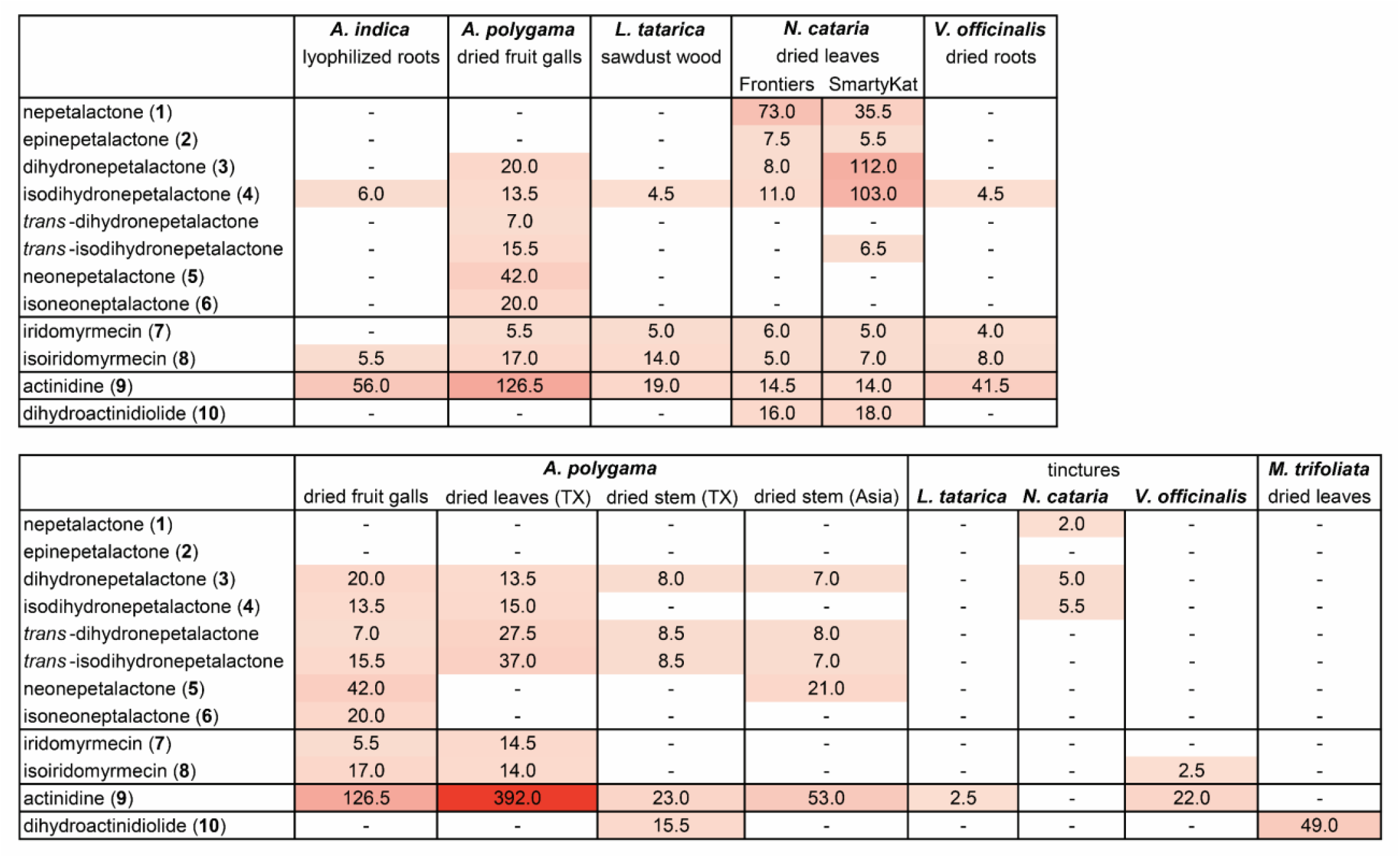
Quantitation of cat-attracting compounds in plants using GC-MS. The plant tissues used for this analysis were fresh samples. They were taken from the same bags of plant material that were used for the 10 × 10-hour testing. Amounts are reported as µg per gram plant material, except for the tinctures (µg/mL tincture). Tinctures were made by adding 5 volumes ethanol (500 ml) to one volume of plant tissue (10, 20 and 50 grams for catnip, Tatarian honeysuckle and valerian root, respectively). Dashes indicate that the compound was not detected. Numbers are rounded to the nearest half. Reported values are the average of three separate extractions of the plant material. Unrounded numbers with standard error of the mean are shown in **Supplementary** Figure 12. Where compounds (**3**) and (**4**) are reduced forms of compound (**1**), *trans*-dihydronepetalactone and *trans*-isodihydronepetalactone are reduced forms of compound (**2**). *Trans*-dihydronepetalactone and *trans*-isodihydronepetalactone were not used in the bioassays with domestic cats.

We found that one of the two brands of catnip contained large amounts of dihydronepetalactone and isodihydronepetalactone (which are reduced form of nepetalactone). However, surprisingly, this difference of about one log did not result in an increased response time of the cats (**Supplementary Figure 4**). In fact, the opposite was observed for cat O: a significantly longer response time was seen for Frontier catnip. *A. polygama* fruit galls also contained a large number of active compounds (n=9), including *trans*-dihydronepetalactone and *trans*-isodihydronepetalactone, which are reduced forms of epinepetalactone. The amount of actinidine in the silver vine fruit galls was about one log more than what was extracted from the catnip leaves. The fruit galls contained neonepetalactone and isoneonepetalactone, which were absent in catnip, but did not contain dihydroactinidiolide. The latter was only found in catnip. Surprisingly, the chemical composition of Tatarian honeysuckle wood was highly similar to that of valerian root and Indian nettle. These plant tissues contained relatively large amounts of actinidine in addition to two or three other compounds (isodihydronepetalactone, isoiridomyrmecin, iridomyrmecin).

GC-MS analysis was also done for *M. trifoliata* (buckbean) leaves, as well as other silver vine tissues (stem and leaves) and three tinctures that we used in our experiments (**Figure 20**). Only dihydroactinidiolide was detected in the *M. trifoliate* leaves. Of the two cats who responded to *M. trifoliate* it was Cat V who responded longest. She was also the one who responded the most frequent and the longest to dihydroactinidiolide. We observed positive responses from all cats, even several hours after applying 2 sprays of the tinctures. This finding suggested effective extraction of some of the cat-attracting single compounds in the tinctures. Indeed, when the amount of single compounds in the tinctures is expressed per gram dried plant material, these numbers surpass those obtained with the methanol:dichloromethane extraction method. However, we used 5 volumes of ethanol per 1 volume of plant tissue and therefore the quantity of single compounds per mL of tincture was relatively low (**Figure 20**). Since roughly 1/5^th^ of a milliliter of tincture was sprayed on the fabric, this implies that domestic cats are able to detect quantities of just a couple of micrograms.

While the fruit galls of the silver vine plant already contained a large amount of actinidine, we found 3 × more actinidine in the leaves, where it was also the dominant cat-attracting compound. Neonepetalactone and isoneonepetalactone were not detected in the leaves, but the other compounds were present in similar amounts when compared to the fruit galls. Interestingly, the response of cat Z where she would not touch the object was seen only for actinidine and the silver vine leaves. As expected, fewer compounds and lower quantities were detected in silver vine stem. Actinidine was still the most abundant compound, but four other compounds were found as well. A piece of silver vine stem harvested in East Asia (sold by Mew Neko, Austin, TX, USA) that was not used in the bioassays, but was found particularly popular among some of the cats (mostly cats N and Z) was analyzed as well and compared to the locally-grown, younger stem. The popular piece of silver vine stem contained neonepetalactone and more actinidine, but no dihydroactinidiolide was detected in the other stem.

## Discussion

In this study, we aimed to gain a better understanding of the “catnip response” in domestic cats by analyzing their behavior to different cat-attracting plants and chemically synthesized volatiles found in these plants. We observed differences between cats in their behavior to these plants and compounds that raise interesting questions about the way these compounds are perceived by cats and the underlying mechanism of olfactory sensation. We will address the most pertinent questions in more detail below.

For this study, cats were exposed to cat-attracting plants and their volatile, active, single compounds for nearly 1,000 hours over a period of more than 2 years. To the best of our knowledge, this is the longest exposure of cats to catnip and catnip-like material ever documented. Following the final olfactory bioassay for this study in late 2020, *L. tatarica* wood and dried *A. polygama* leaves were made available to the cats in a nearly continuous manner. Since authors SB and EMB have been living together with the study cats before, during, and also after the study, we were able to closely monitor any potential negative health effects as the result of exposure to these olfactory stimuli. No adverse health effects, either physically or mentally, were observed, up to the publication date. These results support the current believe that these plants and their active compounds are safe (in the amounts that were available to them in this study) and offer an excellent source of environmental enrichment.

Unfortunately, catnip is sometimes referred to as “kitty crack”, the euphoric, blissful response to the cat- attracting plants considered a “high” or cats “tripping”. This negative association can prevent cat guardians from offering olfactory enrichment. However, the plants that produce THC, the active compound in marijuana or weed (cannabis plant), cocaine or crack (coca plant), or heroin (opium poppy) are not related to any of the cat-attracting plants. Cannabis, the coca plant, and opium poppy are all species in families (Cannabaceae, Erythroxylaceae and Papaveraceae, respectively) that do not include the cat-attracting plant species (families: Euphorbiaceae, Actinidiaceae, Caprifoliaceae and Lamiaceae). Furthermore, the structures of THC, LSD, cocaine and heroin are much more complex than the cat- attracting iridoids and about twice the molecular weight (**Supplementary Figure 13A**). Two major differences between the above-described psychoactive drugs and the cat-attracting compounds are their route of entry and subsequent receptor binding. None of the psychoactive drugs are volatiles. The drugs need to end up in the blood (e.g., intravenous injection, orally, nasal tissue (snorting ≠ smelling), smoking) and subsequently bind receptors in the brain for them to be active; they do not elicit a response after smelling them. Nepetalactone only has an effect when the volatile is bound to the olfactory receptors after the cat inhaled air through the nose; it has no effect when absorbed into the blood after oral administration (Waller, Price and Mitchell 1969). Another difference between psychoactive drugs and cat- attractants is the duration of the response. While the “catnip response” lasts seconds to several minutes at most and can be easily interrupted, the effects of administering cocaine, heroin, LSD or smoking cannabis last for hours and cannot be stopped. Authors SB and EMB, who live with the cats, did not observe any withdrawal, abnormal behavior, or changes in behavior of any of the participating cats after the cat-attractants had been taken away from the cats. In the contrary, we believe we observed more positive interactions between cats in the testing area when the cat-attracting plants or single compounds were present, although this was not measured.

Much about the “catnip response” still seems riddled in mystery. We are clueless as to what the reason for, or biological function of, the response is and why it is only seen in felines. It has been hypothesized that a cat rubbing plant material with insect-deterrent compounds could reduce the number of mosquito bites (Uenoyama et al. 2021) and thereby prevent mortality due to mosquito-borne diseases. However, such diseases are uncommon in felines. Moreover, given the large range of mosquitoes in terms of both geographical spread and species that can serve as a host for blood meals, it would be likely that similar behavior would have evolved in other species. It is known that dihydroactinidiolide is present in secretions of the supracaudal and tail glands of the red fox, as well as in their urine. Our bioassays have demonstrated that domestic cats respond to dihydroactinidiolide. These observations justify revisiting the hypothesis that the “catnip response” is elicited by extreme quantities of compounds similar in structure to semiochemicals that serve in the communication between individuals from the same species (pheromones) (Todd 1963, Albone 1975).

Another big unknown is which olfactory receptors are bound by the volatile cat-attracting molecules. Our study shows that some cats respond more strongly to actinidine than to lactones, while others do not respond to actinidine at all. This suggests that cats may have genetic differences in the receptor(s) that detect(s) these various compounds. Since the number of odorants far exceeds the number of olfactory receptors, it is believed that a single receptor can bind different odorants, but also that the same odorant can bind to different receptors, albeit probably with different affinities. This combinatorial olfactory receptor code and our poor understanding of structure-odor relationships make it difficult to speculate about the molecular mechanism that is involved in the “catnip response” and what might explain the difference in response between cats to the lactones and actinidine.

The most obvious difference between actinidine and all other cat-attracting compounds is that actinidine contains a pyridine ring instead of a lactone, while still retaining the cyclopentane ring. These different features of actinidine may allow for binding to different receptors that are only expressed in some cats, or to mutated versions of the same receptors that bind the type I and II lactones.

We found that the “catnip response” could not be elicited by catnip in cats habituated to silver vine or Tatarian honeysuckle. Since silver vine and honeysuckle both lacked *cis*-*trans*-nepetalactone, *trans*-*cis*- nepetalactone and dihydroactinidiolide, it may be possible that the other type I and II lactones in silver vine and Tatarian honeysuckle bind the same receptor(s) as nepetalactone and dihydroactinidiolide.

It has recently been shown that cats also respond to nepetalactol (Uenoyama et al. 2021), which is a reduced form of nepetalactone and similar in structure, but lacks the carbonyl group of the lactone (**Supplementary Figure 13B**). This suggests the carbonyl functional group may not be required for the lactones to engage with the receptor(s). Keesey and colleagues recently showed that *Actinidia arguta* leaves contain moderate amounts of nepetalactol, but only trace amounts of iridomyrmecin and actinidine, and no nepetalactone (Keesey et al. 2019). However, the *A. arguta* leaves that we tested previously did not elicit the “catnip response” in any of 8 domestic cats (Bol et al. 2017).

Nelson and Wolinsky reported that cats responded positively to both iridomyrmecin and isoiridomyrmecin (Nelson 1968), which is in agreement with the results from Uenoyama *et al*. and our findings. They also tested *cis*-*cis*-iridolactone (molecules XXV and XXVI in reference (Wolinsky et al. 1965)), epimers of iridomyrmecin and isoiridomyrmecin that have not been tested by others (**Supplementary Figure 13B**). Surprisingly, none of the cats responded positively to the *cis*-*cis*-iridolactones, despite their structural similarity to iridomyrmecin and isoiridomyrmecin (Nelson 1968). The only difference between these compounds is the methyl group on the cyclopentane ring, which is inverted on the *cis*-*cis* variants. This suggests that (the orientation of) this methyl group, along with the cyclopentane ring, might play an important role in binding to the receptors. While we did not see statistically significant differences in duration of the “catnip response” between compounds, the response to both neonepetalactone and isoneonepetalactone seemed lower than all other type I and II lactones. While the methyl group on the 5- membered ring is not inverted on neonepetalactone and isoneonepetalactone as it is on the presumably inactive *cis*-cis-iridomyrmecin and *cis*-*cis*-isoiridomyrmecin, it is planar to the cyclopentene ring. This planar orientation of the methyl group is different from all the other active type I and II lactones (**1** – **4**, **7** and **8**, **Figure 12**) and may possibly account for the seemingly reduced response duration of the cats to neonepetalactone and isoneonepetalactone.

Nelson and Wolinsky also found that cats responded positively to another compound that has not been tested by others: the bridged bicyclic matatabiether (**Supplementary Figure 13B**). Interpretation of the results from the work done by Nelson and Wolinsky is challenging however, because the experimental methods of the bioassays were not described in detail. Furthermore, no clear definition was given when a response was considered positive. Sniffing, licking or biting in the absence of head rubbing may have been considered a positive response (Katahira and Iwai 1975, Sakan et al. 1960).

We observed differences in attractiveness of plant materials to cats that may be explained by their chemical composition. The popularity of silver vine fruit galls and leaves may be explained by the presence of some compounds that were not detected in any of the other plants, the large number of cat- attracting compounds present, or the large quantity of actinidine. However, the combined results from the bioassays and GC-MS analysis also suggest there may be other, unidentified, cat-attracting compounds present in the cat-attracting plants, especially Tatarian honeysuckle. While no cat-attracting compounds were detected in any of the fragrances tested, the “catnip response” was seen twice after fabric with Drakkar Noir was made available to cat A. While cats O and V did not respond to actinidine, yet both cats responded to the Tatarian honeysuckle tincture, in which we only detected actinidine. We observed dishabituation of cat O to Tatarian honeysuckle, both after habituation to silver vine fruit galls (twice) and catnip leaves had occurred, while we were not able to detect any compounds in the honeysuckle that were not present or present at lower levels in the silver vine or catnip samples. The chemical composition of valerian root was similar to that of Tatarian honeysuckle and Indian nettle root. However, cat O responded significantly longer to Tatarian honeysuckle than to valerian root. Furthermore, cat Z did respond to Tatarian honeysuckle and Indian nettle root, but never to valerian root. Possibly, some plant samples contained odorants that had a repelling effect on some cats (e.g., valerian root). Indeed, the valerian root tincture did not contain any known cat-attracting compounds that were not also present at comparable levels in the dried valerian root, yet cat Z responded to the tincture and not to the dried plant material. Previous quantitation where compounds were extracted from plant tissues for not 2, but 7 days in dichloromethane yielded a similar pattern: none of the quantitated compounds was found to be only present or at higher levels in Tatarian honeysuckle compared to catnip and silver vine (Bol et al. 2017).

Other cat-attracting single compounds may have been extracted, but not identified. Cats detect the volatile compounds emitted by the plant tissues in the air. For our chemical analysis we chose to extract the compounds from the plant tissues using solvents, to enable accurate identification and quantification, as was done in our previous work (Bol et al. 2017). There may be some discrepancy between naturally emitted volatiles from the plant materials and those that can be extracted with solvents. Headspace (airspace) analysis may provide results that better represent what cats detect than the solvent extraction methods, but most headspace analysis methods (SPME or purge and trap) cannot be easily quantified. Our quantification data however is a useful guide to the presence and relevant amounts of cat-attracting compounds present within each plant material tested in this study. A useful outcome of this study is that we were able to quantify a large group of cat-attracting compounds from the same plant materials that were exposed to the cats for behavioral analyses, enabling us to link some of the chemistry with cat behavior.

Our interest in cat-attracting plants ((Bol et al. 2017) and this work) originates from observations that cats A, N, O, V and Z did not respond to catnip prior to and during the tests done in 2016 (Bol et al. 2017), despite being exposed to the plant material longer and more frequently than any of the other cats in the study. Therefore, it was surprising that the same cats, especially all five, responded to catnip during the tests done for this work. In early 2016, cats A and Z were 9 and 5 years, respectively, and the three littermates were 16 months. It is often claimed that cats younger than 3 – 6 months do not respond to catnip. However, it might be possible that there is not a tight age cutoff, but that the “catnip response” is something that can or perhaps even needs to be acquired over time. Using the dataset from our 2017 publication, we found that of young adult cats (6 – 18 months) almost 50% fewer responded to catnip than the older cats (**Supplementary Figure 14**). In contrast, the high percentage of cats responding to silver vine was equal among all 4 different age groups. Interestingly, the 5 cats in the study only started to respond to catnip months after they responded to silver vine. Long-term non-responders suddenly responding to catnip have been described in literature before, but for these cases it was suspected to be due to fluctuations in hormones of intact female cats (Todd 1963).

Our observations were done in a small group of cats. The advantage of this approach was that it allowed us to test relatively large numbers of cat-attracting plants and single compounds on the same group of domestic cats in a completely stress-free environment. The small, homogenous study population did not prevent us from obtaining answers to the main research questions. With this small group of cats, we were able to demonstrate that the individual cat’s response to other cat-attracting plants is similar to catnip, and that there is substantial variation between cats in the behavior during the “catnip response”. Furthermore, we learned that cats respond to a large number of type I and II lactones, but also to dihydroactinidiolide and actinidine. Based on results from this study and research recently done by Reiko Uenoyama and colleagues we now know that only 10 – 20% of domestic cats respond to actinidine. We were also able to demonstrate that the behavior to actinidine was different from the lactones. However, it is possible that we missed other differences that will only become apparent when larger numbers of domestic cats and a more heterogeneous population (e.g., different breeds) are tested. Another limitation of this study is that the position of the camera, both the angle and distance, made it sometimes difficult to observe certain activity in the testing area. For this reason, we were not able to study position of the ears and whiskers, pupil size and (excessive) salivation (Bol et al. 2017). Furthermore, the fixed camera position sometimes complicated discrimination between behaviors, e.g., licking or head rubbing. While it also would have prevented us from studying any delayed behavior in response to the olfactory stimuli that would have occurred off-camera, nothing indicated that such behavior did occur.

Studying more cats, plants and single compounds will undoubtedly reveal additional cat-attracting plants and single compounds that can elicit the “catnip response”. Some of these molecules may affect even fewer cats than actinidine does. Newly identified cat-attracting volatiles do not necessarily need to come from plants. Both plagiolactone and gastrolactone (**Supplementary Figure 13B**) are similar in structure to the cat-attracting type I and II lactones, and are produced by different species of leaf beetles (Meinwald et al. 1977, Blum et al. 1978). We are not aware of any publication reporting the detection either plagiolactone or gastrolactone in plant tissue. It remains to be determined whether these two compounds can elicit the “catnip response”. Furthermore, it may be possible to synthesize many novel cat-attracting compounds that do not occur anywhere in nature.

In conclusion, we have performed a comprehensive study of the “catnip response” of domestic cats to five plants, 10 single compounds, and several other samples. We observed that while responses between cats were highly variable, the behavior of individual domestic cats to diverse cat-attracting plants as well as all lactones was quite similar. Interestingly, the response to actinidine was most divergent, with several non-responders and a small percentage of cats who preferred actinidine over all other compounds.

Collectively, these results have increased our understanding of the “catnip response” in terms of both behavior and the chemical compounds that elicit it. It has also revealed potential differences in the perception of compounds between cats that warrant further investigation into the underlying genetics of cat odorant perception and the mechanism(s) of action of these compounds.

## Acknowledgements

We are grateful to Yolanda Villarreal Acosta (The University of Texas at San Antonio) for guidance on establishing an institutional animal care and use committee. We gratefully acknowledge Naoki Yamanaka (University of California, Riverside) for translating scientific literature written in Japanese, and Zita Peterlin for editing the manuscript. We thank Dave Algar (Government of Western Australia, Department of Biodiversity, Conservation and Attractions) for providing lyophilized *Acalypha indica* roots. We also thank Frances Mae Tenorio and Maria Brink for help with the analysis of the cat behavior.

## Author contributions

**SB** conceived, designed, and coordinated the study, performed experiments, collected and analyzed data in GraphPad Prism and BORIS, and wrote the manuscript. **AS** synthesized the single compounds, extracted and performed chemical analysis of the samples, created Figure 12, Supplementary Figure 13, and edited the manuscript. **EMB** assisted with experiments, created Supplementary Figures 3 and 7, and edited the manuscript. **GRF** extracted and performed chemical analysis of the samples, analyzed the data, and edited the manuscript.

---

**Supplementary Figure 1.**
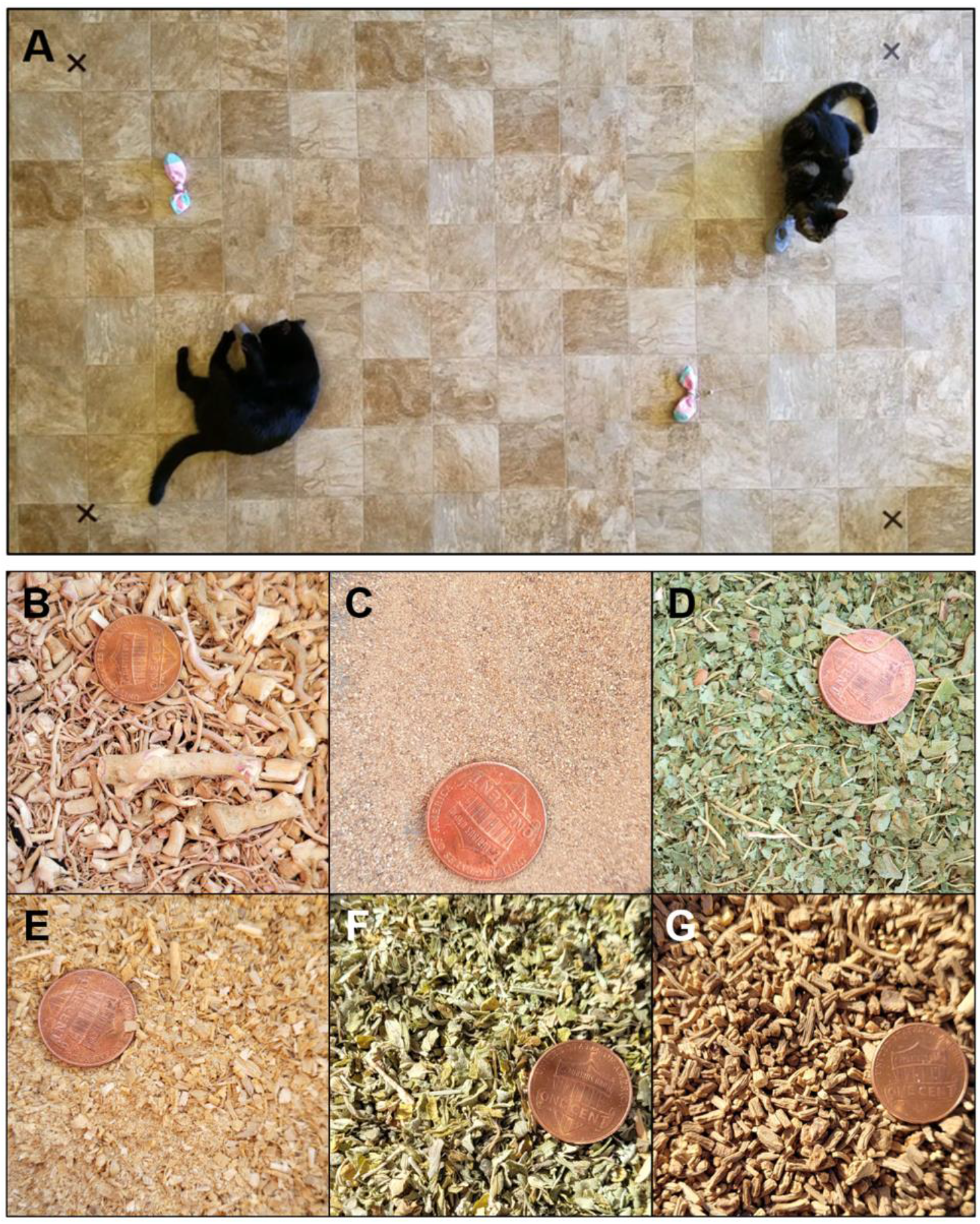
Top (**A**): The testing area with 4 mounted socks. The black x’s served to assure the relevant area of the testing area was being captured by the camera. Bottom: Close-up photographs of the plant materials used in the study: (**B**) lyophilized and cut *Acalypha indica* or Indian nettle root, (**C**) dried, powdered *Actinidia polygama* or silver vine fruit gall, (**D**) dried and cut Texas-grown *A. polygama* leaves, (**E**) *Lonicera tatarica* or Tatarian honeysuckle sawdust, (**F**) dried and cut *Nepeta cataria* or catnip leaves, and (**G**) dried and cut *Valeriana officinalis* or valerian root. A United States penny (19 mm diameter) is used as a size reference.

**Supplementary Figure 2.**
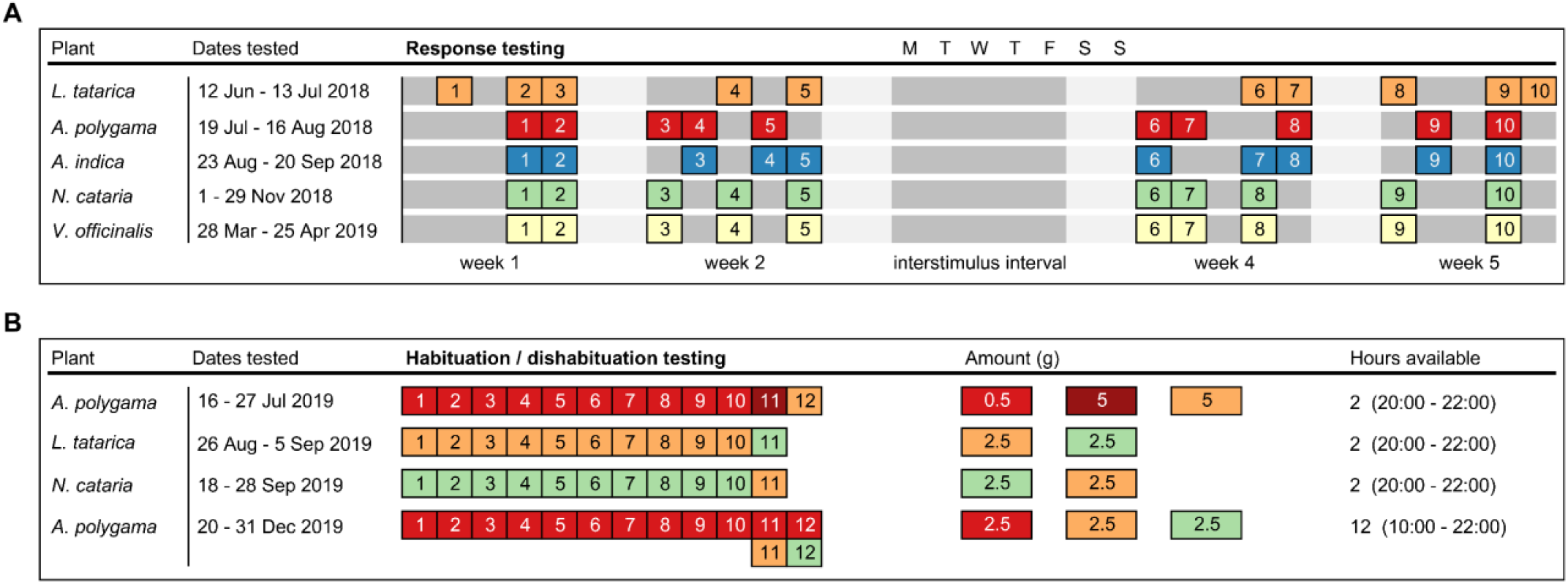
Timeline for testing cat-attracting plants. (**A**) Each cat-attracting plant was tested on 10 different days (no. 1 – 10), 10 hours per day, to learn more about response duration, response frequency, and behavior during the responses. The tests were done in two periods of two weeks, separated by an interstimulus interval of at least 9 days. Testing was done 2 – 3 days per week. There was always at least one week between testing the different plants (see “Dates tested”). (**B**) Three different cat-attracting plants were offered for 10 consecutive days (2 or 12 hours per day) to test for habituation. After 10 days the cats were offered a different plant to test dishabituation. Plant materials are color-coded according to the color scheme used in (**A**).

**Supplementary Figure 3.**
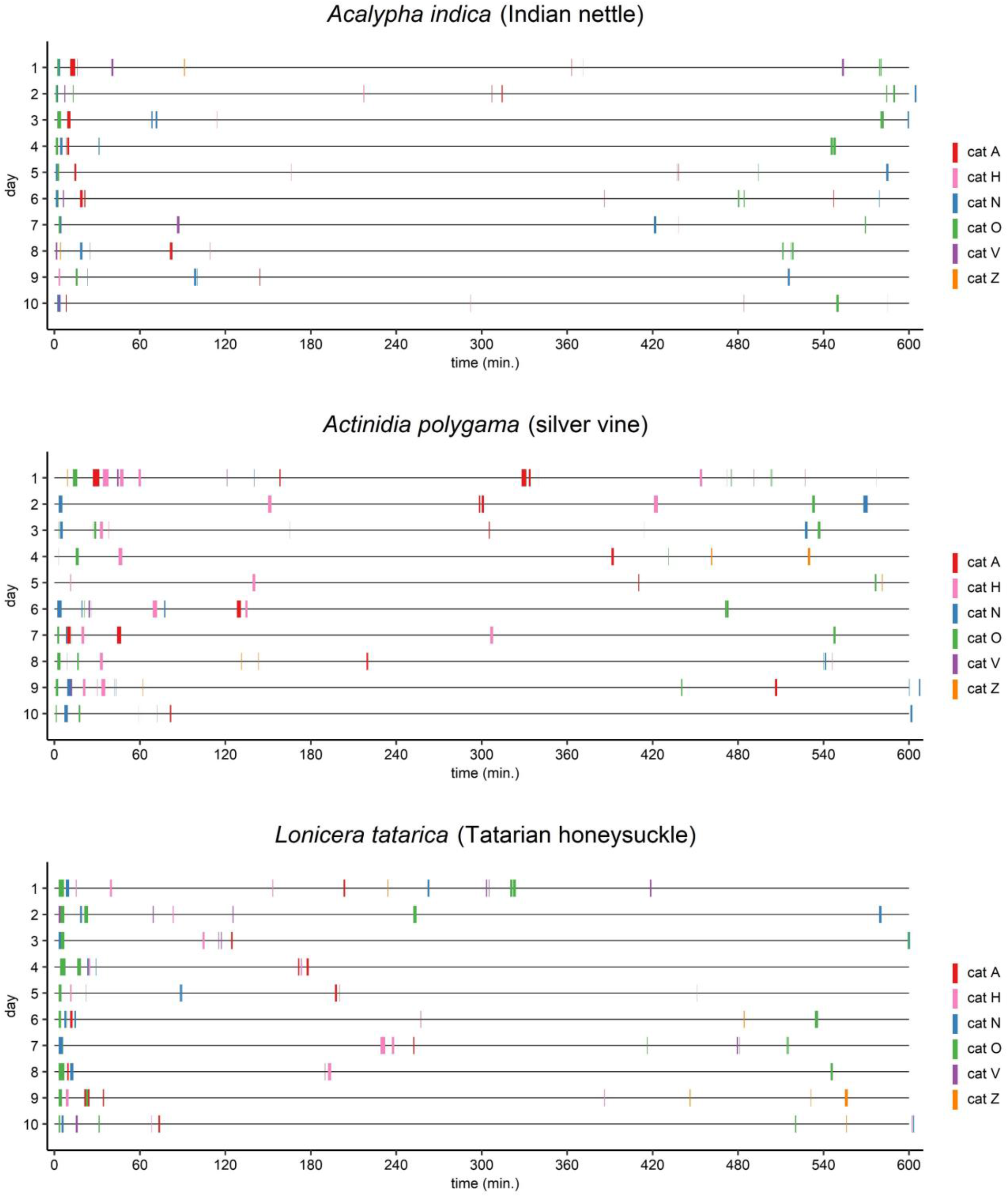
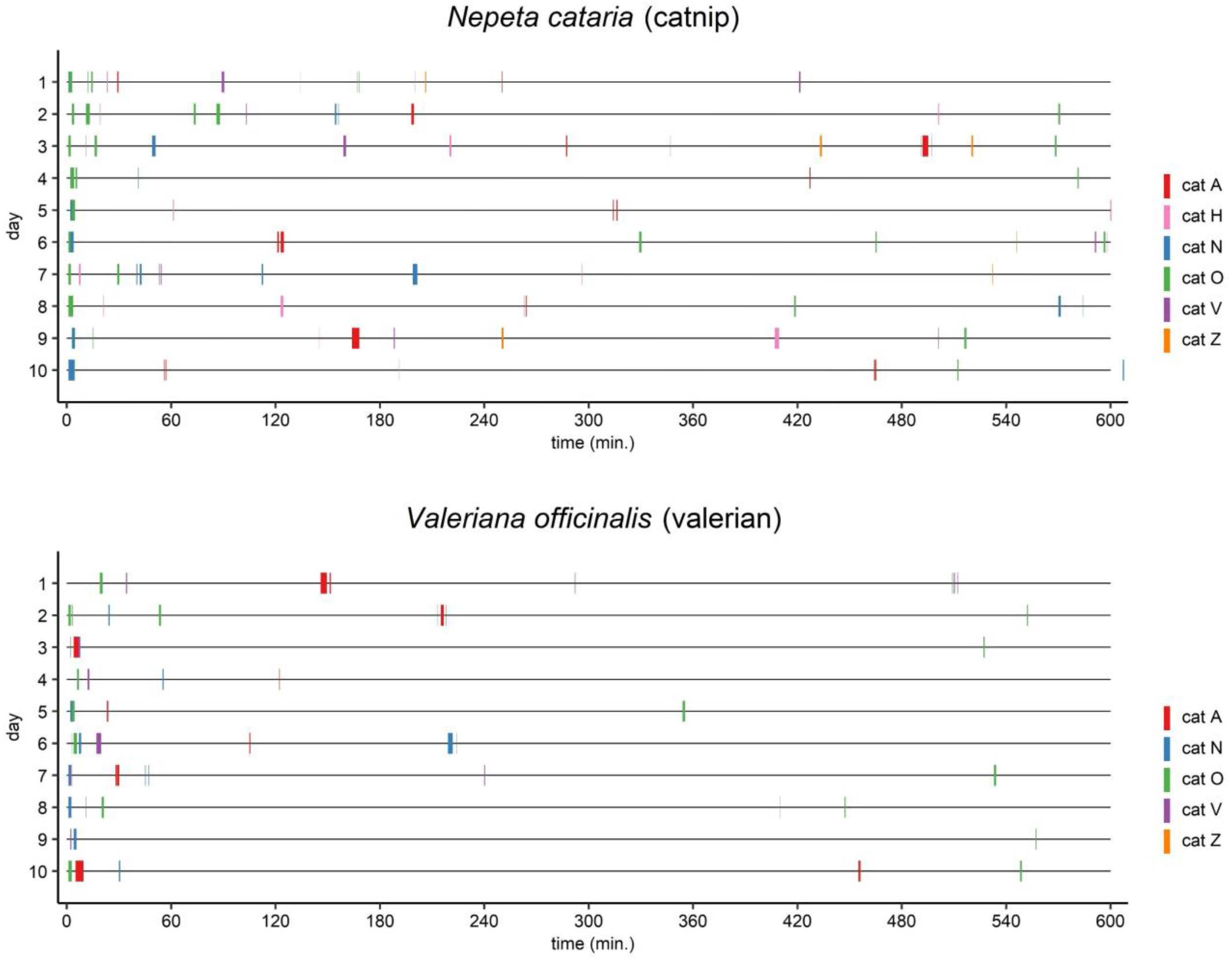
Graphical overview showing time of day of the responses, response frequency and response duration for the 6 domestic cats to *A. indica*, *A. polygama*, *L. tatarica*, *N. cataria*, and *V. officinalis*. Each plant was available on 10 days for 10 hours (600 minutes), between 9:30 and 19:30. Responses that lasted only a couple of seconds sometimes do not show.

**Supplementary Figure 4.**
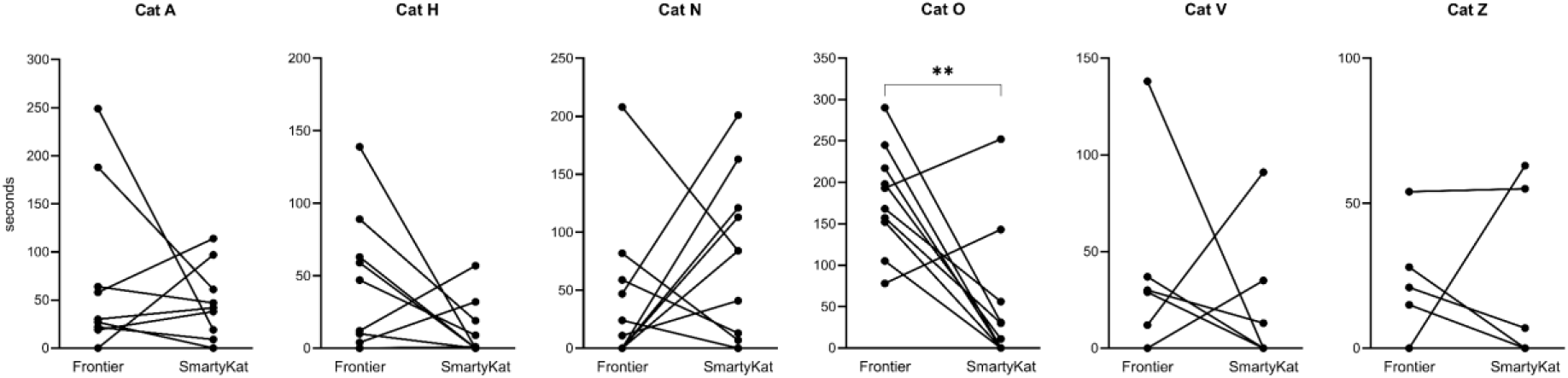
Response duration of 6 domestic cats to two different sources of *N. cataria*. Each dot represents the total response duration on one of the 10 testing days. The median response duration of cat O to catnip from Frontier was significantly longer than the median response duration to catnip from SmartyKat (P = 0.0098, Wilcoxon matched-pairs signed rank test). ** P < 0.01

**Supplementary Figure 5.**
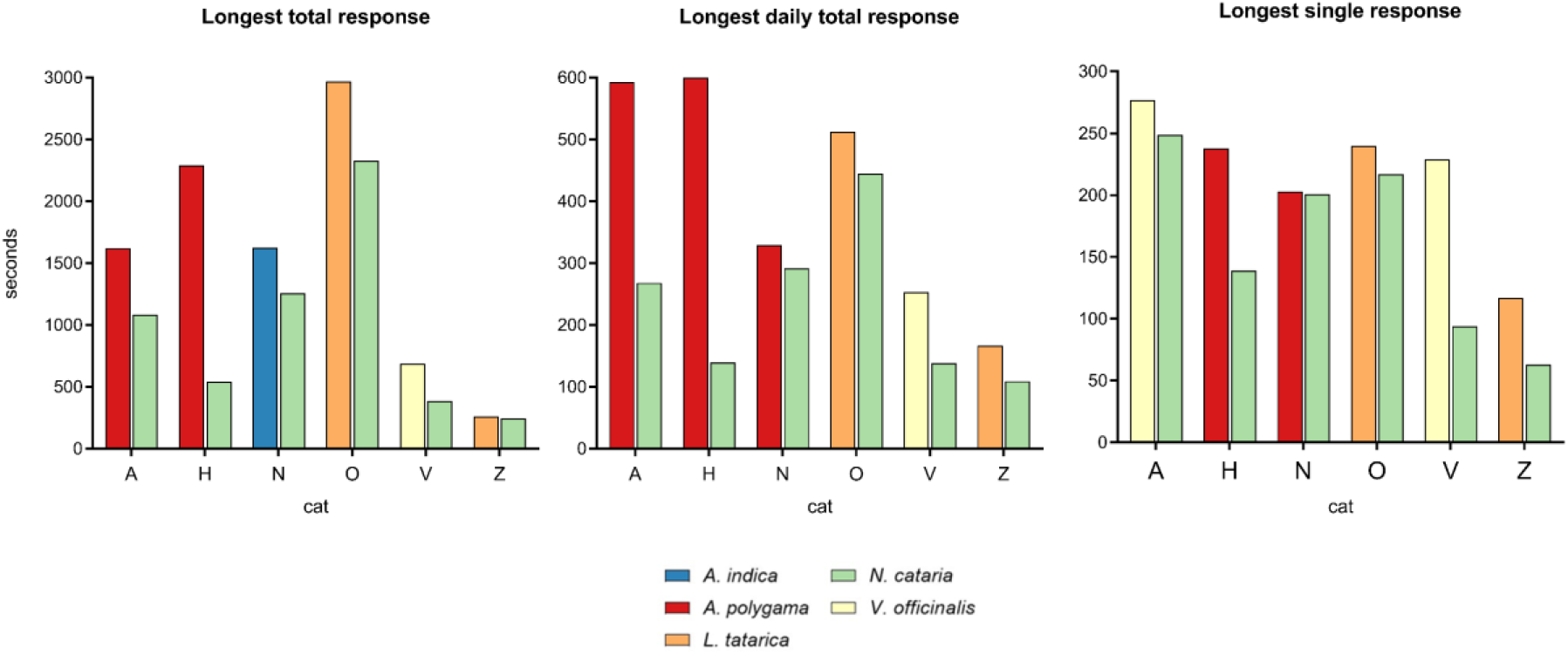
The longest total response (100 hours availability), the longest daily response (10 hours availability), and the longest single response for each cat, compared to *N. cataria* (catnip).

**Supplementary Figure 6.**
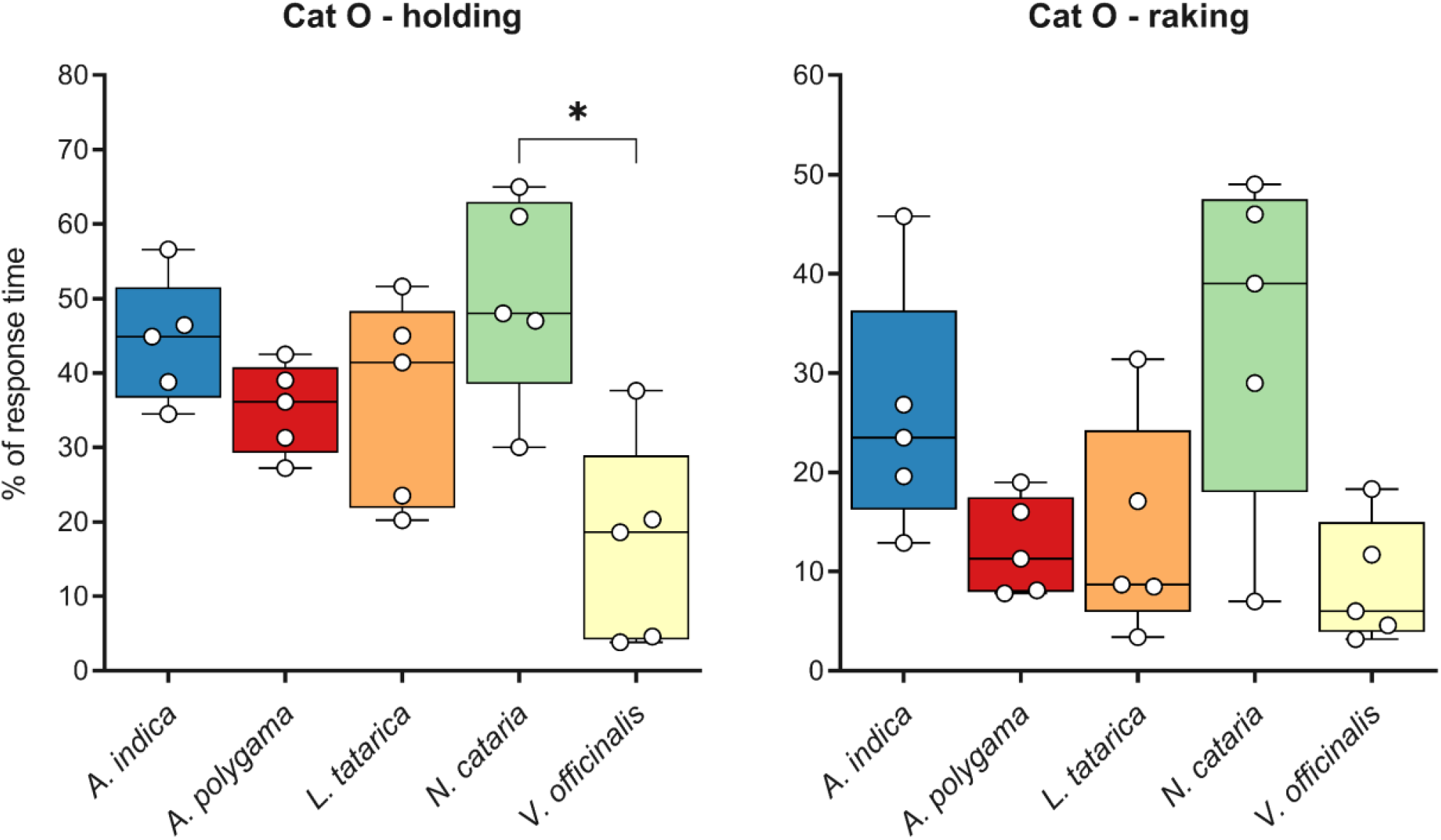
Box and whisker plots showing the time spent holding and raking by cat O in response to 5 cat-attracting plant species. Data from 5 responses nearest to 60 seconds are shown for each plant. Time is expressed as the percentage of the total response duration. This is a different representation of the data shown in Figure 7C. The differences between the plants for both holding and raking were statistically significant (Kruskal-Wallis, P < 0.05). The P value shown is from Dunn’s post-hoc test. * P < 0.05

**Supplementary Figure 7.**
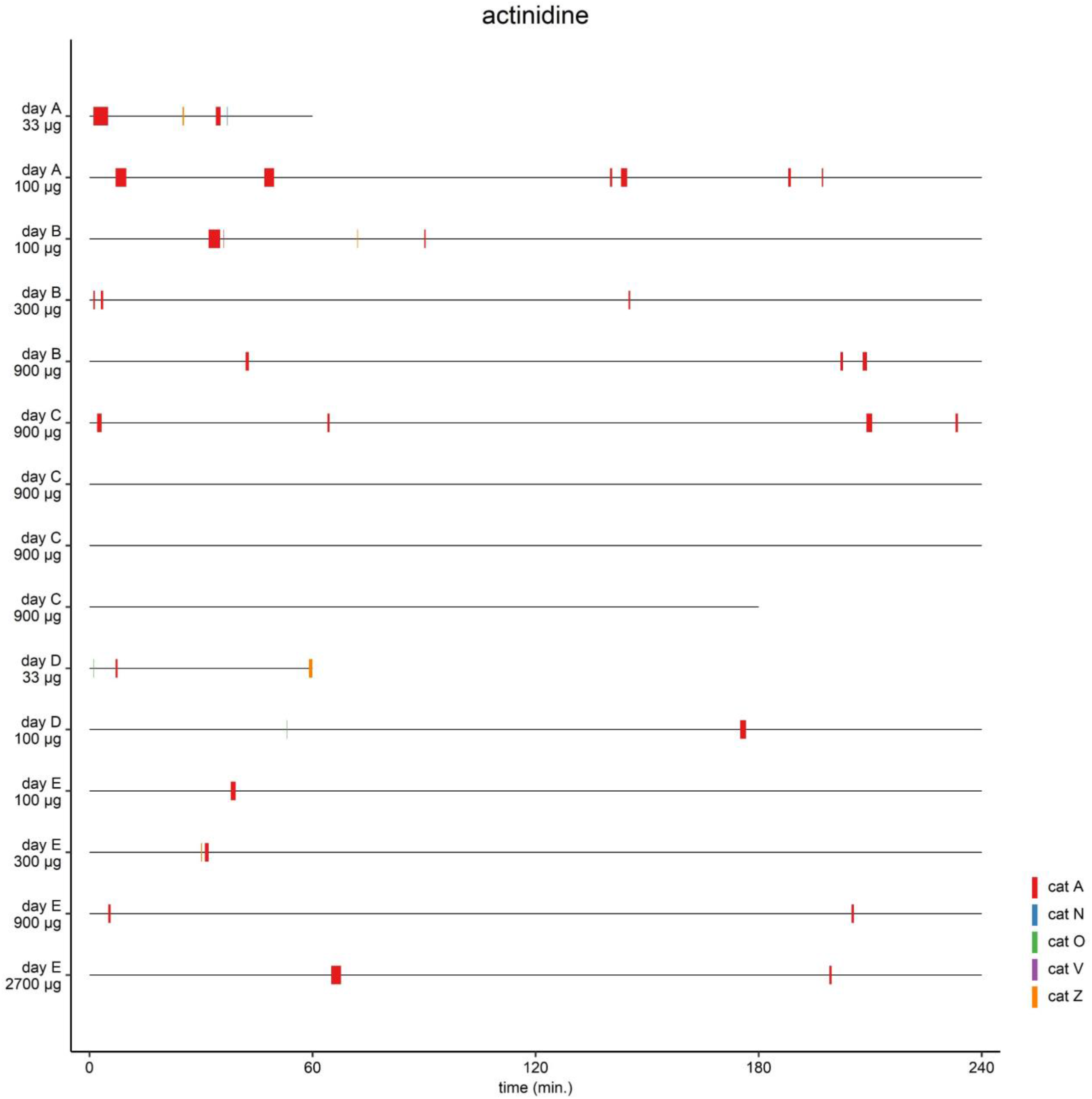
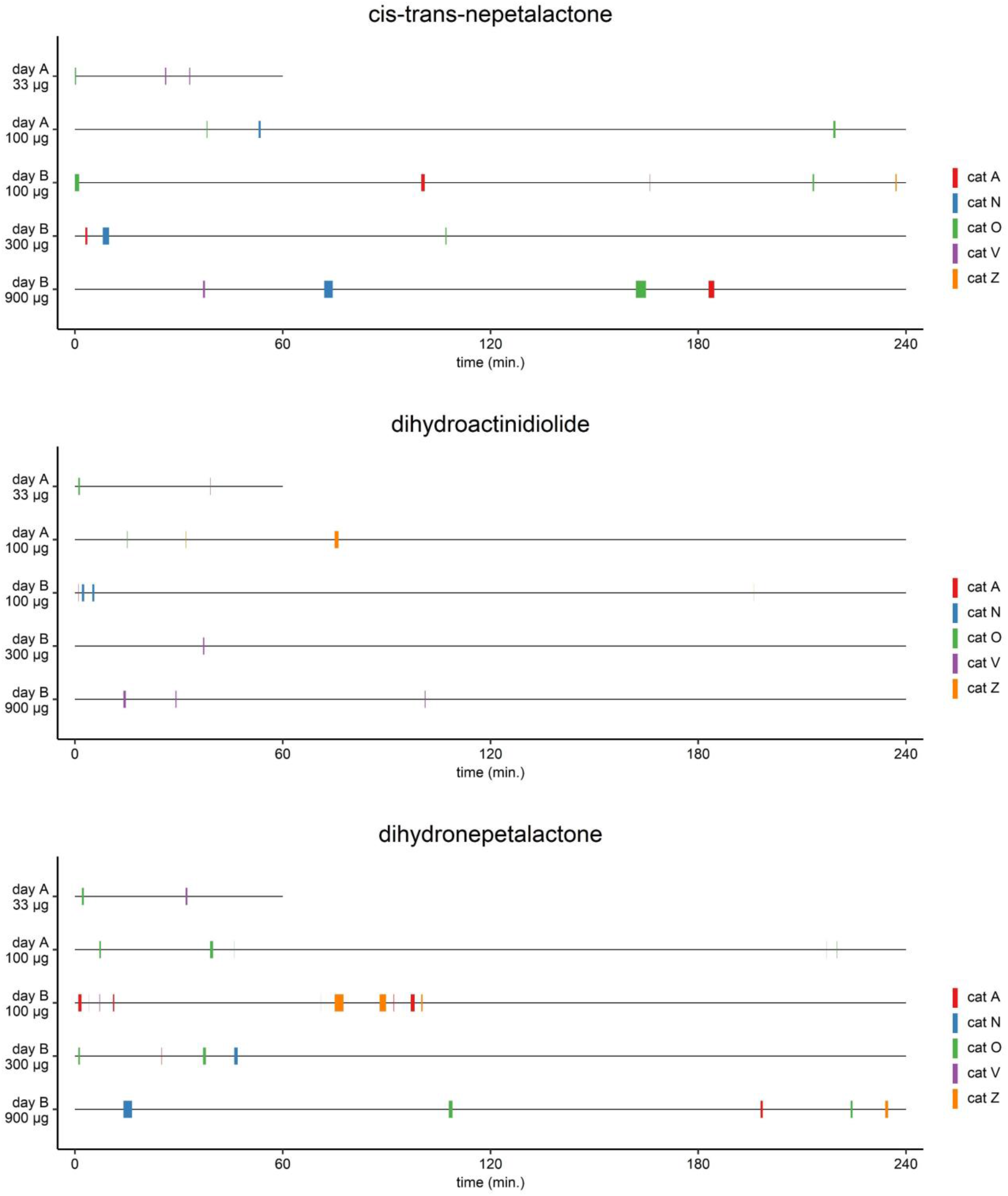
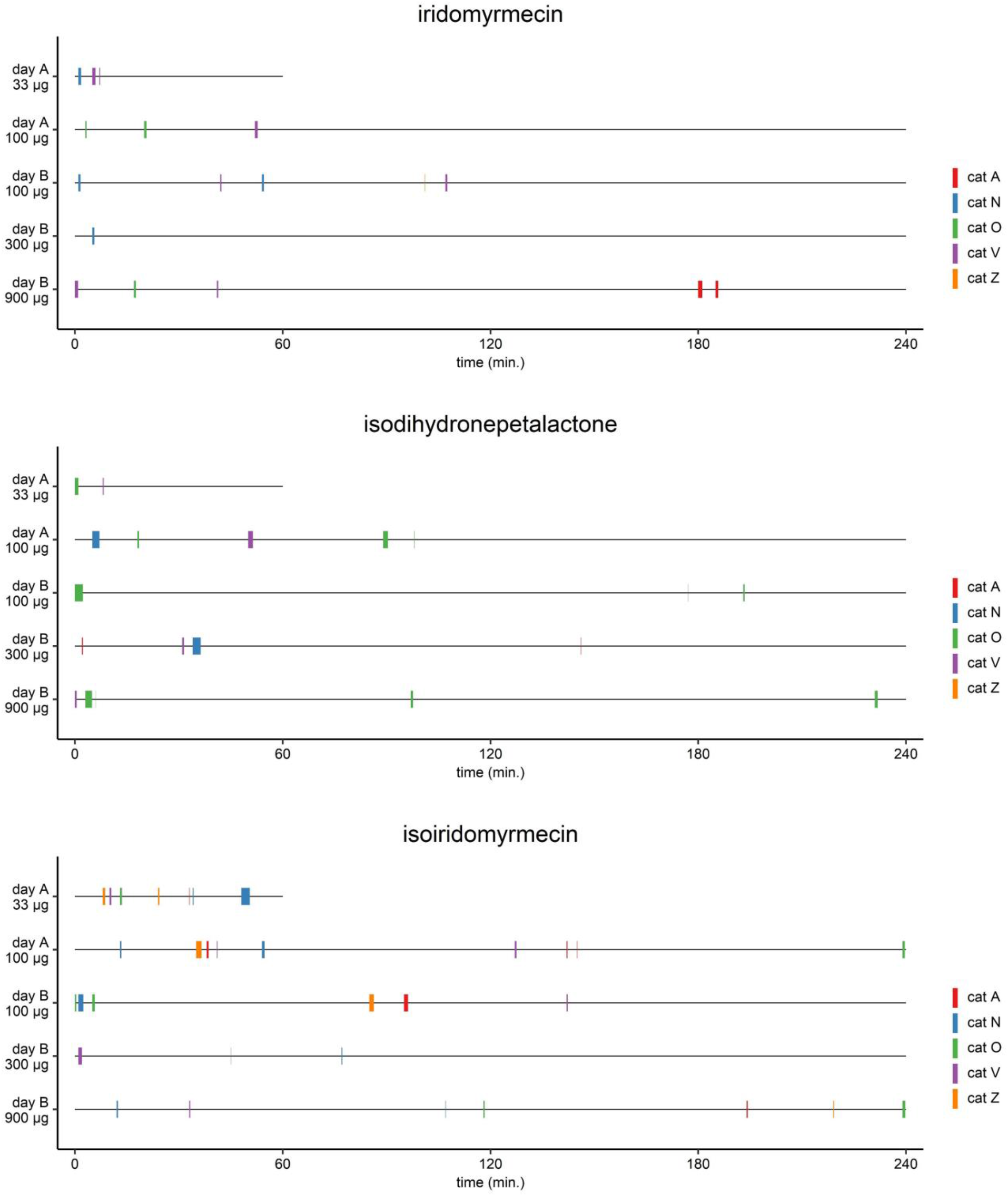
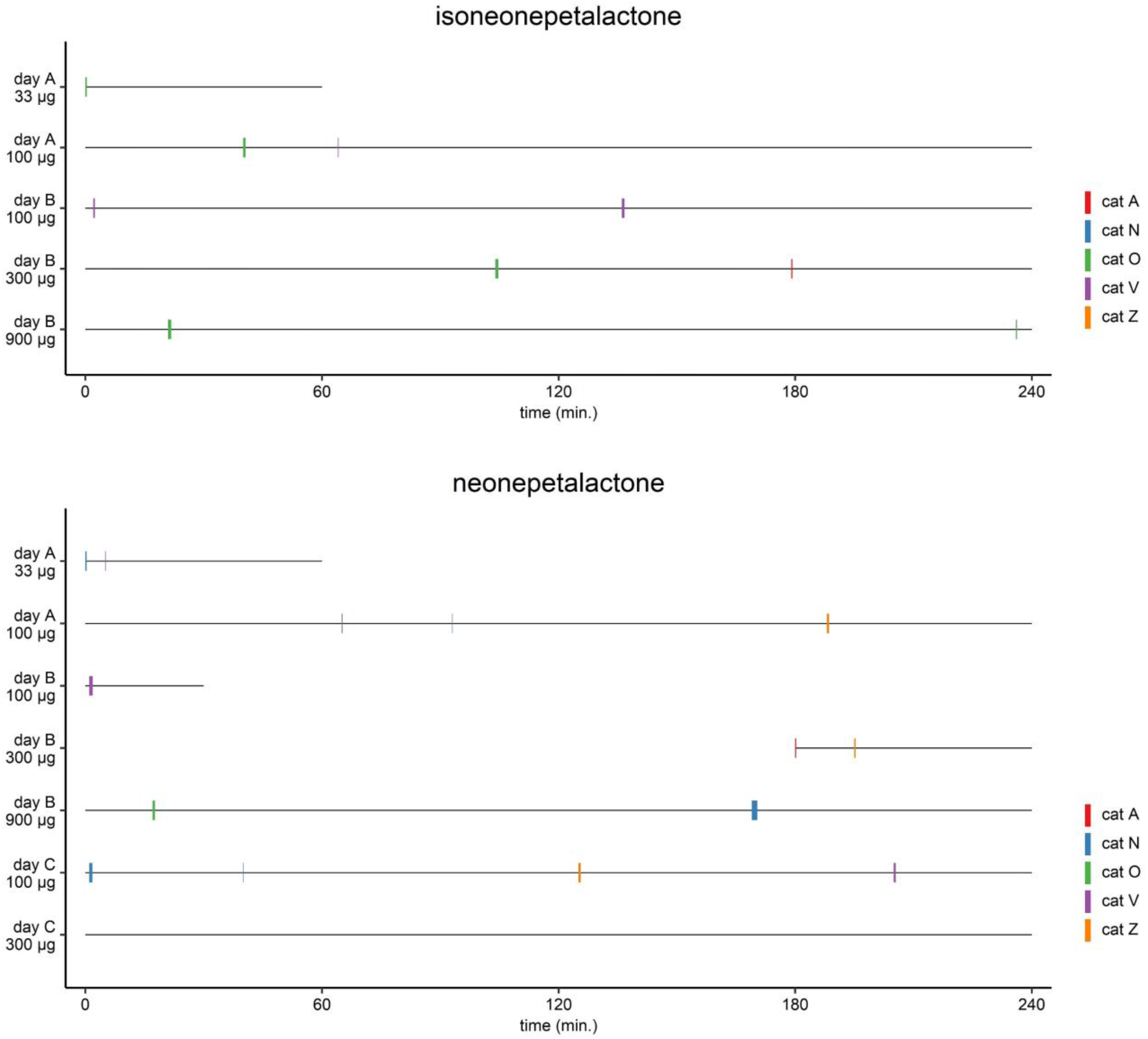
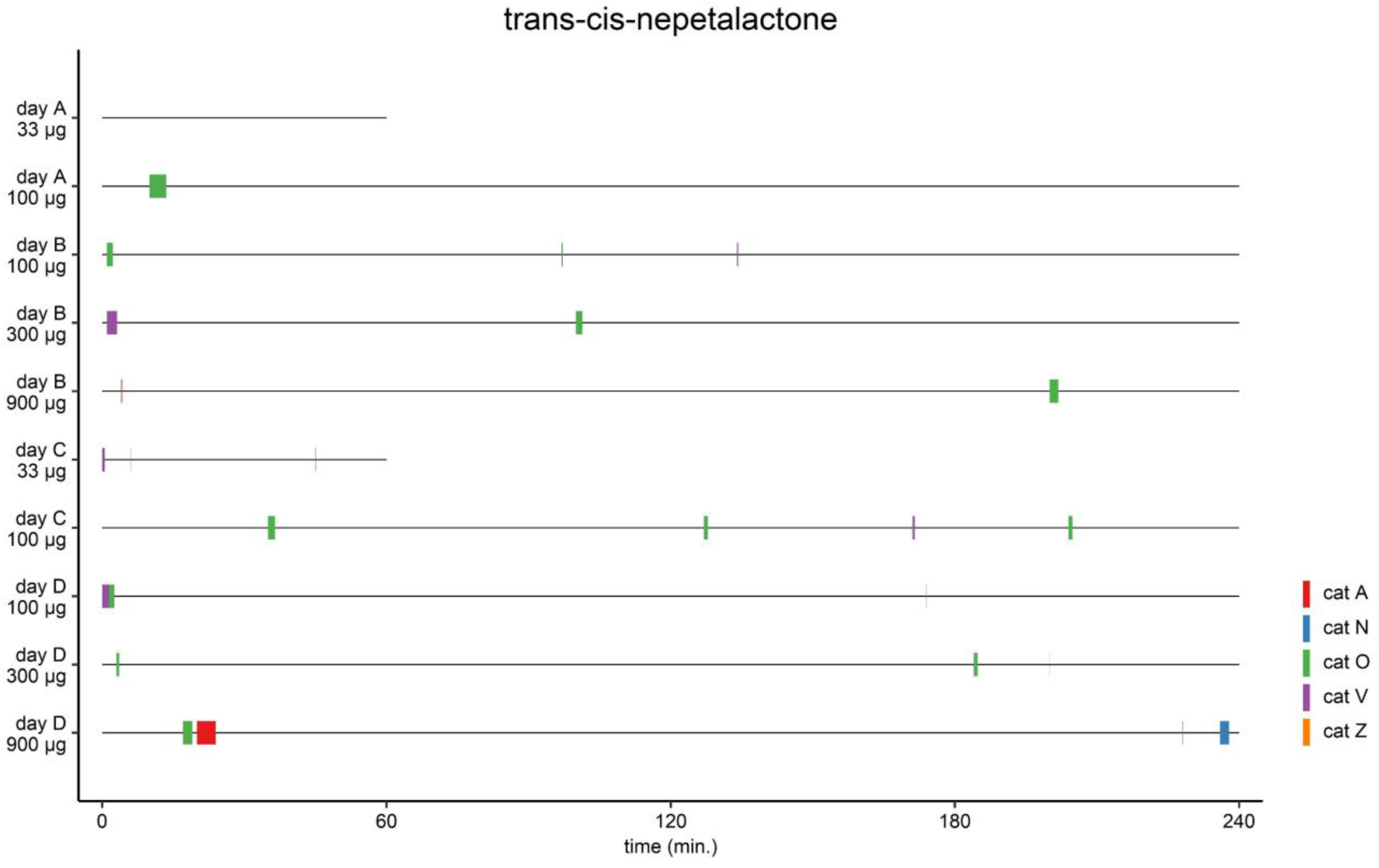
Graphical overview showing time of day of the responses, response frequency and response duration for the 5 domestic cats to the single compounds **1** – **10** (**Table 3**). Each compound was available on at least 2 days for a total of 17 hours. On the first day (day A), 33 and 100 µg were tested for 1 and 4 hours, respectively, typically in the afternoon. On the second day (day B), 100, 300 and 900 µg were tested, each for 4 hours, starting at 7:30, ending at 19:30. Because of the absence of response by some cats (**2** and **9**), technical problems (**5**), or testing a higher amount (2700 µg, **9**) some compounds were tested on additional days or for an extended period of time. Responses of only a couple of seconds sometimes do not show in the figure.

**Supplementary Figure 8.**
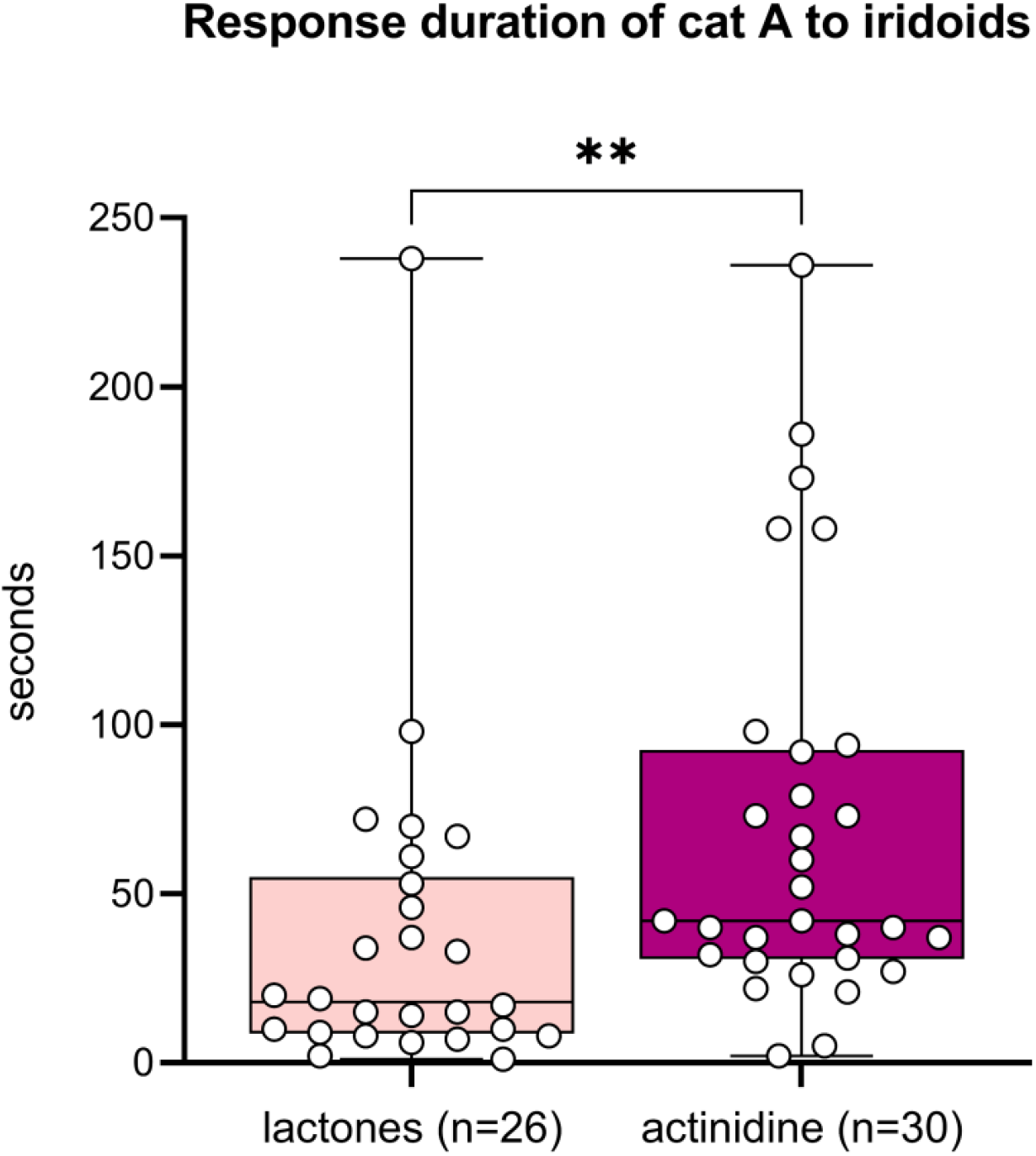
Duration of individual responses of cat A to the lactones and actinidine. Each individual response is shown as a dot. The difference in response duration between the lactones and actinidine is statistically significant (Mann-Whitney test). ** P < 0.01

**Supplementary Figure 9.**
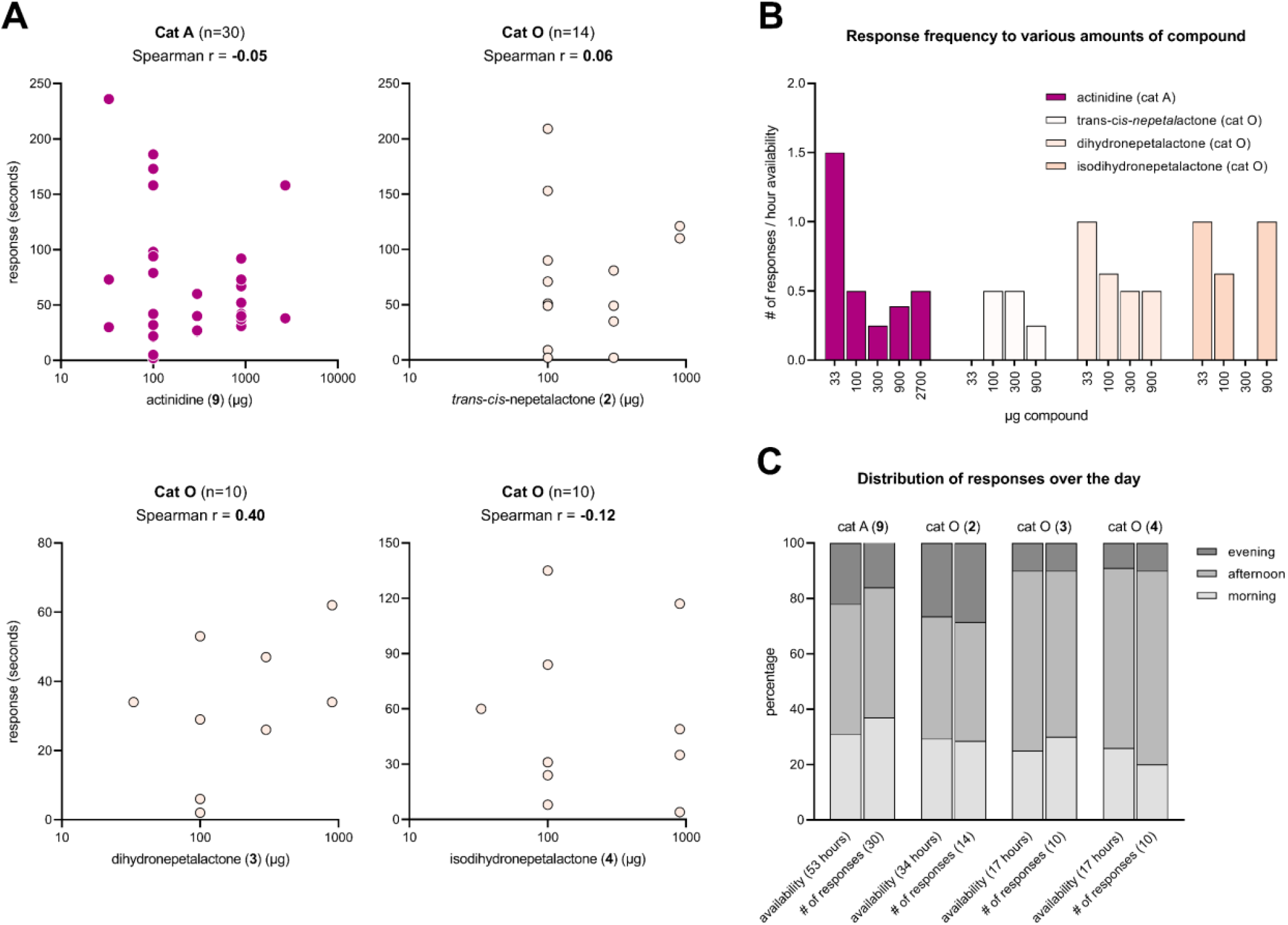
Data from compounds for which at least 10 responses from an individual cat were observed were used to study correlation between response duration/frequency and the amount of compound used, as well between response frequency and time of the day. (**A**) Response time to single compounds plotted against the various quantities of the compounds used in the tests: 33, 100, 300, 900 and 2700 µg. The quantities are shown on a log10 scale. (**B**) Response frequency per hour availability shown for the different quantities of single compounds tested. (**C**) Distribution of responses over the day (morning, afternoon and evening) compared to the distribution the olfactory stimuli were available to the cat (morning, afternoon and evening). Both the number of responses and the time each compound was available to the cat are expressed as a percentage. The total number of responses and hours availability are shown between parentheses. Bold numbers refer to the compound (**Table 3**). The Fisher exact test was used to test for differences in distribution (all P values > 0.05).

**Supplementary Figure 10.**
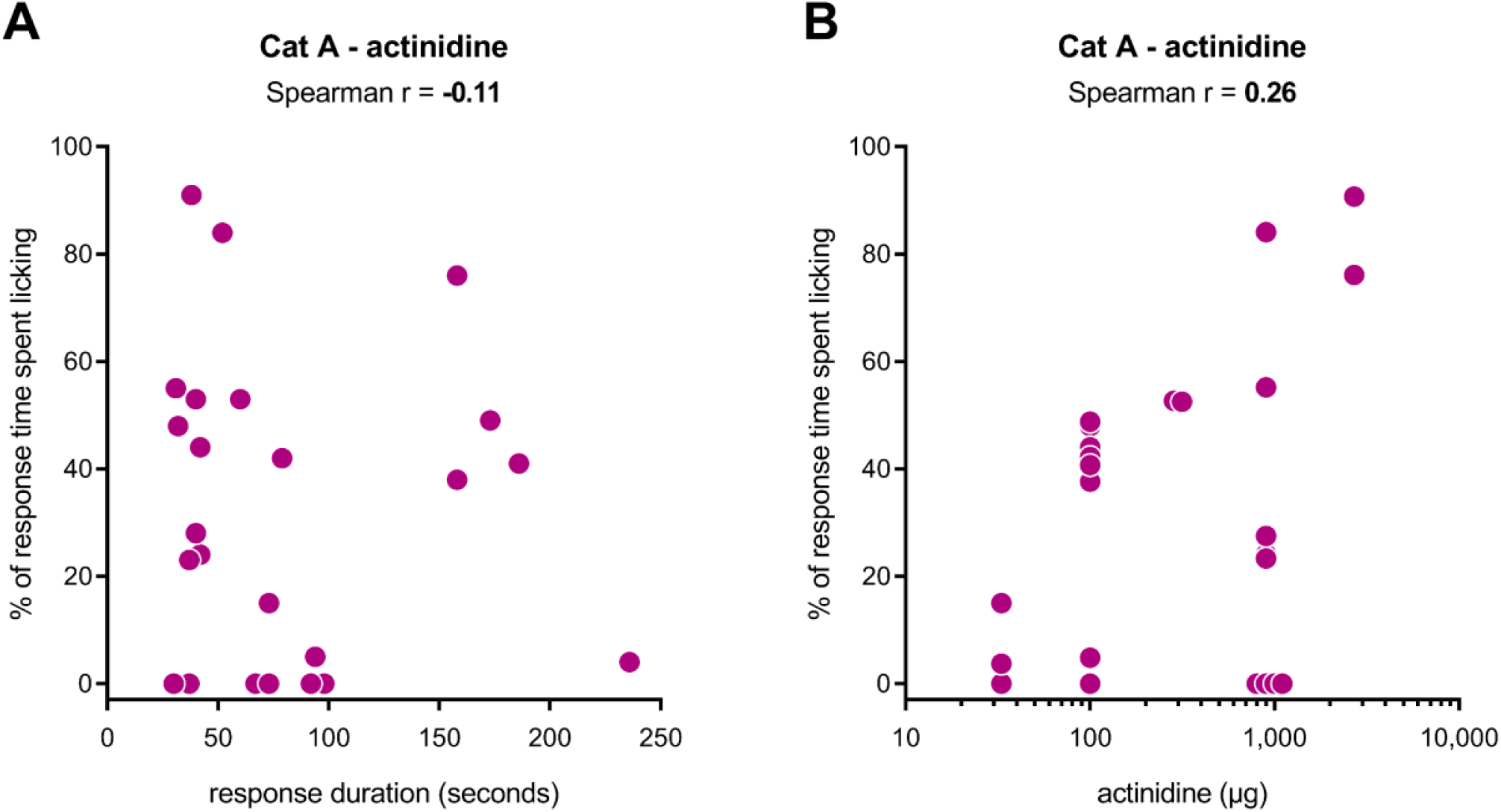
Absence of correlation between time spent licking and response duration (**A**) and time spent licking and actinidine quantity (33, 100, 300, 900 and 2700 µg) (**B**). Data from 24 responses longer than 30 seconds in duration to actinidine of cat A are shown. Some data points were overlapping and were edited for visualization purposes only. Actinidine quantity (µg) is shown on a log10 scale.

**Supplementary Figure 11.**
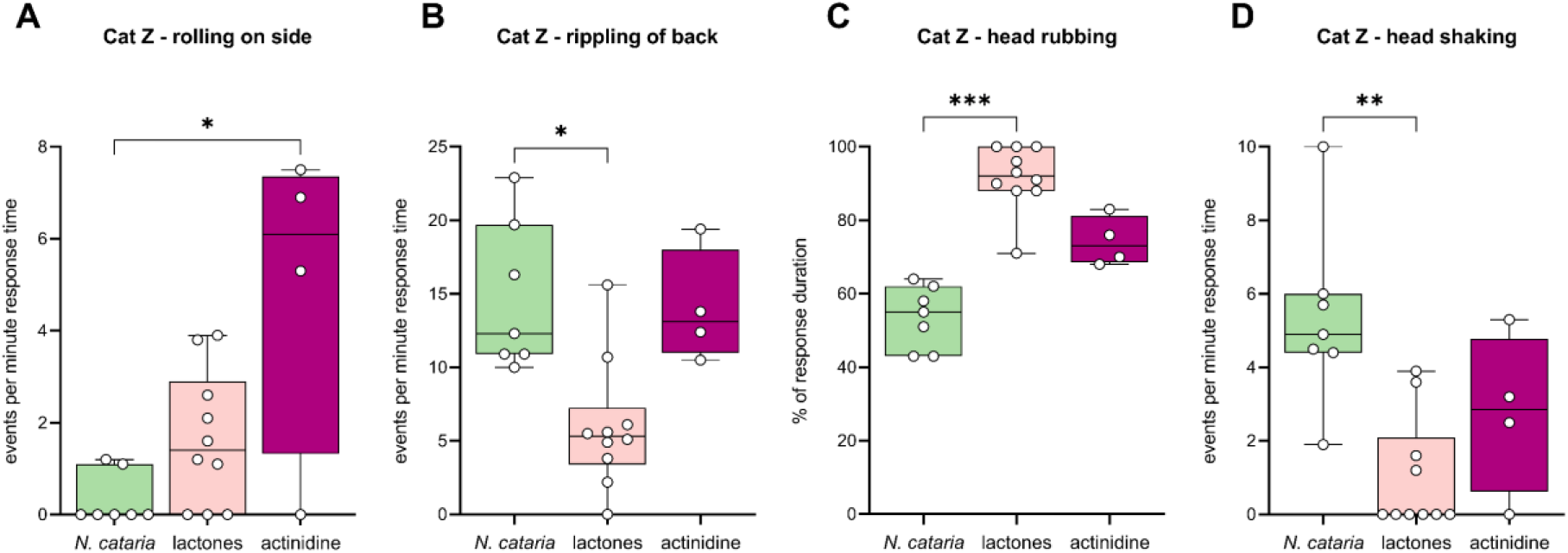
Differences in behavior of cat Z between responses to *N. cataria*, lactones and actinidine.

**Supplementary Figure 12.**
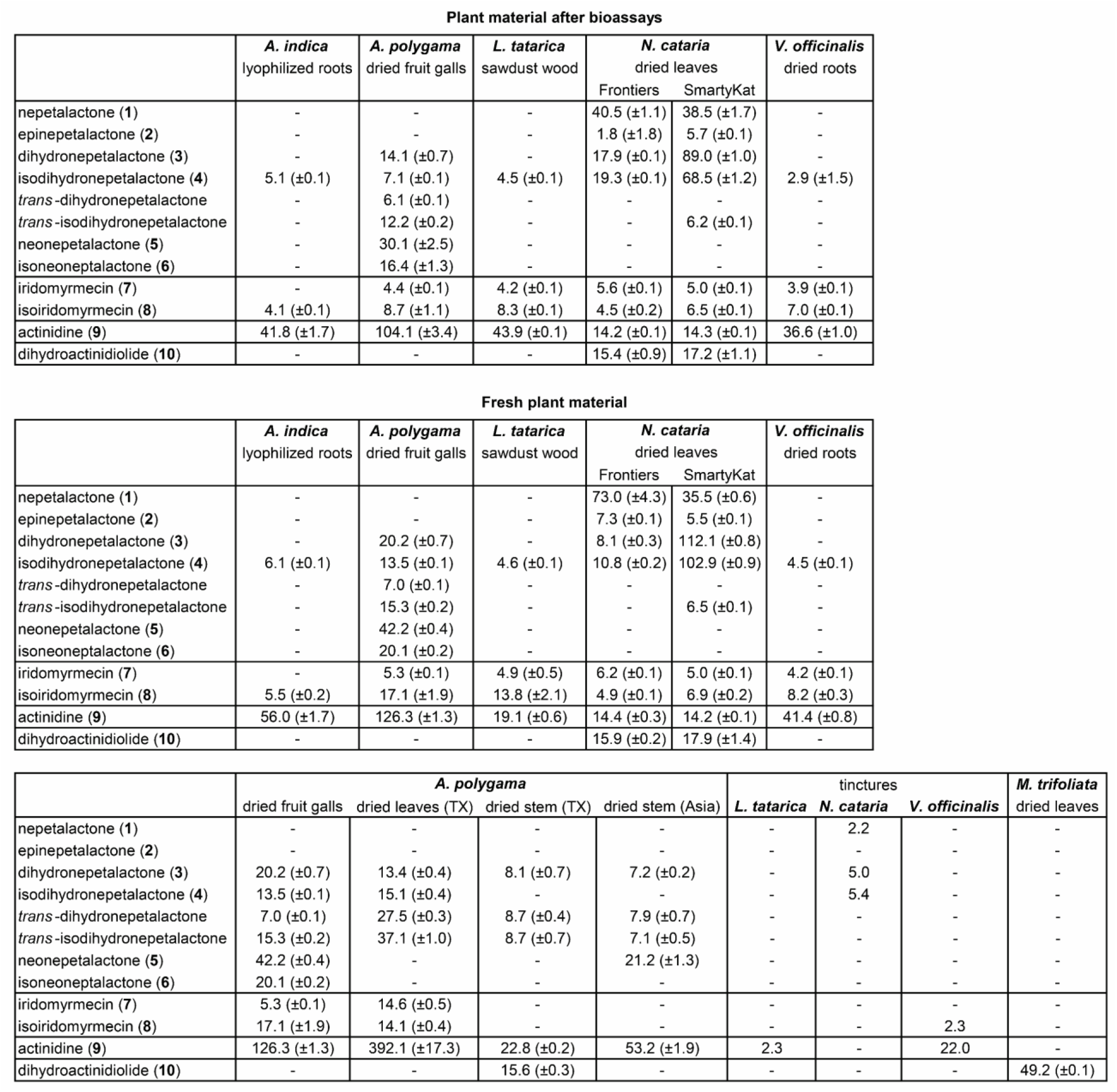
(**top**) Quantitation of cat-attracting compounds in the plant material taken from the socks after testing 10 × 10 hours in a 5-week testing period. Amounts are reported as µg per gram plant material used in this study. Reported values are the average of three separate extractions of the plant material. The standard error of the mean is reported between parentheses. Dashes indicate that the compound was not detected. (**middle and bottom**) Data from Figure 20 reported with the standard error of the mean and unrounded numbers.

**Supplementary Figure 13.**
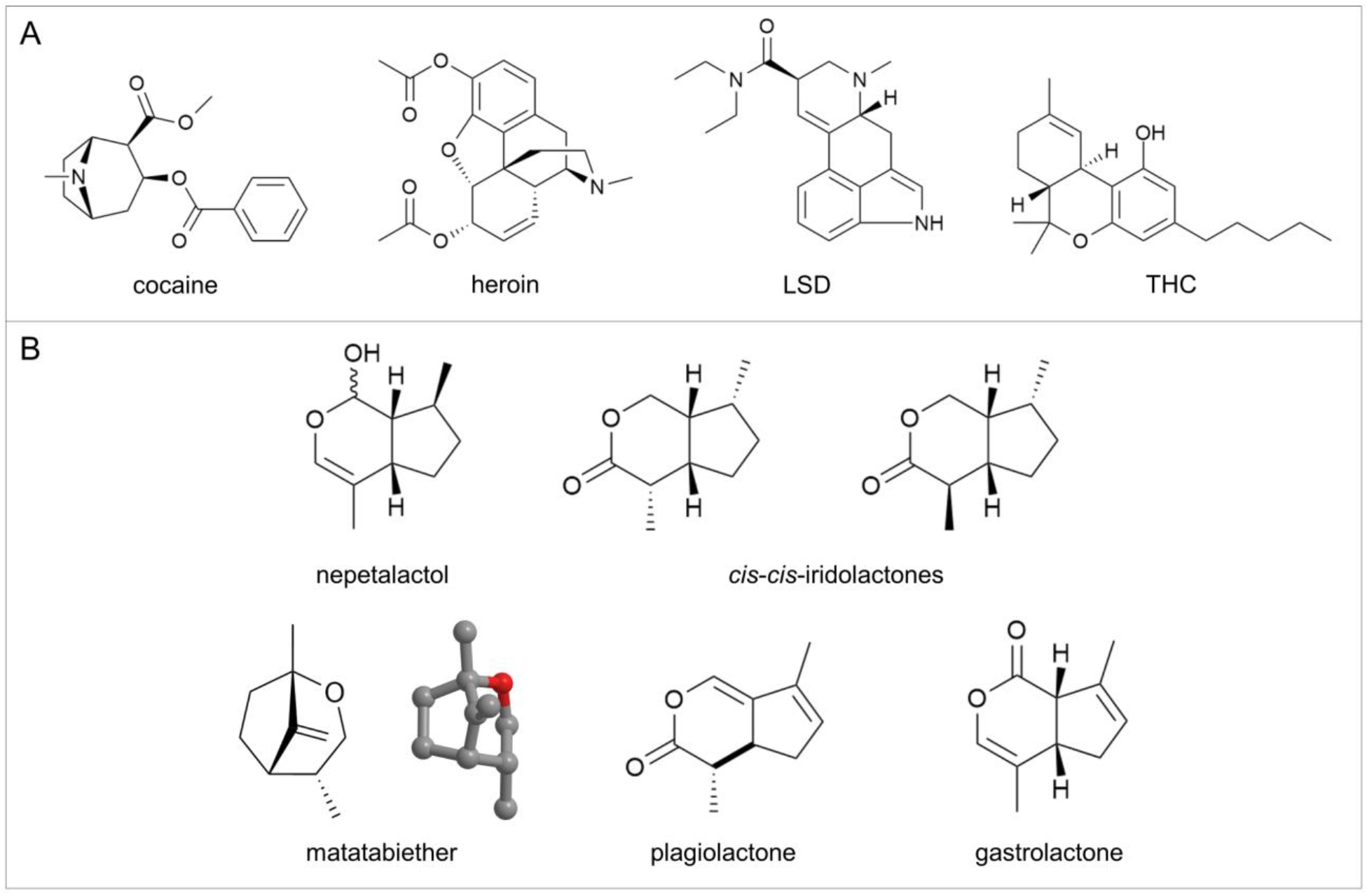
(**A**) The structures of cocaine, heroin, LSD and THC do not resemble the structure of the cat-attracting single compounds (Figure 12). (**B**) The structures of nepetalactol, both *cis*- *cis*-iridolactones, plagiolactone, gastrolactone, and both the 2D and 3D structure of the bridged bicyclic matatabiether.

**Supplementary Figure 14.**
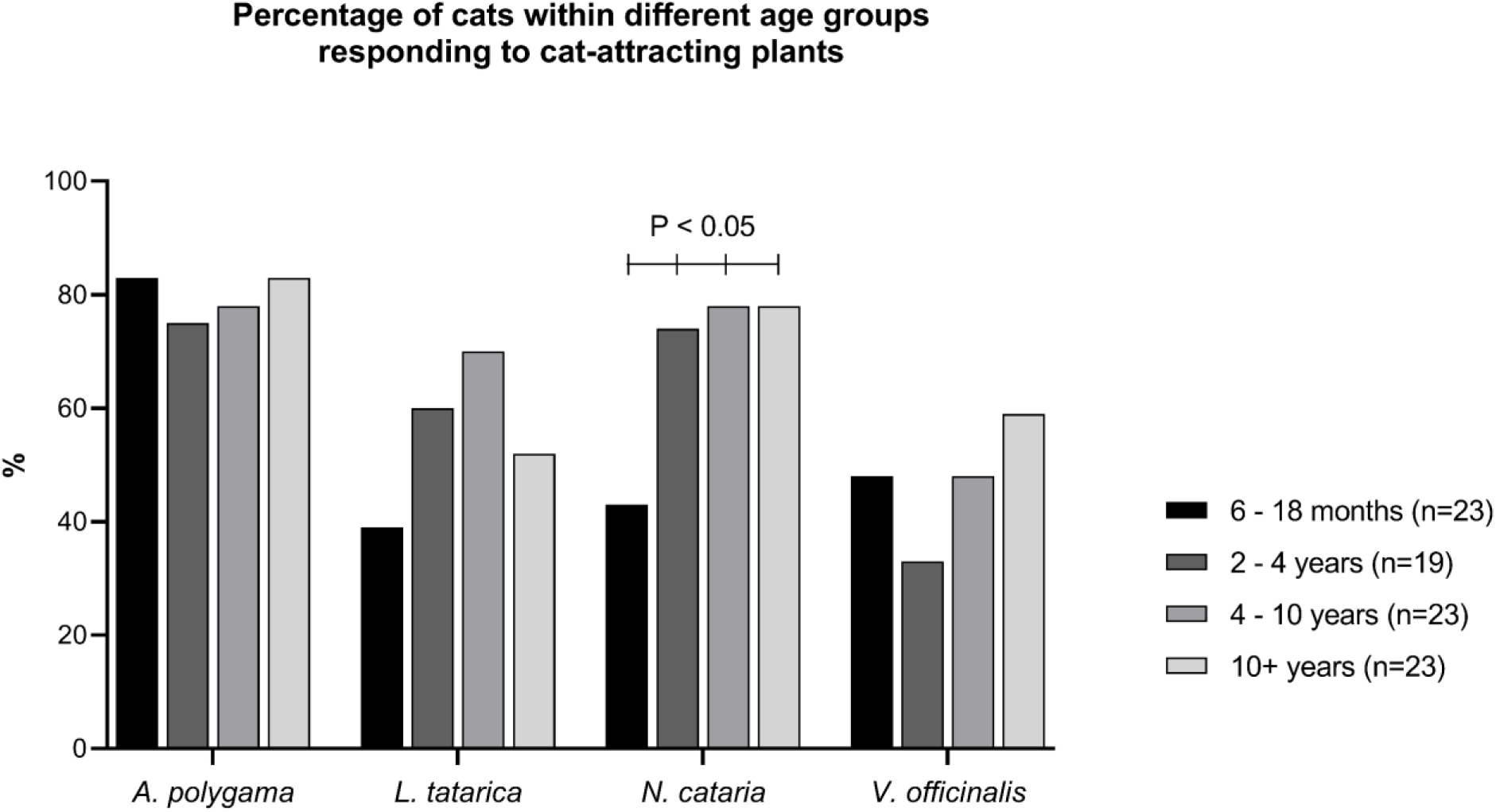
The fraction of domestic cats, within 4 age groups, that responded to a variety of cat-attracting plants. Significantly fewer of the younger cats (6 – 18 months) showed a response to catnip when compared to the 3 older groups (43% versus 74, 78 and 78%; P < 0.05, Fisher’s exact test). There were no cats 18 – 24 months of age.

## Supplementary files

**Supplementary File 1**. Video showing behavior of domestic cats seen in response to cat-attracting plants and single compounds (**Table 4**). Four recordings are shown for each behavior. (MP4)

**Supplementary File 2**. DNA barcoding sequence of *cox1* from *Pseudasphondylia matatabi*, and sequences of *matK*, *rbcL,* and *psbA* – *trnH* from *Actinidia* and *Lonicera* species. (TXT)

**Supplementary File 3**. Raw data. This file contains the raw data from the plant, single compound, and habituation / dishabituation experiments, as well as the BORIS analyses (plants and single compounds). (XLSX)

**Supplementary File 4**. Video of cats A, H, N, O, V and Z responding to catnip. One response is shown for each cat. (MP4)

## Notes

https://figshare.com/articles/media/Video_ethogram_showing_behavior_of_domestic_cats_in_response_to_cat-attracting_plants_/19312481

https://figshare.com/articles/media/Six_domestic_cats_responding_to_catnip/19312484

### Competing Interest Statement

The authors have declared no competing interest.

## References

Albone, E. S. (1975) Dihydroactinidiolide in the supracaudal scent gland secretion of the red fox. Nature, 256, 575.

Beckett, J. S., J. D. Beckett & J. E. Hofferberth (2010) A divergent approach to the diastereoselective synthesis of several ant-associated iridoids. Org Lett, 12, 1408–11.

Beran, F., T. G. Köllner, J. Gershenzon & D. Tholl (2019) Chemical convergence between plants and insects: biosynthetic origins and functions of common secondary metabolites. New Phytol, 223, 52–67.

Bicchi, C., M. Mashaly & P. Sandra (1984) Constituents of Essential Oil of Nepeta nepetella. Planta Med, 50, 96–8.

Blum, M. S., J. B. Wallace, R. M. Duffield, J. M. Brand, H. M. Fales & E. A. Sokoloski (1978) Chrysomelidial in the defensive secretion of the leaf beetle *Gastrophysa cyanea* Melsheimer. Journal of Chemical Ecology, 4, 47–53.

Bol, S., J. Caspers, L. Buckingham, G. D. Anderson-Shelton, C. Ridgway, C. A. Buffington, S. Schulz & E. M. Bunnik (2017) Responsiveness of cats (*Felidae*) to silver vine (*Actinidia polygama*), Tatarian honeysuckle (*Lonicera tatarica*), valerian (*Valeriana officinalis*) and catnip (*Nepeta cataria*). BMC Vet Res, 13, 70.

Ebrahim, S. A., H. K. Dweck, J. Stökl, J. E. Hofferberth, F. Trona, K. Weniger, J. Rybak, Y. Seki, M. C. Stensmyr, S. Sachse, B. S. Hansson & M. Knaden (2015) Drosophila Avoids Parasitoids by Sensing Their Semiochemicals via a Dedicated Olfactory Circuit. PLoS Biol, 13, e1002318.

Eisenbraun, E. J., C. E. Browne, R. L. Irvin-Willis, D. J. McGurk, E. L. Eliel & D. L. Harris (1980) Structure and Stereochemistry of 4aβ,7α,7aβ-NepetaIactone from Nepeta mussini and Its Relationship to the 4aα,7α,7aα- and 4aα,7α,7aβ-Nepetalactones from N. cataria. The Journal of Organic Chemistry, 45, 3811–3814.

Eisner, T. (1964) Catnip: Its Raison d’Être. Science, 146, 1318–20.

Endallew, S. A. & E. Dagne (2020) Isolation, Characterization and Quantification of Civetone from Civet Musk. Chemical Sciences Journal, 11.

Enders, D. & A. Kaiser (1997) A Short Asymmetric Synthesis of (-)-Neonepetalactone. Heterocycles, 46, 631–5.

Formisano, C., D. Rigano & F. Senatore (2011) Chemical constituents and biological activities of *Nepeta* species. Chem Biodivers, 8, 1783–818.

Friard, O. & M. Gamba (2016) BORIS: a free, versatile open-source event-logging software for video/audio coding and live observations. Methods in Ecology and Evolution, 7, 1325–1330.

Hill, J. O., E. J. Pavlik, G. L. Smith III, G. M. Burghardt & P. B. Coulson (1976) Species-characteristic responses to catnip by undomesticated felids. Journal of Chemical Ecology, 2, 239–53.

Ho, H. Y. & Y. S. Chow (1993) Chemical identification of defensive secretion of stick insect, *Megacrania tsudai* Shiraki. J Chem Ecol, 19, 39–46.

Johnson, R. D. & G. R. Waller (1971) Isolation of actinidine from *Valeriana officinalis*. Phytochemistry, 10, 3334–5.

Kanehisa, K., H. Tsumuki & K. Kawazu (1994) Actinidine secreting rove beetles (*Coleoptera: staphylinidae*). Applied Entomology and Zoology, 29, 245–251.

Katahira, K. & E. Iwai (1975) Effect of unilateral lesion of amygdala on unmanifested response to Matatabi (*Actinidia polygama*) in cats. Tohoku J Exp Med, 115, 137–43.

Keesey, I. W., N. Jiang, J. Weißflog, R. Winz, A. Svatoš, C. Z. Wang, B. S. Hansson & M. Knaden (2019) Plant-Based Natural Product Chemistry for Integrated Pest Management of *Drosophila suzukii*. J Chem Ecol, 45, 626–637.

McElvain, S. M., R. D. Bright & P. R. Johnson (1941) The Constituents of the Volatile Oil of Catnip. I. Nepetalic Acid, Nepetalactone and Related Compounds. Journal of the American Chemical Society, 63, 1558–63.

McLean, S., N. W. Davies & D. S. Nichols (2017) Lipids of the Tail Gland, Body and Muzzle Fur of the Red Fox, *Vulpes vulpes*. Lipids, 52, 599–617.

McLean, S., D. S. Nichols & N. W. Davies (2021) Volatile scent chemicals in the urine of the red fox, *Vulpes vulpes*. PLoS One, 16, e0248961.

Meinwald, J., T. H. Jones, T. Eisner & K. Hicks (1977) New methylcyclopentanoid terpenes from the larval defensive secretion of a chrysomelid beetle (*Plagiodera versicolora*). Proc Natl Acad Sci U S A, 74, 2189–93.

Melo, N., M. Capek, O. M. Arenas, A. Afify, A. Yilmaz, C. J. Potter, P. J. Laminette, A. Para, M. Gallio & M. C. Stensmyr (2021) The irritant receptor TRPA1 mediates the mosquito repellent effect of catnip. Curr Biol, 31, 1988–1994.e5.

Miyazaki, M., T. Miyazaki, T. Nishimura, W. Hojo & T. Yamashita (2018) The Chemical Basis of Species, Sex, and Individual Recognition Using Feces in the Domestic Cat. J Chem Ecol, 44, 364–373.

Nangia, A., G. Prasuna & P. Bheema Rao (1997) Synthesis of cyclopenta[c]pyran skeleton. Tetrahedron, 53, 14507–14545.

Nelson, D. L. 1968. Methylcyclopentane Monoterpenes of Nepeta Cataria and Actinidia Polygama (thesis). Purdue University, West Lafayette, IN, USA.

Notredame, C., D. G. Higgins & J. Heringa (2000) T-Coffee: A novel method for fast and accurate multiple sequence alignment. J Mol Biol, 302, 205–17.

Palen, G. F. & G. V. Goddard (1966) Catnip and oestrous behaviour in the cat. Anim Behav, 14, 372–7.

Ray, J. 1660. Catalogus plantarum circa Cantabrigiam nascentium. Cantabrigia.

Regnier, F. E., G. R. Waller & E. J. Eisenbraun (1967) Studies on the composition of the essential oils of three *Nepeta* species. Phytochemistry, 6, 1281–1289.

Sakan, T., A. Fujino & F. Murai (1960) Chemical components in matatabi. I. The isolation of the physiologically active principles, matatabilactone and actinidine. Journal of the Chemical Society of Japan. Pure chemistry section, 81, 1320–4.

Sakan, T., A. Fujino, F. Murai, Y. Butsugan & A. Suzui (1959) On the structure actinidine and matatabilactone, the effect components of *Actinidia polygama*. Bulletin of the Chemical Society of Japan, 32, 315–6.

Sakan, T., S. Isoe & S. B. Hyeon (1967) The structure of actinidiolide, dihydroactinidiolide and actinidol. Tetrahedron Letters, 17, 1623–1627.

Sakan, T., S. Isoe, S. B. Hyeon, R. Katsumura, T. Maeda, J. Wolinsky, D. Dickerson, M. Slabaugh & D. Nelson (1965) The exact nature of matatabilactone and the terpenes of *Nepeta cataria*. Tetrahedron Letters, 6, 4097–102.

Sakan, T., F. Murai, S. Isoe, S. B. Hyeon & Y. Hayashi (1969) The Biologically Active C9-, C10-, and C11-Terpenes from A*ctinidia polygama* Miq., *Boschniakia rossica* Hult, and *Menyanthes trifoliata* L. Nippon kagaku zassi, 90, 507–528.

Salmon, W. 1710. Botanologia. The English Herbal: or, History of Plants. London.

Scaffidi, A., D. Algar, B. Bohman, E. L. Ghisalberti & G. Flematti (2016) Identification of the Cat Attractants Isodihydronepetalactone and Isoiridomyrmecin from *Acalypha indica*. Australian Journal of Chemistry, 69, 169–73.

Starkenmann, C., Y. Niclass, I. Cayeux, R. Brauchli & A.-C. Gagnon (2015) Odorant volatile sulfur compounds in cat urine: occurrence of (+/-)-3,7-dimethyloct-3-sulfanyl-6-en-1-ol and its cysteine conjugate precursor. Flavour and Fragrance Journal, 30, 91–100.

Stepanov, A. V. & V. V. Veselovsky (1997) Stereocontrolled synthesis of (+)- and (−)-iridomyrmecin from citronellene enantiomers. Russian Chemical Bulletin, 46, 1606–10.

Stökl, J., J. Hofferberth, M. Pritschet, M. Brummer & J. Ruther (2012) Stereoselective chemical defense in the *Drosophila* parasitoid *Leptopilina heterotoma* is mediated by (-)-iridomyrmecin and (+)- isoiridomyrmecin. J Chem Ecol, 38, 331–9.

Sun, Z., T. Gao, H. Yao, L. Shi, Y. Zhu & S. Chen (2011) Identification of *Lonicera japonica* and its related species using the DNA barcoding method. Planta Med, 77, 301–6.

Thomas, P., G. Balme, L. Hunter & J. McCabe-Parodi (2005) Using Scent Attractants to Non-Invasively Collect Hair Samples from Cheetahs, Leopards and Lions. *Animal Keepers’* Forum, 32, 342–348.

Todd, N. B. (1962) Inheritance of the catnip response in domestic cats. J Hered, 53, 54–6.

Todd, N. B. 1963. The Catnip Response (thesis). Harvard University, Cambridge, MA, USA.

Tucker, A. O. & S. S. Tucker (1988) Catnip and the catnip response. Economic Botany, 42, 214–31.

Uenoyama, R., T. Miyazaki, J. L. Hurst, R. J. Beynon, M. Adachi, T. Murooka, I. Onoda, Y. Miyazawa, R. Katayama, T. Yamashita, S. Kaneko, T. Nishikawa & M. Miyazaki (2021) The characteristic response of domestic cats to plant iridoids allows them to gain chemical defense against mosquitoes. Sci Adv, 7.

Uetake, K., T. Abumi, T. Suzuki, S. Hisamatsu & M. Fukuda (2018) Volatile faecal components related to sex and age in domestic cats (*Felis catus*). Journal of Applied Animal Research, 46, 766–70.

Waller, G. R., G. H. Price & E. D. Mitchell (1969) Feline attractant, *cis*,*trans*-nepetalactone: metabolism in the domestic cat. Science, 164, 1281–2.

Weihong, B., L. Dawei & L. Xinwei (2018) DNA barcoding of *Actinidia* (*Actinidiaceae*) using internal transcribed spacer, matK, rbcL and trnH-psbA, and its taxonomic implication. New Zealand Journal of Botany, 56, 360–371.

Welzel, K. F., S. H. Lee, A. T. Dossey, K. R. Chauhan & D. H. Choe (2018) Verification of Argentine ant defensive compounds and their behavioral effects on heterospecific competitors and conspecific nestmates. Sci Rep, 8, 1477.

Wolinsky, J., T. Gibson, D. Chan & H. Wolf (1965) Stereospecific syntheses of iridomyrmecin and related iridolactones. Tetrahedron, 21, 1247–61.

